# EBF1 controls an embryonic artery-forming niche that reactivates in pulmonary arterial hypertension

**DOI:** 10.1101/2025.05.02.651303

**Authors:** Wen Tian, Timothy Ting-Hsuan Wu, Shenbiao Gu, Seunghee Lee, Hanqiu Zhao, Adam M. Andruska, Kyle K. Song, Jason L. Chang, Cerianne Huang, Ryan Vinh, Dongeon Kim, Yu Zhu, Evan Bao, Stuti Agarwal, Dan Yi, Aiqin Cao, Junliang Pan, Peter N. Kao, Tushar Desai, Roham Zamanian, Ke Yuan, Lawrence S. Prince, Lindsay D. Butcher, Vinicio A. de Jesus Perez, Roger A. Johns, Xinguo Jiang, Patrick E. Bogard, Jason Hong, Marlene Rabinovitch, Kristy Red-Horse, Mark A. Krasnow, Zhiyu Dai, Mark R. Nicolls

## Abstract

Developmental programs that orchestrate cell fate and tissue architecture during organogenesis can cause disease when reactivated in adults. Here we identify a population of endothelial cells (ECs) defined by the pioneer factor early B cell factor 1 (EBF1) that controls both pulmonary artery (PA) morphogenesis and pulmonary arterial hypertension (PAH). During embryonic development, the PA emerges from an endothelial niche harbored within the vascular plexus, where *Aplnr*^+^ endothelial progenitors give rise to both arterial ECs and EBF1*^+^* ECs. Rather than incorporating directly into the PA intima, EBF1^+^ ECs control the branching and maturation of the PA tree through paracrine vasculotrophic signals that direct plexus expansion, arterialization, and mural cell recruitment. Although essential for development, most EBF1^+^ ECs disappear upon completion of PA morphogenesis, leaving only a rare residual population within the adult *Aplnr^+^* gCap compartment. In PAH, vascular injury reactivates the EBF1 program through recruitment of the pre-existing *Ebf1* lineage together with an additional contribution consistent with *de novo Ebf1* induction in gCaps, reconstituting this artery-forming niche in a maladaptive reprise of PA development. Unlike their transient embryonic counterparts, these EBF1^+^ ECs persist within occlusive intimal lesions and express vasculotrophic signals associated with pathological PA remodeling. Combined with endothelial injury, gCap-specific *Ebf1* overexpression drives a coordinated program of neo-arterialization, occlusive vascular remodeling, and severe PAH. Conversely, gCap-specific genetic deletion of *Ebf1* in mice and endothelial-targeted AAV-mediated *Ebf1* knockdown in rats protect against disease, showing that endothelial EBF1 is required for PAH pathogenesis. These findings demonstrate that reactivation and persistence of a transient embryonic artery-forming niche in adulthood drive pathological vascular remodeling and PAH.

## Main

The pulmonary arteries (PA) of the mammalian lung must balance two seemingly opposing physiological demands: transporting the entire cardiac output for gas exchange while maintaining low resistance and pressure to protect the fragile alveolar capillaries^1^. To achieve the required high vascular capacitance, the PA rapidly bifurcates into right and left main arteries, which further branch into small-caliber arteries and highly distensible precapillary arterioles, each composed of a single layer of endothelium (the intima) ensheathed by smooth muscle cells and an outer adventitial layer^2^. While the most proximal segments of the pulmonary trunk are derived from cardiac progenitors^3,4^, the cellular origins and signal sources that control the expansion, branching, and maturation of the distal PA remain incompletely understood.

The delicate structure of distal PAs also renders them highly susceptible to injury-induced pathology, most notably in pulmonary hypertension (PH), a disease afflicting nearly 80 million people worldwide^5^. Pulmonary arterial hypertension (PAH) is the most lethal and progressive form of PH, characterized by occlusive vascular remodeling of the distal PAs with neointimal expansion, smooth muscle hypertrophy, and adventitial fibrosis^5^. Together, these vascular changes obstruct blood flow, elevate pulmonary vascular resistance, and, ultimately, cause right heart failure and death^6,7^. Despite decades of progress in understanding PAH disease mechanisms^8-12^, including the landmark discovery of pathogenic *BMPR2* mutations^13^, the disease remains incurable, with nearly 40% of patients dying within five years of diagnosis. Advancing therapies to address the underlying vascular pathology requires a better understanding of the cellular origins, gene programs, and molecular signals that initiate, sustain, and propagate PA remodeling.

Here, we draw insight from PA development to better understand PAH pathogenesis. Integrating single-cell RNA sequencing (scRNA-seq), lineage tracing, and genetic perturbation experiments, we show that distal PA morphogenesis is orchestrated by a specialized embryonic endothelial niche defined by EBF1^+^ ECs within the developing pulmonary vascular plexus. Through coordinated expression of paracrine vasculotrophic signals, these EBF1^+^ ECs organize their neighboring plexus and stromal progenitors to establish the layered architecture of the PA. Although critical in development, most of these cells disappear upon completion of PA morphogenesis, leaving only a rare residual population in the adult gCap compartment. This embryonic artery-forming niche is reconstituted in adult PAH through reactivation of the EBF1 program within the adult gCap compartment, giving rise to persistent EBF1^+^ ECs that coordinate neo-arterialization and occlusive vascular remodeling in a maladaptive reprise of development. Our findings establish EBF1 as the central regulator of a transient embryonic artery-forming niche that orchestrates PA morphogenesis and, when pathologically reconstituted in adulthood, drives pulmonary vascular remodeling in PAH.

### PAH neointima features a novel population of EBF1-expressing endothelial cells

To investigate the nature of the pathological vessel-occluding cells that contribute to the neointima in PAH, we performed single-cell RNA sequencing (scRNA-seq) in our previously described rat model of genetic PAH (**Extended Data Fig. 1a)**^14^. In this model, lung inflammation elicits severe vascular remodeling and neointimal formation selectively in animals harboring heterozygous mutations in *Bmpr2* (encoding bone morphogenetic protein receptor type 2, the gene most commonly mutated in heritable PAH^15,16^). Using protocols adapted from mouse and human lung atlases^17-19^, we processed rat lungs from three experimental groups (wildtype (WT), heterozygous silent carriers, and PAH) and profiled vascular endothelial (CD45^-^CD31^+^) and stromal (CD45^-^CD31^-^CD326^-^) cells (**Fig. 1a**). Markers conserved between human and mouse^19^ were used to identify the homologous molecular cell types of the rat pulmonary vasculature. A total of 23,075 cells were captured across all three conditions (11,648 cells from WT, 6,560 from silent carriers, and 4,867 from PAH; **Supplementary Table 5**). We identified all major canonical EC and stromal subtypes, including arterial, venous, general capillary (gCap)^20^, capillary aerocyte (aCap)^20^, lymphatic (Lymph), airway/vascular smooth muscle cells (ASM/VSM), pericytes (Peri), adventitial/alveolar fibroblasts (AdvF/AlvF), and proliferative fibroblasts (Fib-p) (**Fig. 1b, c**, **Extended Data Fig. 1b-d**, and **Supplementary Tables 1**, **2**).

**Fig. 1.**
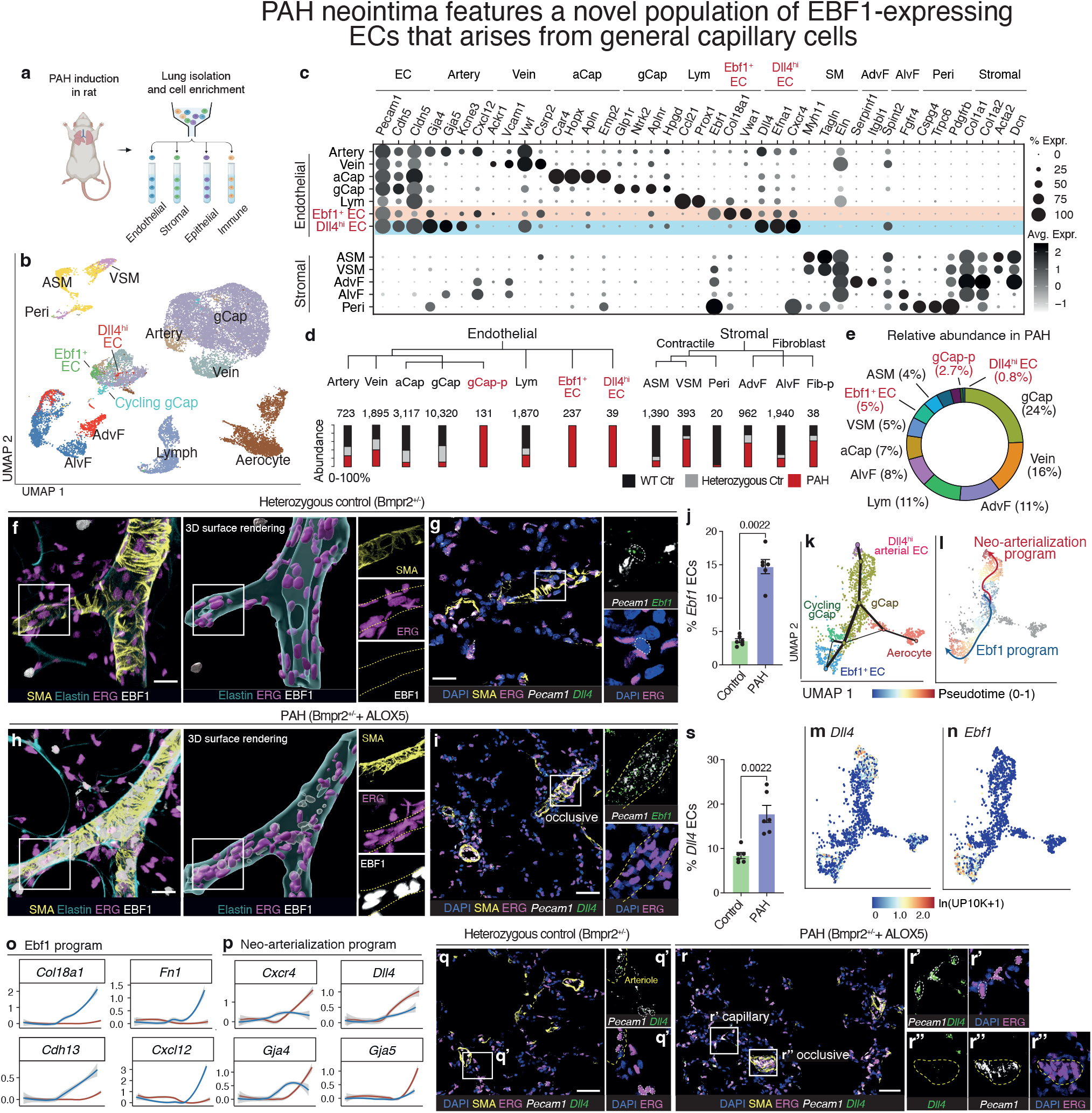
PAH neointima features a novel population of EBF1-expressing ECs that arises from general capillary cells. **a,** Single cell RNA-sequencing (scRNA-seq) workflow. Lungs were harvested from wildtype, Bmpr2^+/-^ heterozygous control, or PAH (Bmpr2^+/-^ + Ad*ALOX5*) rats (n=3 rats per group)^14^, dissociated into single cells, and FACS purified for endothelial (Epcam^-^ CD31^+^CD45^-^), stromal (Epcam^-^CD31^-^CD45^-^), epithelial (Epcam^+^CD31^-^CD45^-^), and immune (Epcam^-^CD31^-^CD45^+^) cells; endothelial and stromal fractions were profiled by scRNA-seq. **b,** Uniform Manifold Approximation and Projection (UMAP) visualization of annotated molecular clusters for endothelial and stromal cells from all conditions. Abbreviations: Art, artery; aCap, capillary aerocyte; gCap, general capillary; EC, endothelial cell; Lym, lymphatic endothelial; ASM, airway smooth muscle; VSM, vascular smooth muscle; Peri, pericyte; AdvF, adventitial fibroblast; AlvF, alveolar fibroblast. **c,** Dot plot (size indicates fraction expressing and shade indicates mean expression) of representative marker genes in annotated cell types. Names of PAH-enriched cell types are highlighted in red. **d,** Relative abundance of cell types in each condition. Numbers indicate total cell counts per type; stacked bars show relative abundance from each condition. Canonical cell types are shown in black; PAH-exclusive types (100% detected in PAH) are highlighted in red. Proliferative subsets are denoted by suffix “-p”. **e**, Relative abundance of all cell types detected in PAH, with PAH-exclusive types highlighted in red. **f-i,** Representative images of lungs from heterozygous control (**f**, **g**) and PAH (**h**, **i**) rats, assessed by whole-mount immunofluorescence and 3D surface rendering (**f**, **h**) or smFISH and immunofluorescence (**g**, **i**). Yellow dashed outlines trace the SMA^+^ vessel wall; white dashed outlines mark individual cells throughout **f-i. f**, **h**, n=6 rats per group. **g**, Inset shows a rare *Ebf1*^+^ERG^-^ stromal cell. **i**, Yellow outline marks an occlusive pulmonary arteriole; insets show *Ebf1*^+^ *Pecam1*^+^ERG^+^ ECs within the lesion. n=6 rats per group. **j,** Quantification of *Ebf1*^+^ ECs (% of ERG^+^ nuclei that have *Ebf1* puncta) within vascular ROIs (pulmonary arterioles [PA] <50 μm in diameter). n=6 rats per group. **k,** UMAP plot overlaid by lineage graph of neointimal ECs (see **Extended Data** Fig. 1) in rat PAH lungs with single cells colored by cell type, lineage graph nodes are sub-cluster centroids with size scaled by number of cells, and edges are scaled by interaction strength between clusters (summed SNN-graph edges; see Methods). **l,** Pseudotime trajectory of bifurcating differentiation from gCap ECs toward *Dll4* arterial ECs (red, “neo-arterialization program”) or *Ebf1* ECs (blue, “Ebf1 program”). Principal curves were computed by Slingshot; cells are colored by pseudotime (0-1). **m, n,** UMAP plots showing marker gene expression (ln [UMIs per 10,000 + 1]). **o, p,** Loess-smoothed pseudotime expression profiles for lineage-associated genes in each program. Significant pseudotime-associated genes were identified by tradeSeq (Full DEG list in **Supplementary Table 3**). **q, r**, Representative smFISH and immunofluorescence images from heterozygous control (**q**) and PAH (**r**) rats visualizing *Dll4* RNA with endothelial markers ERG and *Pecam1*. **q**, Inset (**q’**, white dashed outlines) shows *Dll4*^+^ arterial ECs (*Dll4*^+^*Pecam1*^+^ERG^+^) within the arteriolar lumen (yellow dashed outline). **r**, Insets show *Dll4*^+^ arterial ECs within a distal capillary (**r’**, dashed outlines) and within an occlusive lesion (**r’’**; dashed outline traces the thickened, SMA^+^ vessel wall of the occlusive lesion). n=6 rats per group. **s**, Quantification of *Dll4^+^* arterial ECs (% of ERG^+^ nuclei with *Dll4* puncta) within ROIs. n=6 rats per group. For **j** and **s**, 5-8 vascular ROIs (PA caliber <50 μm) per rat were analyzed and averaged to generate a single biological replicate value. Data are presented as mean ± s.e.m. with individual data points shown. Statistical comparisons in **j** and **s** were performed using two-sided Mann-Whitney tests; p values are indicated on the graphs. Scale bars, 20 μm in **f**, **h** and 25 μm in **g**, **i**, **q**, and **r**.

Although most cell types were recovered across all groups, just three populations – all of which were endothelial – showed marked enrichment in disease, including a proliferative state of gCap cells (gCap-p) and two previously unreported EC clusters (**Fig. 1d**, **Extended Data Fig. 1e-g**, and **Supplementary Table 2**). One minor cluster (0.8% of all PAH cells, referred to as *Dll4*^hi^ arterial ECs) expressed high levels of the Notch ligand *Dll4* alongside arterial markers (*Gja5*, *Cxcr4*)^21^. The other, major cluster (4.9% of PAH cells, referred to as *Ebf1^+^* ECs), did not resemble any known EC subtype and instead expressed mesenchymal genes (*Col18a1*, *Vim*)^22,23^, the motility-regulating atypical T-cadherin (*Cdh13*)^24^, and *Ebf1*, which encodes a pioneer transcription factor^25-28^. Among the PAH-associated endothelial populations, *Ebf1*^+^ ECs were the most abundant and transcriptionally distinct from canonical ECs (**Fig. 1e** and **Extended Data Fig. 1h**). To localize these cells in PAH lungs, we performed immunostaining and single-molecule fluorescent *in situ* hybridization (smFISH). While rare in control PA endothelium, EBF1*^+^* ECs (*Pecam1*^+^*Ebf1*^+^ERG^+^) were prominent within occlusive intimal lesions in PAH lungs (**Fig. 1f-j**). To determine whether EBF1 ECs were also present within occlusive intimal lesions across diverse preclinical models of PAH, we analyzed two additional rat models (monocrotaline^29^ and athymic rat plus SU5416^30^) and two mouse models (lung-specific TNFα-overexpression^31^ and house-dust-mite-induced PAH^32^). In both rat and house-dust-mite-induced mouse models of PAH, EBF1 ECs were readily detected within neointimal lesions and regions of intimal thickening (**Extended Data Fig. 2a-c**). Although the TNFα-overexpression lungs exhibited relatively limited intimal expansion, EBF1^+^ ECs were still induced in the lumen of remodeled arterioles, indicating that induction of EBF1-expressing ECs is a shared feature across diverse preclinical etiologies (**Extended Data Fig. 2d**).

### Neo-arterialization and EBF1 programs in PAH arise from bifurcating differentiation of gCap stem cells

Our scRNA-seq dataset provided broad coverage of ECs yet resolved no distinct *Ebf1*^+^ EC population in either WT or silent-carrier rats. Instead, we identified a distinct, disease-specific proliferative state of *Aplnr*-expressing gCap cells (gCap-p), detected exclusively in PAH lungs. As gCap cells have been shown to function as capillary stem cells in disease^20,33,34^, we asked whether they could be the precursors to PAH neointimal cells. To test this hypothesis, we examined pairwise transitions among endothelial clusters in PAH lungs and reconstructed the transcriptional programs underlying this endothelial transformation. The resulting lineage graph revealed two novel fate trajectories with gCap cells at the root, in addition to their known differentiation into capillary aerocytes^20^. One trajectory connected gCap cells to the *Dll4*^hi^ arterial ECs, whereas the other led to *Ebf1*^+^ ECs (**Fig. 1k-n**), suggesting gCap cells as a common progenitor source for both PAH-enriched endothelial populations.

Differential gene expression analysis identified 120 differentially expressed genes (DEGs) as gCap cells differentiated into *Ebf1*^+^ ECs, defining an “Ebf1 program” (**Fig. 1o** and **Supplementary Table 3**). These transitioning cells retained core EC markers (*Pecam1*, *Cdh5*), while downregulating a key endothelial barrier component (*Cldn5*) and the gCap identity marker (*Aplnr*)^20^. The induced DEGs, alongside *Ebf1*, were involved in ECM synthesis and remodeling (*Col18a1*, *Fn1*)^35,36^, cell-matrix adhesion and migration (*Tm4sf1, S100a4, Fblim1, Pdlim4*)^37-42^, and angiogenesis (*Ptn, Jag2, Cxcl12*)^43-46^. This hybrid transcriptional state, retaining endothelial identity while acquiring migratory and matrix-remodeling capacity, is characteristic of *partial* endothelial-to-mesenchymal transition (pEndoMT^47-50^), an endothelial phenotype documented in human PAH lesions^14,51,52^.

In parallel, we defined a “neo-arterialization program” comprising 248 DEGs as gCap cells appeared to generate new arterial cells (**Fig. 1p** and **Supplementary Table 3**). Cells advancing along this path upregulated key arterial and pre-arterial markers (*Gja4*/CX37, *Gja5*/CX40^53^), *Cxcr4* (critical regulator of arterial morphogenesis^54^), and high levels of *Dll4* (a Notch ligand essential for arterial specification^55^). Visualizing these neo-arterial *Dll4*^hi^ ECs *in situ* revealed their marked expansion in PAH lungs compared with controls, particularly in the capillary regions (**Fig. 1q-s**). Together, trajectory inference and histological validation indicate that PA vascular remodeling involves aberrant activation of gCap cells and their bifurcating differentiation into neo-arterial cells and EBF1^+^ ECs.

### EBF1^+^ ECs are a rich endothelial source of developmental vasculotrophic signals in PAH

A global receptor-ligand analysis across all profiled endothelial and stromal cells in PAH lungs revealed EBF1^+^ ECs as a major endothelial source of vascular developmental signals^56^ (**Extended Data Fig. 3** and **Supplementary Table 4**), with predicted outputs to their gCap progenitors and sister *Dll4*^+^ arterial ECs. Specifically, EBF1^+^ ECs expressed *Apln* (Apelin), which could signal to *Aplnr*^+^ gCap cells to promote their proliferation and guide their migration^57^ into neointimal lesions. They also expressed *Cxcl12*, which could guide migration^54^ of capillary-derived *Cxcr4*^+^*Dll4*^hi^ arterial ECs during neo-arterialization. These cells also expressed Notch ligands (*Jag1*, *Jag2*), which could induce arterial specification^21,55^. Beyond endothelial-directed signals, EBF1^+^ ECs expressed pro-fibrotic TGF-β (*Tgfb1*, *Tgfb2*)^58-60^ and ECM remodeling genes, suggesting a role in recruiting and activating the surrounding mural and stromal compartments during lesion maturation. We term these activities of EBF1^+^ ECs “vasculotrophic” to reflect their broad signaling range across vascular layers.

### Conserved capillary-derived EBF1 and neo-arterialization programs in human PAH

To determine whether EBF1^+^ ECs are present in human disease, we analyzed lung sections from patients with WHO Group I pulmonary arterial hypertension, a primary disease of the pulmonary arteries, obtained through the Pulmonary Hypertension Breakthrough Initiative (PHBI). While EBF1^+^ ECs (*PECAM1*^+^*EBF1*^+^ERG^+^) were rarely detected in control donor lungs, they were prominently localized in both plexiform and neointimal lesions in PAH (**Fig. 2a-c** and **Supplementary Table 6**). Together with our preclinical findings, these data indicate that EBF1 ECs are a conserved feature of PAH neointimal pathology across etiologies.

**Fig. 2.**
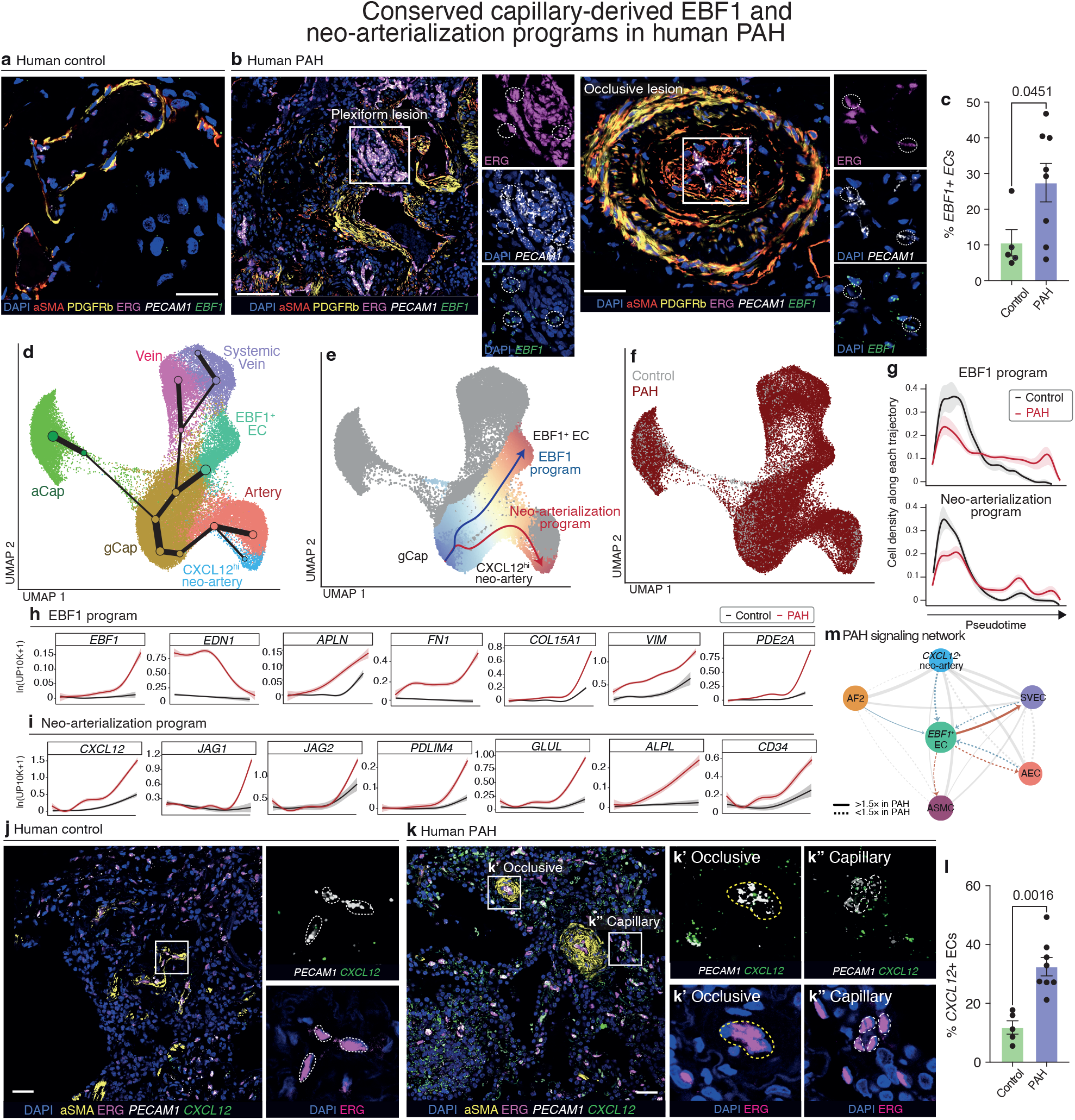
Conserved capillary-derived EBF1 and neo-arterialization programs in human PAH. **a, b,** Representative smFISH and immunofluorescence images of human lungs from control (**a**, n=5) and PAH patients (**b**, n=8). White dashed outlines mark cell or cell-cluster boundaries; yellow dashed outlines trace vessel or lesion walls, throughout **a**, **b**, **j**, and **k**. **b**, Insets show foci of *EBF1*^+^*PECAM1*^+^*ERG*^+^ ECs within a plexiform lesion and an occlusive lesion. **c**, Quantification of *EBF1^+^* ECs (% of ERG^+^ nuclei with *EBF1* puncta) within vascular regions of interest (ROIs). n=5 for controls and n=8 for PAH. **d**, UMAP plot overlaid by lineage graph of all ECs in human control and PAH lungs (see **Extended Data Fig. 4**) with single cells colored by annotated cell type. Lineage graph nodes are sub-cluster centroids with size scaled by number of cells, and edges are scaled by interaction strength between clusters (summed SNN-graph edges; see Methods). **e**, Pseudotime trajectory of bifurcating differentiation from gCap cells toward *CXCL12* neo-arterial ECs (red, “neo-arterialization program”) or *EBF1* ECs (blue, “EBF1 program”). Principal curves were computed by Slingshot; cells are colored by pseudotime (0-1). **f,** UMAP plot colored by condition. **g,** Cell density distributions along the EBF1 program and neo-arterialization program trajectories colored by condition. Lines represent mean kernel density estimates of pseudotime values across cells assigned to each lineage; shaded ribbons indicate 95% bootstrap confidence intervals computed by resampling patients with replacement (n = 200 iterations). Pseudotime is scaled 0–1. **h, i,** Loess-smoothed pseudotime expression profiles for lineage-associated genes. Significant pseudotime-associated genes were identified by tradeSeq (Full DEG list in **Supplementary Table 9**). **j**, **k**, Representative smFISH and immunofluorescence images of human lung sections from control (**j**, n=5) and PAH (**k**, n=8) patients. **j**, Inset shows individual *CXCL12*^+^ ECs (*CXCL12*^+^*PECAM1*^+^ERG^+^). **k**, Insets show *CXCL12^+^*neo-arterial *ECs* within an occlusive lesion (**k’**; yellow outline traces the lesion wall) and individual *CXCL12^+^* neo-arterial *ECs* within a distal capillary (**k’’**). **l**, Quantification of *CXCL12^+^* neo-arterial *ECs* (% of ERG^+^ nuclei with *CXCL12* puncta) within ROIs. n=5 for controls and n=8 for PAH. **m**, Ligand-receptor interactions inferred from human PAH scRNA-seq data between the indicated endothelial and stromal populations. Nodes represent cell states; arrows indicate predicted signaling direction (sender → receiver). Edges involving EBF1^+^ ECs are highlighted (red, signals sent by EBF1^+^ ECs; blue, signals received by EBF1^+^ ECs), with grey edges denoting interactions among other populations. Edge width is proportional to communication probability in PAH. Line style reflects relative interaction strength in PAH compared to control (solid, ≥1.5×; dashed, <1.5×). For **c** and **l**, 5-8 vascular ROIs (PA <100 μm in diameter) per donor were analyzed and averaged to generate a single biological replicate value; data are presented as mean ± s.e.m. with individual data points shown; statistical comparisons were performed using two-sided Mann-Whitney tests; p values are indicated on the graphs. Scale bars, 20 μm in **a**, **b**, **j**, and **k**.

To characterize the transcriptome of these EBF1^+^ ECs, we performed scRNA-seq analysis of 40 human lungs (30 PAH spanning idiopathic, congenital shunt-associated, and drug/toxin-induced etiologies; 10 control donors) from PHBI, capturing 317,414 high-quality transcriptomes (**Supplementary Tables 5**-**7**). After integration, we resolved all major vascular endothelial cell types in both pulmonary and systemic (bronchial) circulations^19^, including arterial ECs, venous ECs, general capillary ECs (gCaps/CAP1), aerocytes (aCaps/CAP2), and lymphatic ECs (LEC), alongside vascular smooth muscle cells and pericytes (**Extended Data Fig. 4** and **Supplementary Table 7**). Endothelial subtype composition was broadly comparable between PAH and controls (**Supplementary Table 7**).

Because *EBF1*^hi^ cells did not form a distinct EC cluster, we performed differential expression analysis across EC subtypes between PAH and control groups to identify cell populations with disease-dependent induction of *EBF1* (**Supplementary Table 8** and **Extended Data Fig. 4**). gCap emerged as the only EC subtype showing significant upregulation of *EBF1* across PAH subtypes (global test FDR = 0.016), with the largest effects in drug/toxin-induced disease (log_2_FC = 3.1, FDR = 0.0075; **Supplementary Table 8**). This capillary-restricted induction of *EBF1* mirrored our rat dataset and prompted trajectory analysis from gCap cells. The resulting lineage reconstruction revealed two PAH-associated trajectories originating from gCap cells (**Fig. 2d-f**). Although the overall composition of cells remained balanced between PAH and controls, PAH cells that entered these trajectories progressed further along pseudotime. The first was an *EBF1*^+^ gCap trajectory: the distribution of cells along this path differed significantly between PAH and controls (Kolmogorov-Smirnov test, p = 0.007), with the most advanced PAH cells showing particularly robust progression to late states (90^th^ percentile pseudotime Wilcoxon test, p = 0.001, **Fig. 2g**). The second was a *CXCL12*^hi^ neo-arterial trajectory which showed a strong overall disease-associated shift: the distribution of cells differed markedly between PAH and controls (Kolmogorov-Smirnov test, p = 0.003), and PAH cells progressed substantially further along this path (median pseudotime Wilcoxon test, p = 0.003, **Fig. 2g**). Both trajectories originated from gCap cells, supporting their function as capillary progenitors in human PAH, analogous to gCap cells in rat disease.

In the human “EBF1 program”, gCap cells upregulated 982 genes and downregulated 161 genes (**Fig. 2h** and **Supplementary Table 9**). These transitioning cells retained core EC markers (*PECAM1*, *CDH5*, *VWF*), while inducing ECM remodeling genes (*FN1*, *COL15A1*, *COL4A1*, *COL4A2*, *SPARC*, *VWA1*), adhesion and migration regulators (*TIAM1*, *MCAM*, *NEDD9*), and the mesenchymal marker *VIM* (encoding vimentin). Thus, similar to rat PAH, *EBF1*^+^ ECs acquire migratory and ECM synthetic features that likely contribute to vascular remodeling.

The *CXCL12*^hi^ neo-arterial trajectory defined a “neo-arterialization program” distinct from the homeostatic arterial endothelial cell state (**Fig. 2i** and **Supplementary Table 9**). These cells upregulated 898 genes and downregulated 161 genes, expressing arterial specification genes (*FOXC1*, *SOX17*, *HEY1*, *GJA5*, *EFNB2*), progenitor marker *CD34*, and Notch ligands (*JAG1*, *JAG2*), together with developmental patterning genes (*HOXD9*, *ENPP2*). The trajectory’s defining feature, *CXCL12* expression, was accompanied by strong expression of ECM remodeling genes (*COL15A1*, *COL4A1*, *SPARC*, *FN1*), angiogenic mediators (*VEGFC*, *LIFR*, *DLL4*), and inflammatory markers (*VCAM1*, *SELE*, *SELP*). Consistent with the trajectory, we visualized significant expansion of *CXCL12*^hi^ neo-arterial ECs in PAH, with prominent localization in the capillary bed and neointima (**Fig. 2j-l**). These findings define a capillary-derived neo-arterialization program in PAH marked by coordinated activation of pro-angiogenic signals, developmental patterning programs, and arterial specification genes.

To map intercellular communication, we performed receptor-ligand analysis of 4,533 interactions across all endothelial and stromal cell types and found 3,556 interactions with elevated activity in PAH (78.4% of total; **Supplementary Table 10**). Among all vascular populations, *EBF1*^+^ ECs were the most polarized endothelial sender (send/receive ratio 1.99, second only to adventitial fibroblasts at 2.27; **Fig. 2m** and **Supplementary Table 10**). *CXCL12*^hi^ neo-arterial ECs were their dominant and most disease-enriched target (89 interactions, 82.4% elevated in PAH; **Supplementary Table 10**), receiving developmental guidance cues and ECM-integrin signals.

Having defined the signaling capacity of EBF1^+^ ECs, we sought to identify the upstream cues responsible for their emergence in disease. In cultured human microvascular ECs, TGFβ-family ligand Activin A and VEGF-A robustly induced *EBF1*, whereas BMP9 suppressed it (**Extended Data Fig. 5**), revealing that imbalances in BMP/TGFβ signaling could induce EBF1 in disease.

Together, these findings establish EBF1^+^ ECs as a conserved pathological feature across preclinical models and human PAH disease. In both rat and human, capillary progenitors undergo bifurcating differentiation into neo-arterial cells and EBF1^+^ ECs, with the latter acting as signaling hubs poised to orchestrate neo-arterialization and stromal recruitment through expression of developmental guidance cues, angiogenic factors, and ECM remodeling programs.

### EBF1^+^ and arterial EC populations arise in parallel from plexus progenitors in the developing lung

Our PAH analyses suggested that *Aplnr*^+^ gCap cells can reprogram into *Ebf1*^+^ ECs expressing vasculotrophic signals normally deployed during embryonic vascular patterning^56^ – a surprising reactivation of morphogenetic programs in the typically quiescent adult endothelium. To trace a possible developmental origin of this program, we examined EBF1 expression and function during PA morphogenesis in the embryonic mouse lung.

The PA forms shortly after lung budding^4,61-63^. At embryonic day 9.0 (E9.0), the ventral foregut evaginates to form the lung primordium, while the splanchnic plexus expands alongside the lung bud to form a primitive endothelial network (the pulmonary plexus) that envelops the branching airways^63,64^. Definitive PA morphogenesis begins around E10, when proximal segments of the primitive PAs first become distinguishable, connecting the cardiac outflow tract via aortic arch VI to the pulmonary plexus^63,65^. The PA then undergoes rapid expansion, branching and maturing in a stereotyped program alongside the airways^63,65^.

To define the progenitor pools and sources of molecular signals controlling PA morphogenesis, and to investigate the possible role of EBF1 in this process, we densely sampled the developing pulmonary vasculature between E11.5 and E15.5 by scRNA-seq (**Fig. 3a**). This yielded 21,222 endothelial and 5,341 stromal cells with high-quality transcriptomes during this critical stage of PA formation (**Supplementary Tables 5, 11**-**12**). Among ECs, the majority (18,827 cells, or 89%) were actively cycling *Aplnr*^+^ plexus progenitors, accompanied by smaller clusters of emerging arterial (*Gja4*^+^*Gja5*^+^*Cxcr4*^+^) and lymphatic (*Ccl21*^+^*Prox1*^+^) ECs. We also identified a minor EC cluster (603 cells, 2.8%) with high *Ebf1* expression, termed “*Ebf1^+^* ECs” (**Fig. 3b, c** and **Extended Data Fig. 6a-d**). These cells expressed canonical EC markers (*Pecam1*, *Cdh5*, *Cldn5*) together with matrix- and mesenchymal-associated features, including the atypical cadherin *Cdh11*^66^ and collagen genes (*Col26a1*, *Col23a1*). These *Ebf1^+^* ECs were molecularly distinct from the *Ebf1*-expressing vascular stromal cells (mainly pericyte and smooth muscle precursors), with no intermediates detected.

**Fig. 3.**
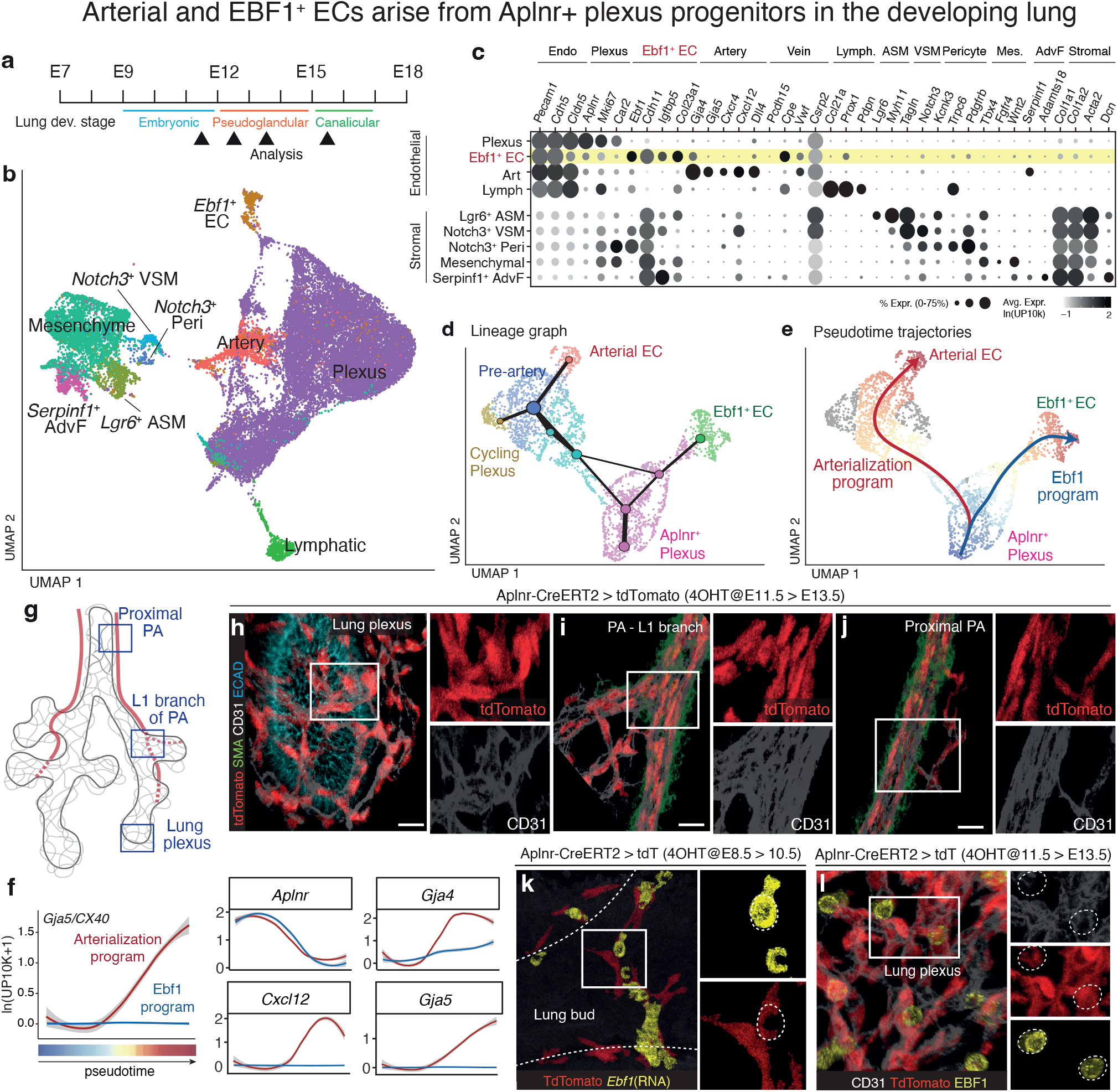
Arterial and EBF1^+^ ECs arise from *Aplnr^+^* plexus progenitors in the developing lung. **a**, Schematic of scRNA-seq profiling of pulmonary vasculature during lung morphogenesis (E11.5-E15.5). Endothelial (CD31*^+^*) and stromal (Epcam^-^CD31^-^ CD45^-^) cells were enriched from wildtype C57BL/6 embryonic lungs at E11.5 (n=12), E12.5 (n=8), E13.5 (n=8), and E15.5 (n=6). Colored bar indicates the corresponding developmental stage. **b**, Integrated UMAP of endothelial and stromal cells from all embryonic ages, annotated by molecular cluster. Abbreviations: VSM, vascular smooth muscle; ASM, airway smooth muscle; AdvF, adventitial fibroblast. **c**, Dot plot of representative marker genes across clusters (fraction expressing and mean expression among expressing cells). **d**, UMAP of developing lung ECs overlaid with a lineage graph. Cells are colored by annotated type. Nodes represent subcluster centroids (size scaled to cell number) and edges are scaled by inferred interaction strength between clusters (see Methods). **e**, Pseudotime analysis overlaid on the UMAP in d, showing bifurcating differentiation of *Aplnr^+^* plexus ECs toward either arterial ECs (red, “arterialization program”) or *Ebf1^+^* ECs (blue, “Ebf1 program”). Arrows indicate principal curves computed by Slingshot. Cells are colored by pseudotime (0-1). **f**, Pseudotime analysis showing trajectory-specific loss of plexus program (*Aplnr*) and induction of arterialization genes along both the Ebf1 and arterialization trajectories. Loess-smoothed expression (*y*-axis) of the indicated genes along the pseudotime trajectories. **g**, Schematic of the developing lung (adapted from Kumar et al. 2014^116^) highlighting relevant vascular structures depicted in **h-j**, including the left proximal pulmonary artery (PA), its first L1 branch, and the distal lung plexus. Note the stereotyped course of the pulmonary artery relative to the airways^63,65^; solid lines indicate PA segments coursing ventrally (visible from this view); dashed lines indicate segments coursing dorsally. **h-j**, Representative whole-mount immunofluorescence of an *Aplnr-*CreERT2*;Rosa26*^tdTomato^lineage-traced embryo at E13.5 following pulse labeling with 4-hydroxytamoxifen (4OHT) at E11.5, showing tdTomato^+^ cells in the lung plexus (**h**), L1 branch of the PA (**i**), and proximal PA (**j**). n=10. **k**, Representative whole-mount smFISH and immunofluorescence image of an *Aplnr-* CreERT2*;Rosa26*^tdTomato^ lineage-traced embryo at E10.5 following pulse labeling with 4OHT at E8.5. n=4. **l**, Representative whole-mount immunofluorescence of an *Aplnr-* CreERT2*;Rosa26*^tdTomato^lineage-traced embryo at E13.5 following pulse labeling with 4OHT at E11.5. n=6. In **k** and **l**, dotted outline highlights *Ebf1* expression in tdTomato**^+^** *Aplnr*-lineage-positive plexus cells. Scale bars, 15μm in **h-j**, 20μm in **k**, and 10μm in **l**.

Lineage reconstruction of these early ECs revealed two trajectories originating from *Aplnr*^+^ plexus ECs at the root, one toward arterial ECs (“arterialization program”) and the other toward *Ebf1^+^* ECs (“Ebf1 program”) (**Fig. 3d, e** and **Supplementary Table 13**). Pseudotime analysis of the arterialization program showed that *Aplnr*^+^ progenitors gradually downregulated *Aplnr*, while upregulating arterial markers (*Cxcl12*, *Gja4*, *Gja5, Eln*), BMP ligands (*Bmp4*, *Bmp6*), and key Notch signaling components (*Hey1*, *Jag1*, *Jag2*, *Notch4*, *Dll4*) that couple cell cycle exit with arterial specification^21^ (**Fig. 3f** and **Extended Data Fig. 7a, b**). To test directly whether *Aplnr*^+^ plexus cells give rise to both pulmonary artery ECs and *Ebf1*^+^ ECs, we performed lineage tracing using *Aplnr-*CreERT2;*Rosa26*^tdTomato^ mice, in which tamoxifen induces tdTomato labeling of *Aplnr*-lineage (*Aplnr*^lin^) cells (**Extended Data Fig. 7c**). We focused on E11.5 onwards, after cardiac progenitors have seeded the proximal primitive PA, at the onset of definitive PA morphogenesis^4,67,68^. Pulse recombination of *Aplnr-*CreERT2 animals at E11.5 robustly labeled plexus ECs (tdTomato^+^CD31^+^) at E12.0 (**Extended Data Fig. 7d**). Over the next two embryonic days (E12.5-E13.5), these *Aplnr*^lin^ plexus ECs progressively incorporated into the growing PA tree, populating new PA branches (**Fig. 3g-j** and **Extended Data Fig. 7e, f)**. By E14.5, the near-complete labeling of the PA tree by *Aplnr*^lin^ plexus ECs, consistent with our prior clonal analysis and lineage tracing results^65^ and comparable to pan-endothelial (*Cdh5*^CreERT2^) fate mapping (**Extended Data Fig. 7g, h**), supports *Aplnr*^+^ plexus progenitors as the major source of PA endothelium during morphogenesis, as they are for the developing alveolar capillary endothelium^20^.

Plexus ECs also gave rise to *Ebf1*^+^ ECs. Examining EBF1 expression in the developing lung, we detected *Ebf1* RNA in the pulmonary vasculature as early as E10.5, coincident with the initial formation of the pulmonary circulation. At this stage, *Ebf1* expression was restricted to a subset of *Aplnr*^lin^ plexus progenitors (tdTomato*^+^Ebf1*^+^ at E10.5; **Fig. 3k**). *Aplnr* lineage tracing beginning at E11.5 showed that, as the pulmonary vasculature expanded, EBF1 expression was most prominent in the pulmonary plexus (CD31^+^tdTomato*^+^*EBF1^+^ at E13.5; **Fig. 3l** and **Extended Data Fig. 8a-d**), where it remained relatively rare. Importantly, this fate transition appeared distinct from arterialization, because EBF1 was rarely expressed in *Gja5*-CreERT2 *(Cx40-*CreERT2)-labeled arterial or pre-arterial ECs (**Extended Data Fig. 8e**). Together, transcriptional trajectory and lineage-tracing analyses demonstrate that *Aplnr*^+^ plexus progenitors give rise to two non-overlapping endothelial fates: arterial ECs of the PA and *Ebf1*^+^ ECs that remain in the pulmonary plexus. We next examined the behavior and function of these *Ebf1*^+^ ECs.

### *Ebf1*^+^ ECs expand and migrate transiently within the pulmonary plexus

To visualize and track *Ebf1*^+^ ECs during pulmonary vasculature development, we generated an *Ebf1-*CreERT2 knock-in mouse line and crossed it with a *Rosa26*^tdTomato^ reporter to lineage label *Ebf1*-expressing (*Ebf1*^lin^) cells (**Extended Data Fig. 9a** and **Supplementary Fig. 1**). Pulse-labeling of *Ebf1*-CreERT2;*Rosa26*^tdTomato^ embryos at the beginning of lung development (E8.5) revealed a broad distribution of *Ebf1*^lin^ cells throughout vascular structures by E9.5-E11.5 (**Fig. 4a** and **Extended Data Fig. 9b-e**). Whole-mount staining confirmed that most *Ebf1*^lin^ cells co-expressed endothelial markers (CD31 and ERG). To map *Ebf1*^lin^ cell fate during PA morphogenesis, we pulse-labeled at E11.5. As before, most *Ebf1*^lin^ cells in the pulmonary plexus were endothelial (CD31^+^ERG^+^tdTomato^+^; **Fig. 4b** and **Extended Data Fig. 10a**, **b**), with a patterned distribution along the expanding PA tree and at nascent branch points (**Extended Data Fig. 10c**). However, plexus-derived *Ebf1*^lin^ ECs were rarely found in the PA intima and showed minimal incorporation into the developing PA endothelium (**Fig. 4c** and **Extended Data Fig. 10d-f**). Instead, beginning at E12, a pool of *Ebf1*^lin^ cells accumulated in the peri-arterial mesenchyme, labeling an emergent stromal compartment (*Ebf1*^lin^ stromal). Could these cells arise from an endothelial-to-stromal transition of the earlier plexus-derived *Ebf1*^lin^ ECs? Two lines of evidence argue against this and instead strongly suggest that *Ebf1*^lin^ stroma is a distinct *Ebf1*-expressing lineage rather than an endothelial derivative. First, lineage-tracing with *Cdh5* or *Aplnr* lines did not label the stromal layer (**Extended Data Fig. 7**). Second, scRNA-seq of the developing lung vasculature identified two distinct *Ebf1-*expressing cell types, with no evidence of interconversion between *Ebf1*^+^ ECs and *Ebf1*-expressing VSM/pericyte precursors.

**Fig. 4.**
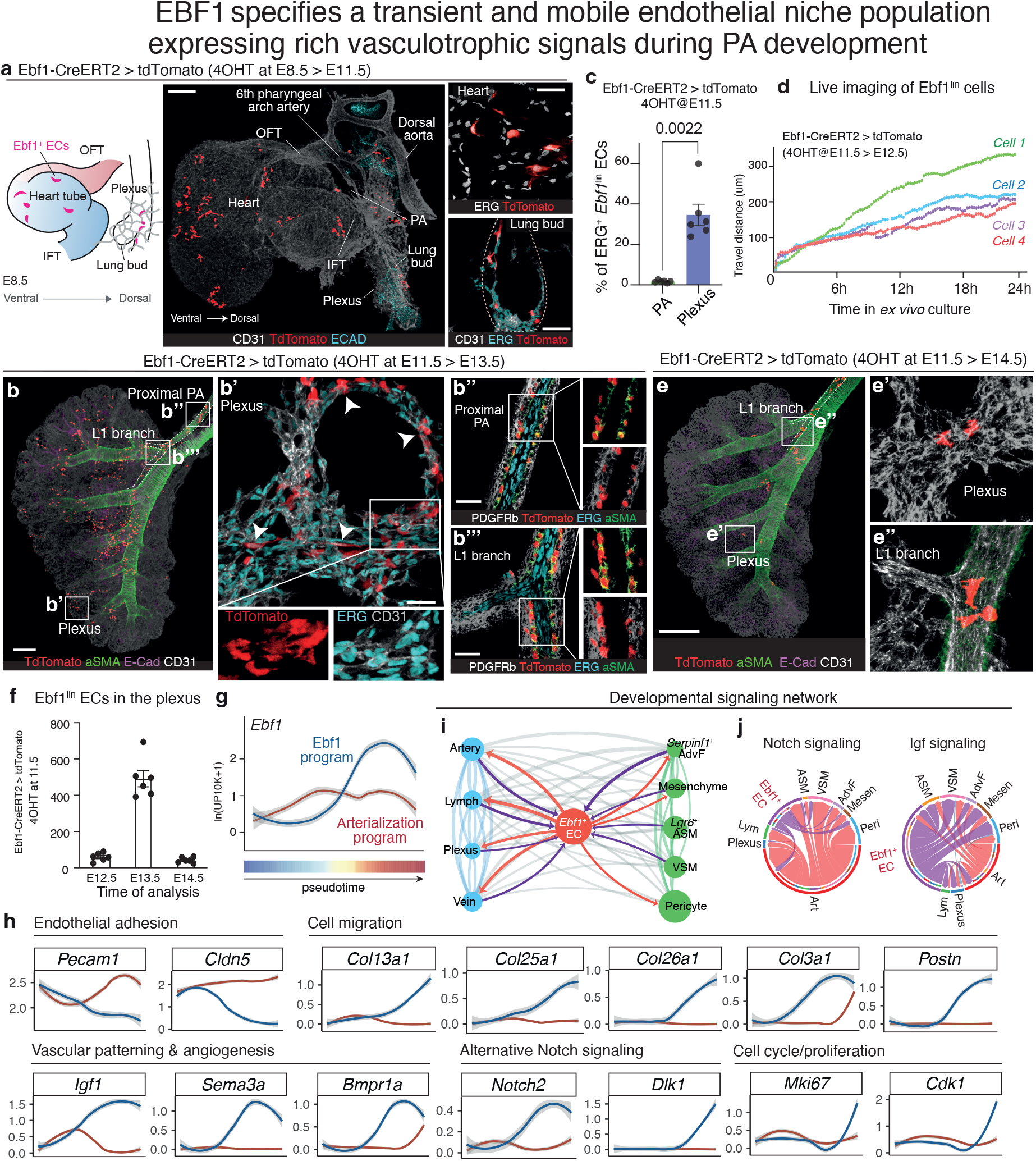
EBF1 specifies a transient and mobile endothelial niche population expressing rich vasculotrophic signals during PA development. **a**, Representative whole mount image of an *Ebf1-*CreERT2*;Rosa26*^tdTomato^ embryo at E11.5 following pulse labeling at E8.5. n=6. Illustration indicates the distribution of *Ebf1* lineage-labeled (*Ebf1*^lin^) ECs within the heart-lung region. OFT, cardiac outflow tract; IFT, cardiac inflow tract; PA, pulmonary artery. Dotted outline traces the lung bud in the inset. **b,** Representative whole-mount images of *Ebf1-*CreERT2*;Rosa26*^tdTomato^embryos at E13.5 showing *Ebf1*^lin^ cells in the lung plexus (**b’**), the proximal PA (**b’’**), and the L1 branch (**b’’’**). n=6. **b’**, Arrowheads and dotted outlines mark *Ebf1*^lin^ ECs (tdTomato^+^CD31^+^ERG^+^) in the lung plexus. **b’’**, **b’’’**, *Ebf1*^lin^ cells in the proximal PA and L1 branch do not express ERG and localize to the smooth muscle actin (SMA)^+^PDGFRb^+^ vessel wall rather than the endothelial intima, consistent with *Ebf1*^lin^ stroma. **c**, Quantification of ERG^+^ *Ebf1^lin^* ECs in proximal PAs and lung plexus at E13.5. ERG positivity was defined as ≥10% overlap between tdTomato^+^ and ERG nuclear surfaces (overlapped volume ratio, Imaris). For PAs, each data point represents one PA (n=6 PAs from 6 embryos, 155 tdTomato^+^ cells total). For plexus, each data point represents one embryo with 3-4 ROIs averaged per embryo (n=6 embryos, 20 ROIs, 353 tdTomato^+^ cells total). Data are presented as mean ± s.e.m. with individual data points shown; statistical comparisons were performed using two-sided Mann-Whitney tests; p values are indicated on the graphs. **d**, Migration distance of *Ebf1*^lin^ cells in *ex vivo* E12.5 lung cultures tracked over 24 hours (See **Extended Data Video 1**). Four representative cells at distinct starting locations are shown. n=6. **e,** Representative whole-mount immunofluorescence of an *Ebf1-*CreERT2*;Rosa26*^tdTomato^embryo at E14.5 following 4OHT at E11.5. Insets show residual *Ebf1*^lin^ cells in the plexus (**e’**) and the L1 branch of the PA (**e’’**). n=6. **f**, Quantification of plexus-located Ebf1^lin^ ECs at indicated embryonic days using Imaris. *Ebf1*^lin^ ECs are defined as *Ebf1*^lin^ (tdTomato^+^) surfaces exhibiting ≥10% overlap with ERG nuclear surfaces (overlapped volume ratio). n=6. **g,** Pseudotime analysis showing *Ebf1* expression in both the Ebf1 program and arterialization program. **h,** Loess-smoothed pseudotime expression profiles for lineage-associated genes, grouped by functional category (endothelial adhesion, cell migration, vascular patterning and angiogenesis, alternative Notch signaling, and cell cycle/proliferation). Significant pseudotime-associated genes were identified by tradeSeq (Full DEG list in **Supplementary Table 13**). **i,** Ligand-receptor interactions inferred from developing lung scRNA-seq data between endothelial and stromal populations. Nodes represent cell states (size scaled by betweenness centrality); edges are scaled by communication probability. Edges involving EBF1^+^ ECs are highlighted (red, signals sent by EBF1^+^ ECs; purple, signals received by EBF1^+^ ECs). **j**, Chord diagrams showing Notch and IGF signaling receptor-ligand interactions between developing lung cell populations. Sectors represent cell states arranged around the circle; ribbons indicate inferred communication links between sender and receiver populations, with ribbon width scaled to communication probability. Scale bars, 200μm in **a**, **b** (left lung lobe), **e**, 25μm in **b’** (plexus), and 20μm in **b’’** (proximal PA) and **b’’’** (L1 branch).

Rather than contributing to the PA endothelium, *Ebf1*^lin^ ECs proliferated and expanded within the pulmonary plexus between E11.5 and E13.5. To visualize their dynamic behavior, we performed time-lapse microscopy on *ex vivo* cultures of *Ebf1* lineage-labeled lungs. We observed clonal expansion and dramatic migration, with individually labeled *Ebf1*^lin^ cells in the pulmonary plexus capable of migrating over 300 μm (approximately spanning 30 plexus cell lengths) within 24 hours (10-12.5 μm/hr; **Extended Data Video 1**, **Fig. 4d**, and **Extended Data Fig. 11a-c**). This expansion window was tightly regulated: *Ebf1*^lin^ cells began to diminish in time-lapse imaging by 48 hours (**Extended Data Fig. 11b**), and whole-mount staining at E14.5 confirmed a significant reduction of lineage-labeled cells (**Fig. 4e, f**). This sharp developmental decline persists into adulthood: *Ebf1*^lin^ ECs remain only sparsely detectable within the healthy pulmonary microvasculature (<5% of CD31^+^ERG^+^ ECs by whole-mount and ∼9.2% by thin-section quantification, **Extended Data Fig. 11d-f** and control group in **Extended Data Fig. 14f**). Thus, *Ebf1*-expressing ECs undergo a transient 10-fold expansion and long-range migration within a tightly regulated developmental window of approximately three embryonic days during PA morphogenesis, followed by rapid attrition, with only rare *Ebf1*^lin^ ECs detectable in the healthy adult lung.

### *Ebf1^+^* ECs provide vasculotrophic signals for the surrounding plexus and stromal progenitors

A molecular analysis of *Ebf1*-expressing ECs explained their behavior and pointed to their function in PA morphogenesis. Trajectory analysis of *Ebf1^+^* ECs revealed a coordinated transcriptional program (the “EBF1 program”) in which *Aplnr*^+^ plexus progenitors initiated *Ebf1* expression, downregulated *Aplnr* and endothelial junction/adhesion molecules (*Pecam1*, *Cldn5*), and upregulated genes linked to ECM remodeling (*Col13a1*, *Col25a1*, *Col26a1, Col3a1*, *Postn*) and cell cycle regulation (*Mki67*, *Cdk1*)^69^, together explaining the proliferative and migratory behavior of *Ebf1*^lin^ ECs (**Fig. 4g, h** and **Supplementary Table 13**).

Receptor-ligand analysis identified *Ebf1*^+^ ECs as a rich source of signals directed toward both endothelial and stromal progenitors (**Supplementary Table 14**). Network analysis revealed that they occupy a uniquely central position (betweenness centrality = 0.236, rank #1, **Fig. 4i**) linking endothelial- and stromal communication during PA development. Candidate vasculotrophic signals (**Fig. 4j** and **Extended Data Fig. 11g**) include the insulin-like growth factor ligands (*Igf1*, *Igf2*)^70^ and *Apln*^71,72^, all implicated in vascular proliferation and migration; Notch signaling components (*Notch2*, *Dlk1*) that could promote arterial specification^44,73,74^; the arterial patterning factor *Cxcr7^75-77^*; and stromal recruitment pathways (TGFβ, PDGFβ, ECM components)^10,78-85^, many of which are dysregulated in PAH.

Thus, the EBF1 program defines a transient population of plexus-resident *Ebf1^+^* ECs that couple developmental vasculotrophic signals with ECM remodeling and cell migratory features, bearing striking similarity to the EBF1^+^ ECs observed in PAH. We propose that these *Ebf1^+^* ECs function as specialized niche cells with unusual mobility to accommodate the expanding arterial tree during PA morphogenesis. We tested this hypothesis using genetic perturbation experiments.

### *Ebf1^+^* ECs control pulmonary vascular development

To delete *Ebf1* specifically in ECs, we crossed a pan-endothelial Cre driver (*Cdh5-*CreERT2) with mice carrying Cre-dependent conditional deletion alleles of *Ebf1* (*Ebf1*^fl^, with *loxP* sites flanking *Ebf1* exons 6-16)^86^, as well as a *Rosa26*^tdTomato^ reporter. Homozygous knockout of *Ebf1* (*Ebf1*^fl/fl^) during the embryonic stage (recombination at E8.5, **Fig. 5a**) resulted in embryonic lethality with no viable *Cdh5-*CreERT2*;Ebf1*^fl/fl^ pups recoverable at birth, consistent with prior global knockout studies showing perinatal lethality^87,88^. In contrast to heterozygous controls (*Ebf1*^fl/wt^) and *Cre^-^* littermates, *Cdh5-*CreERT2*;Ebf1*^fl/fl^ embryos exhibited pronounced defects in pulmonary vasculature development, including around 75% reduction in plexus volume, 40% reduction in PA caliber, defective PA branching, and diminished smooth muscle coverage (**Fig. 5b-d** and **Extended Data Fig. 12a-c**). Single-cell RNA-seq of *Ebf1*^fl/fl^ lungs revealed a dearth of *Ebf1*^+^ ECs, consistent with a cell-autonomous requirement for *Ebf1* in specifying *Ebf1*^+^ ECs (**Supplementary Table 12**). To localize the compartment of endothelial *Ebf1* function, we deleted *Ebf1* in defined endothelial subsets. Plexus-targeted deletion using *Aplnr-*CreERT2 produced a similar developmental delay and reproduced major vascular defects: reduced plexus volume, decreased PA caliber, diminished PA branching, and impaired smooth muscle coverage (**Fig. 5e-h** and **Extended Data Fig. 12d**), phenocopying the pan-endothelial deletion. In contrast, arterial-targeted deletion using *Gja5*-CreERT2 did not impair PA development (**Extended Data Fig. 12e**). Likewise, deleting *Ebf1* from the early *Pdgfr*α*^+^* mesenchymal lineage^89-91^ at E8.5, prior to its differentiation into fibroblast, pericyte, and smooth-muscle fates, had no detectable effects on PA morphogenesis (**Extended Data Fig. 12f**). Together, these compartment-restricted deletions localize the essential early requirement for *Ebf1* specifically to the endothelial plexus.

**Fig. 5.**
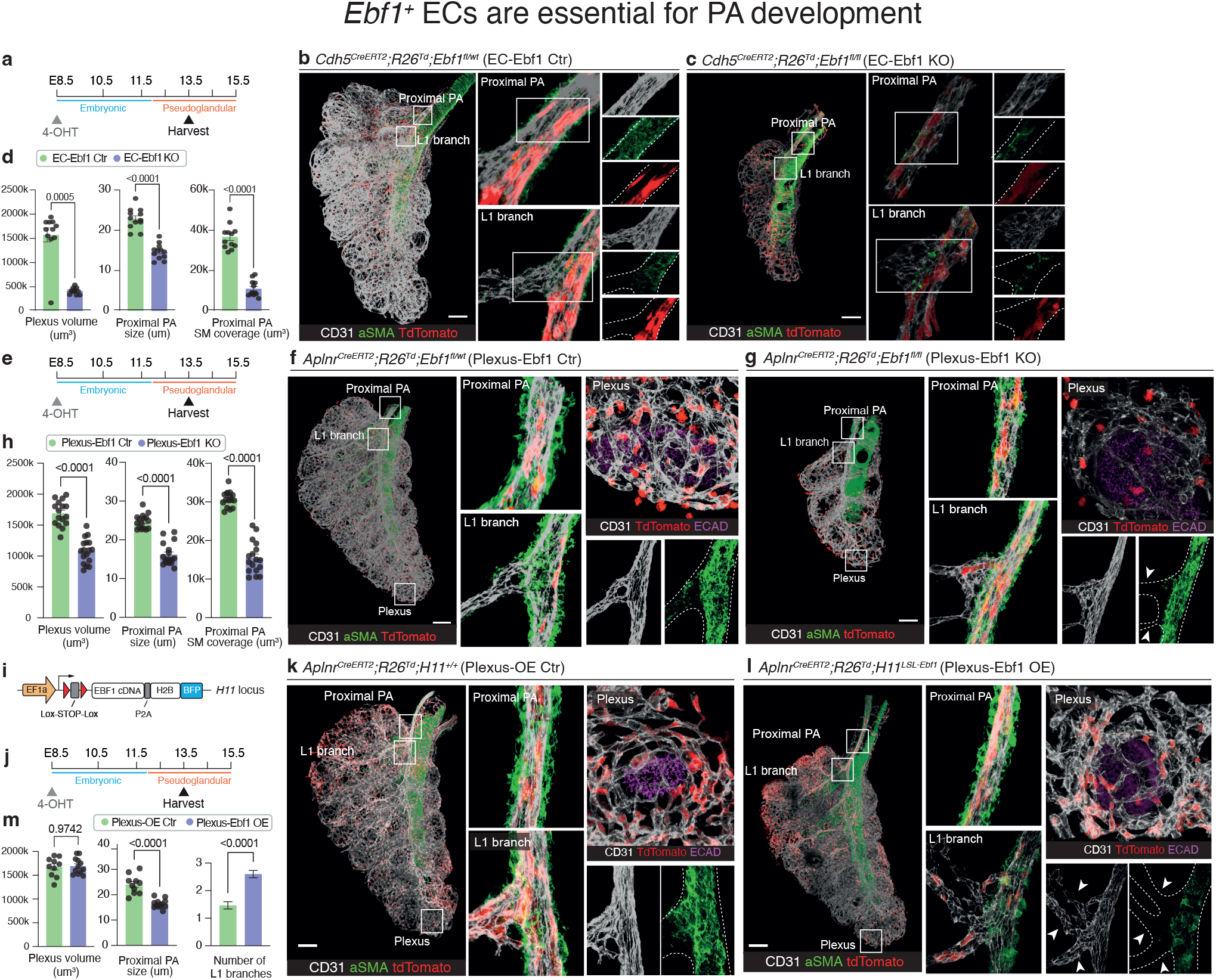
*Ebf1^+^* ECs are essential for PA development. **a,** Experimental schematic for **b-d** (pan endothelial deletion), with recombination initiated by 4OHT at E8.5 (onset of the embryonic stage of lung development). Dashed outlines trace PA branching throughout **b**, **c**, **f**, **g**, **k**, and **l**. **b, c,** Representative whole-mount immunofluorescence of *Cdh5-*CreERT2*;Rosa26*^tdTomato^*;Ebf1*^fl/wt^ (EC-Ebf1 control, n=12) and *Cdh5-*CreERT2*;Rosa26*^tdTomato^*;Ebf1*^fl/fl^ (EC-Ebf1 KO, n=12) lungs at E13.5. **d**, Quantification of plexus volume, PA diameter, and smooth muscle actin (SMA) coverage on proximal PA in the indicated genotypes. n=12. **e,** Experimental schematic for **f-h** (plexus-restricted deletion). **f, g,** Representative whole-mount immunofluorescence of *Aplnr-* CreERT2*;Rosa26*^tdTomato^*;Ebf1*^fl/wt^ (Plexus-Ebf1 control, n=16) and *Aplnr-* CreERT2*;Rosa26*^tdTomato^*;Ebf1*^fl/fl^ (Plexus-Ebf1 KO, n=17) lungs. ECAD marks airway epithelium in plexus insets. Arrowheads indicate an immature L1 branch in Plexus-Ebf1 KO. **h**, Quantification of plexus volume, PA diameter, and SMA coverage on proximal PA in the indicated genotypes (n=16 for Plexus-Ebf1 control, n=17 for Plexus-Ebf1 KO). **i**, Schematic of the LSL-Ebf1 knock-in cassette at the *H11* locus, consisting of an EF1a promoter driving a floxed (Lox-STOP-Lox) transcriptional stop cassette upstream of EBF1 cDNA, a P2A self-cleaving peptide, and a nuclear-localized (H2B) BFP reporter. **j**, Experimental timeline for **k**-**m**. **k, l,** Representative whole-mount immunofluorescence of *Aplnr-*CreERT2*;Rosa26*^tdTomato^*;H11^+/+^*(Plexus-OE control, n=10) and *Aplnr-*CreERT2*;Rosa26*^tdTomato^*;H11*^LSL-Ebf1^ (Plexus-Ebf1 OE, n=12) lungs. ECAD marks airway epithelium in plexus insets. Arrowheads indicate increased immature L1 branches in Plexus-Ebf1 OE. **m**, Quantification of plexus volume, PA diameter, and number of L1 branches in the indicated genotypes (n=10 for Plexus-OE control, n=12 for Plexus-*Ebf1* OE). For **d**, **h**, and **m**, data are presented as mean ± s.e.m.; statistical comparisons were performed using two-sided Mann-Whitney tests; p values are indicated on the graphs. Scale bars, 150μm in **b**, **c**, **f**, **g**, **k**, and **l**.

Notably, both *Cdh5*- and *Aplnr*-lineage-labeled ECs in the *Ebf1*^fl/fl^ lungs could still trace into the PA intima in the absence of *Ebf1*, demonstrating preserved arterial differentiation capacity in these *Ebf1*-null ECs. The PA defects caused by *Ebf1* deletion therefore cannot be explained by a failure of arterial specification. Thus, *Ebf1*^+^ ECs — despite being largely excluded from the PA intima and dispensable for arterial differentiation — are nonetheless required for proper PA formation. Disruption of the EBF1 program in this minor plexus population produces disproportionate defects in PA caliber, branching, and muscularization, consistent with a niche-cell role mediated through paracrine mechanisms.

We next examined how the *Ebf1*^+^ ECs coordinate PA development across developmental time. Deletion of *Ebf1* during the embryonic phase (E8.5) impaired plexus expansion and arterialization: differential expression analysis of these plexus progenitors revealed 796 DEGs in *Cdh5*-CreERT2;*Ebf1*^fl/fl^ lungs (**Extended Data Fig. 12g** and **Supplementary Table 15**), including downregulation of proliferative markers (*Top2a*, *Mki67*), Notch signaling (*Notch3*, *Jag1*, *Jag2*) and angiogenic genes (*Angptl1*, Tek/*Tie2*), alongside upregulation of stress response molecules (*Hspa1a*, *Hspa1b*, *Gadd45g*). These molecular changes explain the impaired survival, proliferation, and arterialization of plexus progenitors observed morphologically. Complementing these loss-of-function data, ectopic *Ebf1* expression in plexus ECs (*Aplnr*-CreERT2; *H11*^LSL-EBF1OE^) at E8.5 reduced proximal PA size and increased immature PA branching (**Fig. 5i-m** and **Extended Data Fig. 12h**), whereas arterial-lineage *Ebf1* overexpression produced no detectable vascular phenotype (**Extended Data Fig. 12i**). Together, these findings demonstrate that, early in PA development, *Ebf1*^+^ ECs are critical for survival, proliferation, and arterialization of plexus progenitors that form the PA tree.

In contrast, later deletion of *Ebf1* during the pseudoglandular phase (E11.5) produced distinct phenotypes: widespread hemorrhage, malformed distal plexus, selective loss of smooth muscle coverage in distal PAs, and dilated pulmonary veins (**Extended Data Fig. 12j-l**). Differential expression analysis of the mesenchyme revealed 194 DEGs (**Extended Data Fig. 12m** and **Supplementary Table 16**), including downregulation of smooth muscle genes (*Actg2*, *Myh11*), actin-binding proteins (*Tagln*, *Cnn1*), and growth factors (*Igf1*, *Pdgfc*). These findings reveal that over developmental time, the function of the *Ebf1^+^* ECs shifts from plexus expansion and arterialization (**Extended Data Fig. 12n**) to the muscularization of mesenchymal progenitors.

To explicitly test the paracrine mechanism, we performed *in vitro* tube-formation assays to assess whether EBF1-expressing cells enhance vascular network formation by neighboring cells. We mixed a majority population of EBF1-negative human microvascular ECs with a minority population of EBF1-overexpressing ECs and quantified network formation. EBF1-overexpressing ECs significantly enhanced total network formation compared to controls, and this enhancement was abolished by IGF1R inhibition (**Extended Data Fig. 13**). These findings demonstrate that EBF1^+^ cells promote angiogenesis in neighboring cells through paracrine IGF1 signaling. This recapitulates the vasculotrophic function observed *in vivo*.

Taken together, these experiments suggest that EBF1 specifies a transient endothelial niche within the vascular plexus that organizes PA morphogenesis through paracrine signaling. This niche dynamically modulates its function over developmental time – first supporting plexus progenitor survival and arterialization to form the PA intima, then orchestrating mesenchymal maturation to build the muscular PA layers.

### The EBF1 program is reactivated in the adult gCap compartment to drive vascular remodeling and PAH

Although most EBF1^+^ ECs disappear upon completion of PA morphogenesis, our developmental and disease analyses suggested that adult pulmonary vascular remodeling in PAH reactivates the EBF1 program within the *Aplnr*-expressing capillary compartment. We therefore directly tested this hypothesis in the Sugen/Hypoxia (SuHx) mouse model of PAH^92-94^, which combines weekly treatment with the VEGFR2 inhibitor SU5416 (20 mg/kg) and chronic hypoxia to produce vascular remodeling of the distal PAs. We performed lineage tracing using *Aplnr*-CreERT2;*Rosa26*^tdTomato^ mice—the same driver used to label plexus progenitors during development and, in the adult, *Aplnr*-expressing gCap cells and their descendants.

In healthy adult lungs, *Aplnr*-lineage gCaps were broadly distributed throughout the capillary bed, with only a rare subpopulation expressing *Ebf1* (3.8% of tdTomato^+^ERG^+^ cells; **Extended Data Fig. 14a-c**), consistent with the sparse *Ebf1*^lin^ labeling established above (**Fig. 4f** and **Extended Data Fig. 11d-f**). Following SuHx, *Aplnr*^lin^ gCap cells accumulated within the occlusive intimal lesions of remodeled arterioles over three weeks, constituting the majority of the ERG^+^ ECs within these lesions (78.8%, p = 0.0002; **Fig. 6a-d**); they also accounted for 82.0% of EBF1^+^ ECs located within these remodeled arterioles (“disease-associated EBF1^+^ ECs”, p = 0.0041; **Fig. 6e**). These findings establish the adult *Aplnr* lineage as the principal endothelial compartment contributing to pulmonary vascular remodeling and the predominant source of disease-associated EBF1^+^ ECs.

**Fig. 6.**
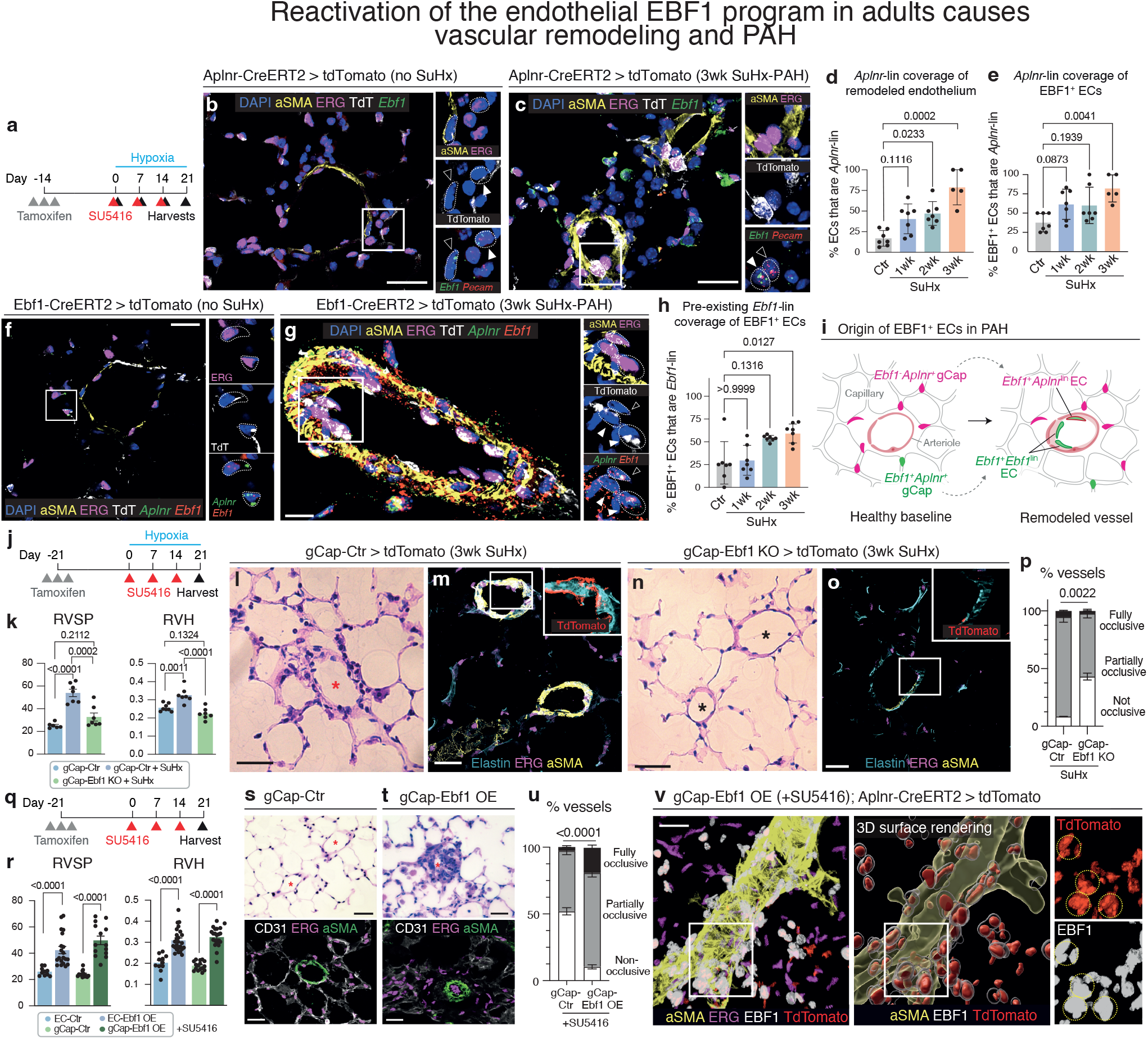
Reactivation of the endothelial EBF1 program in adults causes vascular remodeling and PAH. **a,** Experimental schematic for lineage tracing of *Aplnr-*expressing or *Ebf1*-expressing cells in adult *Aplnr*-CreERT2;*Rosa26*^tdTomato^ or *Ebf1*-CreERT2;*Rosa26*^tdTomato^ mice during SU5416 plus hypoxia (SuHx)-induced pulmonary vascular remodeling in **b-h**. **b**, **c,** Representative lung sections from control *Aplnr*-CreERT2;*Rosa26*^tdTomato^ mice (**b**) or after 3 weeks of SuHx (**c**). **b**, Filled arrowhead indicates an *Aplnr*^lin^ (tdTomato^+^) *Ebf1*^+^ EC; open arrowhead indicates a non-*Aplnr*^lin^ (tdTomato^-^) *Ebf1*+ stromal cell (likely pericyte/VSM). **c**, Filled arrowhead indicates an *Aplnr*^lin^ (tdTomato^+^) *Ebf1*^+^ EC; open arrowhead indicates an *Aplnr*^lin^ (tdTomato^+^) *Ebf1*-negative EC within the occlusive intima. n=5-7 per group. **d**, Quantification of *Aplnr*-lineage coverage of the distal or remodeled endothelium (tdTomato^+^ERG^+^ cells/ ERG^+^ cells) in healthy lungs and after 1, 2 or 3 weeks of SuHx. **e**, Quantification of *Aplnr*-lineage coverage of disease-associated EBF1^+^ ECs (*Ebf1*^+^tdTomato^+^ERG^+^ cells/ *Ebf1^+^*ERG^+^ cells) in healthy lungs and after 1, 2 or 3 weeks of SuHx. **f**, **g**, Representative lung-section images from *Ebf1*-CreERT2;*Rosa26*^tdTomato^ mice at baseline (**f**) and after 3 weeks of SuHx (**g**). **f**, Outlined cells indicate rare pre-existing *Ebf1*^+^*Aplnr*^+^ cells labeled by *Ebf1*-lineage tracing (tdTomato^+^). **g**, Filled arrowhead indicates an *Ebf1*^lin^ (tdTomato^+^) *Ebf1*^+^*Aplnr*^+^ EC within the occlusive intima, consistent with recruitment of a pre-existing *Ebf1*^+^*Aplnr*^+^ cell; open arrowhead indicates a tdTomato-negative *Ebf1*^+^*Aplnr*^+^ EC within the occlusive intima, consistent with *de novo Ebf1* induction in a gCap. n=5 per group. **h,** Quantification of the contribution of the pre-existing *Ebf1* lineage to disease-associated EBF1^+^ ECs (*Ebf1*^+^tdTomato^+^ERG^+^ cells/ *Ebf1^+^*ERG^+^ cells) in healthy lungs and after 1, 2 or 3 weeks of SuHx. **i**, Working model summarizing the complementary *Aplnr*- and *Ebf1*-lineage-tracing results. In the healthy adult lung, rare *Ebf1*^+^*Aplnr*^+^ gCaps reside within the broader *Aplnr*^+^ gCap compartment. During pulmonary vascular remodeling, most disease-associated EBF1^+^ ECs are *Aplnr*-lineage derived. Within this population, a substantial fraction derive from the pre-existing *Ebf1*^+^*Aplnr*^+^ gCap pool (yielding *Ebf1*^+^*Ebf1*^lin^ ECs), with an additional contribution consistent with *de novo Ebf1* induction in previously *Ebf1*^-^*Aplnr*^+^ gCaps (yielding *Ebf1*^+^*Aplnr*^lin^ ECs). Cell outline denotes lineage origin (magenta, *Aplnr*-lineage; green, *Ebf1*-lineage), and fill denotes current Ebf1 expression (green, *Ebf1*-positive). **j**, Experimental schematic for genetic deletion of *Ebf1* in adult *Aplnr*^+^ gCap cells during SuHx-induced PAH. Tamoxifen was administered to *Aplnr*-CreERT2;*Rosa26*^tdTomato^ control mice (gCap-Ctr) and *Aplnr*-CreERT2;*Rosa26*^tdTomato^*;Ebf1*^fl/fl^ mice (gCap-Ebf1 KO) before SuHx exposure. Mice received SU5416 (20 mg/kg weekly) and were maintained under hypoxia for 3 weeks. **k**, Hemodynamic measurements in gCap-Ctr mice at baseline and in gCap-Ctr or gCap-Ebf1 KO mice after 3 weeks of SuHx. Right ventricular systolic pressure (RVSP) was measured by right heart catheterization. Right ventricular hypertrophy (RVH) is presented as right ventricle mass/ (left ventricle + septum) mass. n=7 mice per group. **l-o**, Representative H&E and immunofluorescence images from gCap-Ctr and gCap-Ebf1 KO lungs after 3 weeks of SuHx. The red asterisk marks a remodeled PA in gCap-Ctr lungs, and black asterisks mark less-remodeled PAs in gCap-Ebf1 KO lungs. Insets show tdTomato (TdT)-labeled *Aplnr*^lin^ ECs within the vascular endothelium. n=7 mice per group. **p**, Quantification of vascular occlusion in gCap-Ctr and gCap-Ebf1 KO lungs after 3 weeks of SuHx. Pulmonary arterioles were classified as non-occlusive, partially occlusive or fully occlusive. n=7 mice per group. **q,** Experimental schematic for **r-v. r,** Hemodynamic measurements of SU5416-treated EC control (*Cdh5-* CreERT2*;Rosa26*^tdTomato^*;H11^+/+^*, n=10), gCap controls (*Aplnr-*CreERT2*;Rosa26*^tdTomato^*;H11^+/+^*, n=14), EC-Ebf1 OE (*Cdh5-*CreERT2*;Rosa26*^tdTomato^*;H11*^LSL-Ebf1^, n=20), and gCap-Ebf1 OE (*Aplnr-*CreERT2*;Rosa26*^tdTomato^*; H11*^LSL-Ebf1^, n=14) mice at day 21. **s, t**, Representative H&E and immunofluorescence images of d21 lung sections from gCap-Ctr and gCap-Ebf1 OE mice treated with SU5416. n=6. Red asterisks mark distal PAs. **u,** Quantification of vascular occlusion in day 21 lungs from gCap control and gCap-Ebf1 OE mice with SU5416. n=10. **v**, Representative whole-mount immunofluorescence and 3D surface rendering of an occlusive lesion from SU5416-treated gCap-Ebf1 OE lung. Insets show EBF1^+^*Aplnr*^lin^ (tdTomato^+^) ECs, derived from gCaps; dashed outlines mark these cells in the inset channels. n=6 mice per group. For **d**, **e**, and **h**, 3-5 distal PAs (<40 μm) or vascular regions were analyzed per timepoint; ROI results were averaged to generate one biological replicate value. For **d**, **e**, **h**, **k**, **p**, **r** and **u**, data are presented as mean ± s.e.m.; statistical comparisons for **p**, **r**, and **u** were performed using two-sided Mann-Whitney tests; comparisons for **d**, **e**, **h**, and **k** were performed using Kruskal-Wallis tests followed by Dunn’s multiple-comparisons tests; p values are indicated on the graphs. Scale bars, 25μm in **b**, **c**, **f**, **g**, **l**, **m**, **n**, **o**, **s**, and **t**, and 20μm in **v**.

To determine the contribution of the rare pre-existing *Ebf1* lineage, we labeled *Ebf1*-expressing cells before disease onset using *Ebf1*-CreERT2;*Rosa26^tdTomato^*. At baseline, most *Ebf1*-lineage ECs expressed *Aplnr* (78.4% of *Ebf1*^lin^ ERG^+^ cells), establishing adult *Ebf1*-lineage ECs as a rare subpopulation harbored within the *Aplnr*^+^ gCap compartment (**Fig. 6f** and control group in **Extended Data Fig. 14f**). Despite constituting less than 10% of the endothelium at baseline, the *Ebf1*-lineage progressively accumulated within the occlusive intimal lesions of remodeled arterioles, accounting for 59.9% of actively EBF1-expressing ECs after three weeks of SuHx (p = 0.0127; **Fig. 6f-h** and **Extended Data Fig. 14d-f**). The same lineage relationships were observed by *Aplnr*-CreERT2 and *Ebf1*-CreERT2 labeling in a mechanistically distinct HDM model of PAH^32^ (**Extended Data Fig. 14g-m**). Together, the predominance of *Aplnr*-lineage cells among disease-associated EBF1^+^ ECs, with substantial contribution from the rare pre-existing *Ebf1*^lin^ subset within this gCap pool, indicates at least two sources of disease-associated EBF1^+^ ECs: the pre-existing *Ebf1*^lin^ gCaps and an additional contribution consistent with *de novo Ebf1* induction from the broader gCap compartment (**Fig. 6i**). Because *Ebf1*-CreERT2 also labels a distinct pericyte/VSM lineage, *Ebf1* lineage tracing alone cannot formally exclude a non-endothelial source; however, complementary *Cdh5*-CreERT2 tracing labeled 94.4% of disease-associated EBF1^+^ ECs, confirming their predominantly endothelial origin (**Extended Data Fig. 14n-p**).

To determine whether the progressive accumulation of pre-existing *Ebf1*-lineage ECs resulted from preferential proliferation, we quantified Ki67-expressing cells separately within *Ebf1*-lineage and non-*Ebf1*-lineage ECs during SuHx-mediated remodeling. Although proliferation increased markedly in both populations following SuHx exposure, the fraction of Ki67^+^ ECs remained comparable between lineages at each time point examined (**Extended Data Fig. 14q-s**), arguing against preferential proliferation of the pre-existing *Ebf1* lineage as the basis for its accumulation. The enrichment of *Ebf1*-lineage ECs within remodeled vessels may instead reflect earlier proliferation, preferential recruitment to or retention within remodeling vessels, or a combination of these processes.

We next asked whether reactivation of the EBF1 program within adult gCaps is required for pulmonary vascular remodeling. To test this genetically, we deleted *Ebf1* in adult *Aplnr*^+^ gCap cells before SuHx exposure (**Extended Data Fig. 15a**). *Aplnr*-CreERT2 controls exposed to three weeks of SuHx developed severe pulmonary hypertension, with an RVSP of 54.0 ± 8.4 mmHg. In contrast, gCap-specific *Ebf1* deletion reduced RVSP by 39%, to 32.7 ± 9.5 mmHg, decreased RVH by 31% (p = 0.0002 and p < 0.0001, respectively; **Fig. 6j, k**), markedly attenuated intimal occlusion (**Fig. 6l-p**), and reduced the abundance of *Ebf1^+^Aplnr*^lin^ cells in the distal pulmonary arterioles (**Extended Data Fig. 15b**). Thus, reactivation of the EBF1 program within the adult gCap lineage is required for occlusive vascular remodeling and PAH.

We returned to the rat model in which we identified neointimal EBF1^+^ ECs, testing the endothelial requirement for *Ebf1* through an independent loss-of-function approach. We used the endothelial-tropic AAV-BI30 capsid^95^ with a dual-vector PiggyBac system to stably deliver a membrane-targeted tdTomato reporter for permanent lineage labeling of transduced ECs and their progeny, together with either a scrambled or *Ebf1*-targeting shRNA. AAVs were administered two weeks before PAH induction (**Extended Data Fig. 15c** and Methods). In control (scrambled-shRNA) animals, intratracheal delivery initially transduced 44 ± 3.7% of ERG^+^ pulmonary ECs, predominantly in the distal capillary bed (**Extended Data Fig. 15d**). These reporter-labeled ECs were subsequently detected within occlusive intimal lesions of remodeled arterioles, independently supporting a capillary endothelial origin. In *Ebf1*-shRNA animals, endothelial *Ebf1* knockdown protected against PAH: treated rats showed no rise in RVSP or RVH and no evidence of occlusive vascular remodeling (**Extended Data Fig. 15e-h**). *Ebf1* silencing also blunted neo-arterialization, reducing capillary *Dll4* expression and PA number and branching (**Extended Data Fig. 15i-k**). Thus, endothelial *Ebf1* is required in rat PAH for both neo-arterialization and occlusive vascular remodeling.

Having established that endothelial EBF1 is required for this coordinated remodeling program, we asked whether reactivating EBF1 in adult capillaries is sufficient to drive it. We induced *Ebf1* expression in adult endothelium using either pan-endothelial (*Cdh5*-CreERT2) or capillary-restricted (*Aplnr*-CreERT2) drivers. Although postnatal *Ebf1* overexpression induced distal PA muscularization (**Extended Data Fig. 16a-d**), it did not produce severe PAH. We therefore challenged *Ebf1*-overexpressing mice with SU5416 (20 mg/kg/week), a VEGFR2 inhibitor that alone does not cause severe PAH in mice without concurrent hypoxia (**Fig. 6q**). After three weeks, Cre-only control mice exposed to SU5416 showed no evidence of disease. In contrast, both pan-endothelial and capillary-restricted *Ebf1* overexpression elevated RVSP and RVH, and produced prominent intimal remodeling of distal PAs, with the most severe occlusive lesions observed in capillary-targeted animals (**Fig. 6r-u**). Thus, endothelial EBF1 reactivation converts a subthreshold endothelial injury into severe pulmonary hypertension, with the strongest effect when confined to the gCap compartment.

To determine whether gCap-derived cells directly contribute to neointimal lesions, we examined *Aplnr* lineage-labeled cells in gCap-Ebf1 OE lungs treated with SU5416. *Aplnr*^lin^ ECs (tdTomato^+^ERG^+^) were broadly detected within remodeled PAs (**Extended Data Fig. 16e, f**) and comprised a substantial fraction of EBF1^+^ cells within them (EBF1^+^tdTomato^+^ERG^+^; **Fig. 6v**), establishing a direct gCap contribution to the formation of occlusive intimal lesions. In parallel, the neo-arterialization marker *Cxcl12* was strongly induced both within occlusive distal PAs and throughout the surrounding capillary bed, accompanied by increased PA number and branching (**Extended Data Fig. 16g-k**). Thus, EBF1 reactivation in adult capillaries, together with endothelial injury, generates both occlusive intimal lesions and neo-arterialization.

This gain-of-function phenotype recapitulates the bifurcating differentiation of capillary progenitors inferred in rat and human PAH and mirrors the plexus-progenitor bifurcation that builds the pulmonary arterial tree during development. Together with the loss-of-function data, these findings establish that endothelial EBF1 is necessary for PAH pathogenesis and, when coupled with endothelial injury, sufficient to reactivate the embryonic artery-forming program, driving coordinated occlusive vascular remodeling and neo-arterialization.

## Discussion

Originally discovered for its role in B cell development, the pioneer transcription factor EBF1 has since been recognized to regulate cell fates in multiple neural and mesodermal lineages across metazoans^26,27,88,96-102^. Here, our search for novel regulators of PAH revealed a new function for EBF1 in specifying a rare population of mobile and transient endothelial cells that dynamically organize PA morphogenesis through vasculotrophic signaling. In adulthood, pathological reactivation of the EBF1 program within the gCap compartment reconstitutes this embryonic artery-forming niche in a maladaptive reprise of development in which, instead of disappearing, these reactivated EBF1^+^ ECs persist in neointimal lesions to drive PAH.

Our findings show that the developing pulmonary vascular plexus, an immature endothelial network that envelops the nascent lung buds and expands in parallel with branching morphogenesis of the airways and the PA tree^63,65^, progressively gives rise to pulmonary arterial ECs that form the PA intima^65^ and a novel, transient population of *Ebf1*-expressing cells (*Ebf1*^+^ ECs) that are nevertheless critical for PA development. Together with prior clonal analysis and lineage tracing establishing that plexus progenitors are multipotent, giving rise to multiple vascular fates including pulmonary arterial ECs^20,65^, our single-cell transcriptomic analysis resolves two bifurcating differentiation trajectories within the *Aplnr*-expressing progenitor pool and defines their gene-expression programs, which we term the “arterialization program” and “Ebf1 program”. These *Ebf1*^+^ ECs proliferate and migrate extensively yet incorporate minimally into the PA intima. If *Ebf1*^+^ ECs do not trace into the arterial wall, how do they control PA morphogenesis? Our scRNA-seq and genetic experiments suggest that *Ebf1*^+^ ECs function as niche cells during PA formation. Through coordinated expression of vasculotrophic signals, *Ebf1*^+^ ECs organize plexus progenitor expansion and arterialization together with maturation of surrounding stromal progenitors to build the PA tree (**Extended Data Fig. 16l**). This function localizes to the endothelial plexus during the early developmental window we tested: pan-endothelial and plexus-targeted deletion produced severe vascular defects, whereas arterial or early stromal deletion did not impair PA morphogenesis. Our data do not exclude stromal *Ebf1* function at later developmental stages or in other contexts. Related work examines the cellular, molecular, and patterning mechanisms of pulmonary artery morphogenesis (Bogard et al., in preparation).

Unlike most epithelial stem cell niches with well-defined locations and architectures^103,104^, *Ebf1*^+^ ECs act dynamically in space and time, broadly distributing throughout the pulmonary plexus and intermingling with the pre-arterial ECs, while sequentially activating distinct functions through vasculotrophic signals. They also differ fundamentally in lineage organization: whereas epithelial niches are typically stromal support cells derived from distinct germ layers, *Ebf1*^+^ ECs and arterial ECs are sister cell types arising from *Aplnr*^+^ plexus progenitors through bifurcating differentiation, enabling their coupled specification. The organizing function of *Ebf1*^+^ ECs may require the expression of genes associated with the partial EndoMT-like endothelial phenotype to achieve delamination and long-distance migration (reminiscent of the neural crest^105^) to accommodate the rapidly expanding arterial tree. Similar to the Spemann-Mangold organizer, the classic vertebrate signaling center that patterns the embryonic body axis^106^, *Ebf1^+^* ECs are transient yet essential organizing centers, coordinating cell fate decisions and vascular architecture across tissue compartments to induce PA formation. Whether analogous transient endothelial organizers exist in other vascular beds remains an open question with implications for vascular development and disease.

Perhaps the most unusual feature of these *Ebf1*^+^ ECs is their dynamic 10-fold expansion (E11.5-E13.5) followed by rapid attrition (E14.5) from the embryonic lungs within just three embryonic days. While *Aplnr*^+^ plexus progenitors rely on *Ebf1*^+^ ECs for expansion and differentiation during this window, the developmental EBF1^+^ niche contracts dramatically after PA morphogenesis, leaving only a rare residual population within the adult *Aplnr*^+^ gCap compartment. The tightly regulated disappearance of *Ebf1*^+^ ECs is reminiscent of developmental programmed cell death, including the transcriptionally controlled apoptosis in precisely 131 of the 1,090 cells in the *C. elegans* somatic lineage^107^ and the BMP-driven interdigital apoptosis critical for mammalian finger formation^108^. Their tight temporal control may restrict *Ebf1*^+^ EC activity to PA morphogenesis, and failure of this control following pathological reactivation may instead drive vascular remodeling in PAH.

BMP signaling is an attractive candidate for controlling this developmental program, given the well-established role of *BMPR2* deficiency in PAH. Our data reveal that BMP/TGFβ signaling imbalance regulates EBF1 expression itself: PAH-associated factors (Activin A, VEGF-A) induce EBF1 whereas BMP9 suppresses it. Whether BMP signaling also governs the fate or survival of EBF1^+^ ECs remains an open question. However, BMP signaling is unlikely to be sufficient for their induction, nor is endothelial *Ebf1* expression itself sufficient for severe vascular remodeling: *Bmpr2* heterozygosity alone did not generate a distinct EBF1^+^ EC population in rats, while forced endothelial *Ebf1* expression produced severe PAH only in the context of vascular injury. The two-hit requirement mirrors clinical observations that severe PAH requires multiple insults, with *BMPR2* mutations alone showing incomplete penetrance^15,109^. Vascular injury therefore appears to serve at least two separable functions, expanding the EBF1 lineage and activating their artery-forming program, while BMP signaling may gate their induction and, potentially, termination. Nevertheless, this tight regulation of EBF1 has clinical relevance, as sotatercept – the first disease-modifying therapy approved for PAH – sequesters activin-class ligands to rebalance BMP/TGFβ signaling^110-113^, though it is not curative. One possibility is that this incomplete response reflects the challenge of persistence: while restoring BMP/TGFβ balance may limit EBF1 reactivation, it may not eliminate the EBF1^+^ ECs already established in neointimal lesions that sustain and drive vascular remodeling. In the future, identifying the signals that not only induce EBF1 but also control the fate of the cells it specifies may provide the dual targeting required to reverse vessel-occluding neointima in PAH.

Our unified approach to PA development and disease, focusing on the embryonic function and adult reactivation of EBF1, has several advantages. The stereotyped developmental sequence of the PA allowed us to map cellular behavior and function at single-cell resolution, and the tools available for mouse genetic experiments led us to identify EBF1^+^ ECs as specialized cells derived from *Aplnr*^+^ plexus progenitors that organize artery formation through vasculotrophic signaling. The causal role of EBF1 in disease was then established through complementary genetic experiments. The artery-forming program, defined by a bifurcating differentiation trajectory observed in rat PAH and established genetically in mouse development, was reconstituted experimentally in adult mice. Capillary-restricted *Ebf1* overexpression combined with vascular injury generated both arms of this program: EBF1^+^ neointimal organizers and neo-arterial structures, producing severe PAH. Conversely, endothelial *Ebf1* knockdown protected against disease and blocked neo-arterial formation, while gCap-specific genetic deletion attenuated hemodynamic disease and neointimal occlusion. These experiments establish EBF1 as a key regulator of a coordinated artery-forming program whose precise temporal control – transient in development, persistent in disease – determines outcome.

Yet disease is not a direct replay of development (**Extended Data Fig. 16m**). Two differences distinguish pathological reactivation of the EBF1 program. (1) Origin: whereas embryonic EBF1^+^ ECs arise through normal plexus differentiation, adult reactivation requires both BMP dysregulation and vascular injury, and occurs through expansion and accumulation of a rare residual EBF1 lineage within the *Aplnr*^+^ gCap compartment, together with an additional contribution from *de novo* EBF1 induction in gCaps, consistent with a dual origin of EBF1^+^ ECs in PAH. (2) Persistence: embryonic EBF1^+^ ECs disappear after organizing PA formation, while pathological EBF1^+^ ECs generate neointima and persist there. Within neointimal lesions, EBF1^+^ ECs organize this remodeling through vasculotrophic signaling that instructs neo-arterialization while recruiting pericytes, stimulating smooth muscle cells, and activating fibroblasts – a distorted redeployment of their embryonic PA-organizing function. Although hybrid endothelial-mesenchymal phenotypes in PAH and other vascular diseases (notably atherosclerosis) are speculated to arise from EndoMT lineage conversion, our results indicate that EBF1^+^ ECs retain endothelial identity while acquiring migratory and ECM-remodeling features. This partial phenotype shift may enable their organizer function and, when pathologically sustained, drive vascular remodeling.

The resulting disease model addresses enduring questions about the cellular origins and mixed phenotype of PAH neointimal lesions, the surprising remodeling potential of normally quiescent pulmonary endothelium^114^, and the source of coordinated angiogenic cues. Previous lineage tracing studies using distinct PH models have variably implicated endothelium^52^ or smooth muscle^115^ as cell of origin for neointima, reflecting the heterogeneity in mouse PH models and the challenge of mapping them onto human disease. Our approach – interrogating preclinical rat and clinical human data at high resolution, then using genetics to guide lineage perturbation – identified the pulmonary capillary bed as the source of both neointimal and neo-arterial cells. This aligns with recent reports of capillary arterialization in a genetic HIF-2α-driven model of PH^34^. While multiple canonical endothelial markers and complementary genetic strategies support our lineage interpretations, the *Aplnr*- and *Ebf1*-lineage experiments were performed in separate animals using CreERT2 drivers that may have different recombination efficiencies, so their labeling cannot be directly compared to precisely quantify the relative contribution of pre-existing lineage expansion versus *de novo* induction; additional fate-mapping approaches, including dual recombinase systems, will be needed to distinguish the lineage contributions and cellular behaviors of distinct *Ebf1*-expressing subsets and to resolve lineage intermediates among closely apposed endothelial and mural compartments during development and across diverse PAH etiologies.

Important questions remain open. What are the direct chromatin targets of EBF1 that regulate endothelial reprogramming *in vivo*? What signals govern EBF1^+^ EC fate and trigger their death in development – and why do they persist in disease? Nevertheless, our findings establish that EBF1 controls an embryonic artery-forming program that, when pathologically reactivated in adulthood, drives pulmonary vascular remodeling and PAH. Targeting this program may intercept vessel-occluding disease before it becomes irreversible, offering new therapeutic strategies for PAH and potentially other vascular diseases where developmental organizing programs are pathologically reactivated.

## Supplementary Tables

**Supplementary Table 1 Marker genes in rat PAH scRNA-seq analysis**

**Supplementary Table 2 Cell counts per cluster in rat PAH scRNA-seq analysis**

**Supplementary Table 3 Pseudotime DEGs in rat PAH scRNA-seq analysis**

**Supplementary Table 4 Receptor-ligand interactions in rat PAH scRNA-seq analysis**

**Supplementary Table 5 Integration quality metrics across scRNA-seq datasets**

**Supplementary Table 6 Demographic information for human samples.**

**Supplementary Table 7 Human PAH scRNA-seq dataset composition**

**Supplementary Table 8 EBF1 differential expression in human PAH**

**Supplementary Table 9 Trajectory program DEGs in human PAH scRNA-seq analysis**

**Supplementary Table 10 Receptor-ligand interactions in human PAH scRNA-seq analysis**

**Supplementary Table 11 Marker genes in mouse lung developmental scRNA-seq analysis**

**Supplementary Table 12 Cell counts per cluster in mouse lung developmental scRNA-seq analysis**

**Supplementary Table 13 Pseudotime DEGs in mouse lung developmental scRNA-seq analysis**

**Supplementary Table 14 Receptor-ligand interactions in mouse lung developmental scRNA-seq analysis**

**Supplementary Table 15 DEG analysis of early endothelial EBF1 deletion**

**Supplementary Table 16 DEG analysis of late endothelial EBF1 deletion**

## Methods

### Mice

Adult mice used in the postnatal experiments were between the ages of 8 and 12 weeks and maintained on a C57BL/6 background (both male and female). Mice were housed with a 12-h light–dark cycle at 18–23 °C and 40–60% humidity in the animal facility at the Palo Alto Veterans Medical Center. Most mouse lines were described earlier^117-119^, including the *Aplnr-* CreERT2, *Cdh5-*CreERT2, *Gja5-*CreERT2, and *Rosa26*^tdTomato^ reporter (The Jackson Laboratory, B6.Cg-*Gt(ROSA)26Sor^tm9(CAG-tdTomato)Hze^*/J, Stock# 007909), and *Ebf1*^flox^ (The Jackson Laboratory, B6.Cg-*Ebf1^tm2.1Mbu^*/J, Stock#028104). For embryonic studies, timed pregnancies were determined by defining the day of plug formation as E0.5. In pulse-chase, *Ebf1* deletion, and *Ebf1* overexpression studies, E/Z-4-hydroxytamoxifen (4OHT, Cayman Chemical, #17308), used for its shorter half-life and precise temporal control, was dissolved in ethanol at a concentration of 2.5 mg/ml and was injected into the peritoneal cavities of pregnant dams at 2.5-10mg/kg body weight. In all other lineage tracing experiments, tamoxifen (Sigma, T5648) was dissolved in corn oil at a concentration of 10 mg/ml and was given to the pregnant dams through oral gavage at 50-100 mg/kg. Dosing and dissection schedule for each study are indicated in their corresponding data sections. For adult lineage-tracing experiments, *Aplnr*-CreERT2;*Rosa26*^tdTomato^, *Cdh5*-CreERT2;*Rosa26*^tdTomato^, and *Ebf1*-CreERT2;*Rosa26*^tdTomato^ mice received tamoxifen (75 mg/kg, s.c) 14 days before disease induction or tissue collection to label *Aplnr*-expressing gCap cells, *Cdh5*-expressing ECs, or *Ebf1*-expressing cells, respectively. For adult gCap-specific *Ebf1* deletion studies, *Aplnr*-CreERT2; *Rosa26^tdTomato^*;*Ebf1^fl/fl^*mice and corresponding reporter controls received tamoxifen (200 mg/kg, s.c.) once daily for three consecutive days, from day - 23 to day -21, followed by a 21-day washout before disease induction. Other postnatal Cre-induction experiments used tamoxifen at the doses and schedules indicated in the corresponding experimental sections. All mouse experiments and care were conducted in accordance with the procedures approved by the Institutional Animal Care and Use Committee (IACUC) guidance.

### Generation of *Ebf1 ^P2A-CreERT2^* and *H11^lsl-Ebf1^* mouse strains

The *Ebf1*^CreERT2-P2A^ mouse strain was generated by homology-directed repair at the endogenous *Ebf1 locus* aided by CRISPR-Cas9 endonuclease activity in C57BL/6 mice by Applied Stemcell Inc. A targeting construct of cDNA encoding tamoxifen-inducible Cre recombinase (CreERT2) and a P2A self-cleaving peptide sequence followed by an FRT-flanked neomycin-resistance cassette was generated. A mixture of 2 guide RNAs, spCas9 mRNA, and the repair donor was injected into the C57BL/6J embryos. Recognition sequences of the gRNAs used in this study are as follows: 5g2F: 5’-ACC AAT GTC ACG TGT GGA TTG GG-3’ and 3g2R: 5’-TCA AGG CAA TTC TTT CAC ATG GG-3’. The donor *Ebf1* sequence (see details in **Supplementary Fig. 1)**, with homology arms at both sides, was constructed and used as a repair donor template. Linearized donor DNA and CRISPR-Cas9 complex were injected into C57BL/6 fertilized zygotes, which were then implanted into the oviducts of pseudopregnant female mice. Founders were identified by genotyping.

To generate the *H11^lsl-Ebf1^* mouse strain, a similar strategy was used by Applied Stemcell. A transgene of EF1a promoter-STOP (floxed)-*Ebf1* cDNA-P2A-H2B-mTagBFP2 was knocked into H11 locus, in C57BL/6J background using CRISPR-Cas9 technology. A mixture of two guide RNAs, spCas9 mRNA, and donor DNA was injected into the C57BL/6J embryos. Recognition sequences of the guided RNAs are as follows: H11-L3: 5’-TGA TGG AAC AGG TAA CAA AGG -3’ and H11-R2: 5’-GCT TAT CTT GAA CTC TTG TGG-3’. The donor DNA (see sequence details in **Supplementary Fig. 2)**, with homology arms at both sides, was constructed and used as a repair donor template. The injected embryos were transferred into the oviduct of CD-1 foster mothers. Pups were screened by PCR. Founders were sequenced and then bred to wild-type C57BL/6J male to produce F1 heterozygotes.

### Rat models of PAH

All rats used in this study were between the ages of 4-8 weeks (both male and female). For the *Bmpr2^+/-^* model, *Bmpr2^+/-^* and *Bmpr2^+/+^*rats were constructed by the Nicolls lab in the SAGE laboratory^14^. PAH was induced through intratracheal instillation of adenovirus expressing *Alox5* using a published protocol.

For monocrotaline model^29^, Sprague-Dawley rats (6-8 weeks, 120-150 g) were given a single subcutaneous injection of monocrotaline (60 mg/kg in DMSO, Sigma) or vehicle control.

To model PAH with immunodeficiency^30^, inbred athymic nude rats (constructed by the Nicolls lab; 4-6 weeks) were treated with the VEGFR2 inhibitor SU5416 (Cayman Chemical, subcutaneous) at 20 mg/kg dissolved in DMSO, whereas control animals received matched vehicle. For all models, animals were maintained in normoxia (20% O□) and monitored daily. All animals were euthanized at post-treatment week 3 for hemodynamics.

To assess the efficacy of endothelial-targeted knockdown of *Ebf1* in a rat PAH model, we generated an adeno-associated virus packaged with the endothelial-tropic AAV-BI30 capsid (an engineered AAV9 variant reported to efficiently transduce endothelial cells)^95^. The knockdown cassette contained three independent miR30-based shRNA hairpins targeting *Ebf1* concatenated within a single vector to enhance robustness; matched control viruses encoded non-targeting scramble hairpins (**Supplementary Fig. 3**). Viruses were produced at high titer (Vectorlab) and delivered to adult rats by intratracheal instillation (1×10¹¹ vg per rat in sterile PBS). Where indicated, vectors also encoded tdTomato to enable tracking of transduced cells. This platform additionally incorporates a two-vector PiggyBac system to enable stable genomic integration and durable lineage labeling *in vivo*: one vector encodes a hyperactive, nuclear-localized PiggyBac transposase (hyPBase; *pAAV[Exp]-SFFV>hyPBase(ns):T2A:H2B-mNeonGreen:WPRE*), and the companion vector carried the PiggyBac-flanked (transposon) cargo comprising the shRNA cassette (and a tdTomato reporter). Following co-transduction, hyPBase catalyzes integration of the PB-flanked cargo into the host genome, supporting long-term persistence of the knockdown cassette and stable labeling of transduced endothelial cells and their progeny. Following viral delivery, animals were maintained for 2 weeks to allow stable expression and knockdown before PAH induction (or treated in parallel with disease induction as indicated). For PAH studies, rats were subjected to the indicated *Bmpr2* PAH protocol, followed by hemodynamic assessment and histological analysis of pulmonary vascular remodeling at endpoint. Endothelial transduction was evaluated by whole-mount immunofluorescence staining for endothelial markers (ERG) together with reporter signal. AAV-BI30 endothelial transduction efficiency was quantified from three-dimensional confocal image stacks using Imaris. Three vascular volumes (100 × 100 × 50 μm) were analyzed per rat. ERG nuclei were identified as endothelial cells, and AAV-transduced ECs were defined as tdTomato ERG cells. Transduction efficiency was calculated as tdTomato ERG cells divided by total ERG cells × 100. Values from the three vascular volumes were averaged to generate a single biological replicate for each animal (n = 5 rats). The *Ebf1* target sequences were 5′-TCCTGGCAGTCTCTGATAACAT-3′ (TRCN0000086580), 5′-GGCGCGACTGTGATCATCATAG-3′ (TRCN0000321383), and 5′-CGCCTTCTAACCTGCGGAAATC-3′ (TRCN0000374141). Experimental protocols were approved by the Veteran Affairs Palo Alto Animal Care and Use Committee.

### Mouse models of PAH

For SU5416 plus hypoxia (SuHx) studies^92^, mice received SU5416 (20 mg/kg, s.c., dissolved in DMSO) on days 0, 7 and 14 and were maintained under hypoxia (10% O_2_) beginning on day 0. Lungs were collected after 1, 2 or 3 weeks of SuHx exposure. To trace the fate of Ebf1-expressing cells during SuHx-induced pulmonary vascular remodeling, *Aplnr*-CreERT2;*Rosa26^tdTomato^*, *Ebf1*-CreERT2;*Rosa26^tdTomato^*and *Cdh5*-CreERT2;*Rosa26^tdTomato^* mice were administered tamoxifen 14 days before disease induction. This washout period minimized continued recombination after initiation of vascular injury. Control animals were collected at the end of the washout period without disease induction. To test whether *Ebf1* expression in adult gCap cells is required for PAH, *Aplnr*-CreERT2; *Rosa26^tdTomato^*;*Ebf1^fl/fl^*mice (gCap-Ebf1 KO) and *Aplnr*-CreERT2;*Rosa26^tdTomato^* littermate controls (gCap-Ctr) received tamoxifen 21 days before the first SU5416 injection. Right ventricular systolic pressure and right ventricular hypertrophy were measured on day 21, followed by lung collection. To test whether endothelial *Ebf1* overexpression sensitizes adult mice to vascular injury, *Cdh5*-CreERT2;*Rosa26tdTomato;H11LSL-Ebf1* (EC-Ebf1 OE) and *Aplnr-* CreERT2;*Rosa26tdTomato;H11LSL-Ebf1* (gCap-Ebf1 OE) mice, together with the corresponding Cre-positive H11+/+ littermate controls, were induced with tamoxifen (200 mg/kg s.c. for 3 continuous days) 21 days before SU5416 treatment. Following this washout period, mice were maintained under normoxic conditions and received SU5416 (20 mg/kg, s.c.) on days 0, 7 and 14. Right ventricular systolic pressure and right ventricular hypertrophy were measured on day 21, followed by lung collection. For postnatal endothelial *Ebf1* overexpression, *Cdh5-CreERT2;Rosa26tdTomato;H11LSL-Ebf1* mice and Cre-negative littermate controls received tamoxifen on postnatal days 0-2 and were analyzed at postnatal day 30.

For the house-dust-mite-induced model^32^, PAH was induced in adult mice (8-10 weeks) by house-dust-mite (HDM) extract (Stallergenes Greer, catalog number: XPB70D3A25). To induce PAH, animals were dosed intranasally with HDM (50 µl in PBS) on a 5-days-on/2-days-off schedule for 5 weeks. Lungs were collected immediately following hemodynamic assessment. Control mice received 50 µl PBS on the same schedule. For lineage-tracing experiments, *Ebf1*-CreERT2;*Rosa26^tdTomato^, Aplnr*-CreERT2;*Rosa26^tdTomato^*and *Cdh5*-CreERT2;*Rosa26^tdTomato^* mice were administered tamoxifen 14 days before house-dust-mite-induced remodeling, and were administered house-dust-mite extract intranasally for 3 and 5 weeks using the same dosing schedule.

In the TNFα overexpression mouse model^31^, lung-specific TNFα-overexpressing mice (*Tg^+/SFTPC-Tnfa^*) and transgene-negative littermate controls were maintained on a C57BL/6 background and analyzed at 8 weeks of age. When the transgene-positive mice exhibited spontaneous PAH and pulmonary vascular remodeling, lungs were inflation-fixed, paraffin-embedded, and sectioned for histological assessment.

### Hemodynamic measurement

Animals were anesthetized with ketamine hydrochloride and xylazine injected intraperitoneally prior to right heart catheterization. Right ventricular systolic pressure (RVSP) measurements were obtained by insertion of a Micro Tip pressure transducer (model SPR-671, 1.4F; Millar Instruments) through the jugular vein into the RV. Signals were recorded continuously with a TC-510 pressure control unit 236927/R17 (Millar Instruments) coupled to a Bridge Amp (AD Instruments). Data were collected with the Powerlab 4/30 data acquisition system (AD Instrument) and analyzed with Chart Pro software (AD Instrument). RV was dissected from left ventricle (LV) and septum (S) and weighed. Fulton index of RV/(LV+S) was calculated to determine the degree of right ventricular hypertrophy (RVH). All quantification was performed in a blinded fashion.

### Tissue preparation for mouse and rat lungs

Immediately after hemodynamic measurements, lungs were perfused via the right ventricle with ice-cold PBS until blanched. The trachea was cannulated and lungs were inflated with 1.5% agarose in PBS, then excised and fixed in 10% formalin overnight (4 °C or room temperature).

For preparation of thin sections, fixed lungs were rinsed in PBS and cryoprotected in sucrose/PBS at 4 °C (10% w/v for 24 h, 20% w/v for 24 h, then 30% w/v until tissue sank).

Lungs were blotted, oriented in a cryomold, embedded in OCT, and snap-frozen on dry ice (or in a dry ice/isopentane slurry). Blocks were stored at -80 °C until sectioning. Cryosections (5-10 µm) were cut on a cryostat, mounted on charged slides, air-dried, and stored at -20 °C until immunofluorescence staining.

For preparation of thick (vibratome) sections, fixed lungs were rinsed in PBS and embedded in agarose (2% in PBS) in an embedding mold. Once solidified, the agarose block was trimmed and sectioned on a vibratome in ice-cold PBS to generate 150-200 µm sections. Free-floating sections were collected with a paintbrush, transferred to PBS in multi-well plates, and stored at 4 °C until staining and clearing (typically within a week). For long-term storage, sections were dehydrated through a graded ethanol series (e.g., 50%, 70%, 80%, 90%, 100%) and stored at - 20 °C in 100% ethanol until use.

### Immunofluorescent staining of frozen lung sections

For staining of frozen sections (5-10 µm), lung slides were equilibrated to room temperature, rehydrated in PBS, and permeabilized with 0.2% Triton X-100 in PBS for 10 min. Sections were blocked in 5% normal donkey (or goat) serum with 1% BSA in PBS for 1 h at room temperature. Primary antibodies against α-smooth muscle actin (Sigma, C6198; 1:400), CD31/PECAM1 (BD Bioscience, 550274, 1:100), ERG (Abcam, ab92513, 1:100), EBF1 (Abcam, ab221033, 1:100, or SCBT, sc-13706, 1:50), tdTomato (Rockland, 600-901-379; 1:50), Ki67 (Invitrogen, A5-1450, 1:50), or FLAG (Sigma, F1804, 1:50) were diluted in blocking buffer and incubated overnight at room temperature in a humidified chamber. Sections were washed 3× in PBST and incubated with species-appropriate Alexa Fluor-conjugated secondary antibodies (2 h at room temperature, protected from light). Alexa Fluor 750 hydrazide (Abnova, U0261; reconstituted to 0.5 mg/ml in PBS, 1:200) or Alexa Fluor 633 hydrazide (Invitrogen, A30634; 0.5 mg/ml, 1:800) were used to visualize elastin fibers and were added along with secondary antibodies. Nuclei were counterstained with DAPI (5-10 min), sections were washed, and coverslipped with antifade mounting medium. Images were acquired using Leica Stellaris confocal microscope at 40X or 63X magnification. Maximum projection of the Z-stack images was performed using Imaris 10 or 11.

### Wholemount immunofluorescent staining for thick lung sections

For staining of thick lung sections (150-200 µm), vibratome sections were equilibrated to room temperature, rinsed in PBS, and permeabilized in 0.5% Triton X-100 in PBS for 60 min with gentle agitation. Sections were blocked in 5% normal donkey (or goat) serum + 1% BSA in PBS for 2 h at room temperature. Primary antibodies against α-smooth muscle actin (Sigma, C6198; 1:600), CD31/PECAM1 (BD Bioscience, 550274; 1:500), ERG (Abcam, ab92513; 1:300), EBF1 (Abcam, ab221033; 1:200, or SCBT, sc-13706; 1:200), tdTomato (Rockland, 600-901-379; 1:250), Ki67 (Invitrogen, A5-1450, 1:200), or FLAG (Sigma, F1804; 1:100) were diluted in blocking buffer and incubated for 48-72 h at 4 °C with gentle rocking. Sections were washed in blocking buffer for 4-5 h, then incubated with species-appropriate Alexa Fluor-conjugated secondary antibodies for 48 h at 4 °C. Alexa Fluor 750 hydrazide (Abnova, U0261; reconstituted to 0.5 mg/ml in PBS, 1:200) or Alexa Fluor 633 hydrazide (Invitrogen, A30634; 0.5 mg/ml, 1:800) were used to visualize elastin fibers and were added along with secondary antibodies. Sections were washed in PBST. For optical clearing prior to imaging, sections were dehydrated through a graded methanol series (50%, 80%, 100%; 30 min each) followed by a second 100% methanol step, then transferred into BABB (benzyl alcohol:benzyl benzoate, 1:2) until optically transparent (∼30-60 min, depending on thickness). Cleared sections were mounted in BABB using glass spacers. Images were acquired on a Leica Stellaris confocal microscope (40× or 63×) using Z-stacks. Maximum intensity projections and 3D rendering were performed using Imaris 10 or 11.

### Human lung samples

Lung sections were obtained from PAH patients following lung transplantation. Control subject tissue was collected during lobectomy or pneumonectomy for localized lung cancer; pulmonary arteries were studied at a distance from the tumor areas. Transthoracic echocardiography was performed preoperatively in the control cohort to rule out PAH. All lung slices were obtained deidentified from the Pulmonary Hypertension Breakthrough Initiative (PHBI) Network (#54182).

For scRNA-seq of human lung tissues, samples from patients with PAH and healthy donors were obtained through the Pulmonary Hypertension Breakthrough Initiative (PHBI) under a PHBI Network Institutional Review Board-approved protocol, with informed consent obtained at the transplant procurement sites. The use of archived human lung tissues and cells was approved by the University of Arizona Institutional Review Board (#1907824872). All tissues and cells were provided in a de-identified manner and were therefore considered non-human subjects research for the purposes of this study. Single cells were isolated from lung tissue and loaded onto the 10x Genomics Chromium Single Cell Controller to generate barcoded single-cell libraries. Single-cell libraries were generated using the 10x Genomics Fixed RNA Profiling kit. Single-cell RNA sequencing was performed on an Illumina NovaSeq X Plus platform using paired-end 150-bp reads.

### Single molecule fluorescent *in situ* hybridization (smFISH) and immunofluorescent staining and quantification

smFISH was performed using the RNAscope Multiplex Fluorescent Reagent Kit v2 (Advanced Cell Diagnostics, ACD) according to the manufacturer’s instructions. Briefly, paraffin sections were deparaffinized in xylene (2 × 5 min) and rehydrated through 100% ethanol (2 × 5 min). Target retrieval was performed by heating slides in RNAscope Target Retrieval solution in a steamer for 17 min. Slides were rinsed in distilled water, immersed in 100% ethanol for 5 min, air-dried, treated with RNAscope Hydrogen Peroxide for 10 min at room temperature, and rinsed briefly in water. Sections were then incubated with Protease Plus at 42 °C for 32 min, followed by probe hybridization, signal amplification, and HRP-based detection per the ACD protocol. Probes and fluorophore channels were as follows: human *EBF1* (C3; CF680 tyramide, Biotium 92196), mouse *Ebf1* (C4; CF680 tyramide, Biotium 92196), rat *Ebf1* (C4; CF680 tyramide, Biotium 92196), human *PECAM1* (C4; CF594 tyramide, Biotium 92174), rat *Pecam1* (C3; CF594 tyramide, Biotium 92174), mouse *Pecam1* (C3; CF594 tyramide, Biotium 92174), human *CXCL12* (C1; CF488A tyramide, Biotium 92171), rat *Dll4* (C2; CF488A tyramide, Biotium 92171), mouse *Cxcl12* (C2; CF488A tyramide, Biotium 92171), human *APLNR* (C2; CF594 tyramide, Biotium 92174), and mouse *Aplnr* (C1; CF594 tyramide, Biotium 92174). Following smFISH, sections were processed for immunofluorescence. Slides were incubated with primary antibodies (ERG, αSMA, tdTomato, and/or PDGFRβ) overnight at room temperature, washed in PBST (3 × 5 min), and incubated with species-appropriate fluorescent secondary antibodies for 2 h at room temperature. After PBST washes (3 × 5 min), nuclei were counterstained with DAPI (5 min), and slides were mounted in ProLong Gold antifade reagent (Invitrogen). Images were acquired on a Leica Stellaris confocal microscope using 40× or 63× objectives.

Quantification of smFISH signals (*EBF1, CXCL12*, and *DLL4*) and immunofluorescent signals of ERG in ECs in lung sections was performed using QuPath. For each PAH subject, 5-8 regions of interest (ROIs) containing vascular remodeling (plexiform lesions and/or occlusive PAs) were analyzed; for controls, 5-8 ROIs containing small PAs (<100 µm in human, <50 µm in mouse and rat) without remodeling were selected using the same criteria. ROIs were drawn to capture the vessel wall/lesion while excluding tissue folds, airspaces, and areas with obvious artifacts or high background. Multichannel fluorescence images were imported with verified pixel size and channel assignments (DAPI for nuclei; ERG for nuclear endothelial annotation; *EBF1*, *DLL4,* or *CXCL12* RNAscope channel for transcript signals). Cells were segmented by DAPI-based nuclear detection with fixed parameters applied across all samples, followed by a uniform cytoplasmic expansion to approximate cell boundaries. ERG ECs were identified by applying a uniform ERG intensity threshold to segmented nuclei and classifying ERG cells as ECs. ERG EC density was reported as ERG cells per ROI area and/or as a fraction of total nuclei within each ROI. *EBF1*, *Dll4*, or *CXCL12* RNAscope signals were quantified in the same segmented cells using an automated, threshold-based detection approach (or puncta detection where applicable), with a single set of detection thresholds defined on representative images and then applied identically to all ROIs. For each subject, ROI-level measurements were averaged to generate a single value. Endothelial-restricted *EBF1*, *Dll4*, *CXCL12* expression was reported as the fraction of ERG ECs that were *EBF1*-, *Dll4*-, or *CXCL12*-positive, and as *EBF1, Dll4,* or *CXCL12* signals per ERG EC (e.g., puncta/cell or intensity/positive area per cell). All measurements were exported as Prism files for downstream statistical analysis.

Quantification of lineage-traced ECs was performed using QuPath on multichannel confocal images acquired using identical settings across experimental groups. Nuclei were segmented using DAPI, and ERG^+^ nuclei were classified as ECs using a fixed intensity threshold. *Ebf1*^lin^ ECs were defined as tdTomato^+^ERG^+^ cells in *Ebf1*-CreERT2;*Rosa26^tdTomato^*lungs. *Aplnr*-lineage *Ebf1*^+^ ECs were defined as tdTomato^+^ERG^+^ cells meeting the predefined *Ebf1* smFISH positivity threshold in *Aplnr*-CreERT2; *Rosa26^tdTomato^*lungs. For each mouse, 5-8 vascular regions of interest containing small PAs and adjacent capillary regions were analyzed. *Ebf1* expression within *Aplnr-*lineage ECs was calculated as *Ebf1*^+^tdTomato^+^ERG^+^ cells / tdTomato^+^ERG^+^ cells × 100; *Aplnr*-lineage contribution to *Ebf1*^+^ ECs as *Ebf1*^+^tdTomato^+^ERG^+^ cells / *Ebf1*^+^ERG^+^ cells × 100; and lineage contribution to the remodeled endothelium as tdTomato^+^ERG^+^ cells/ ERG^+^ cells × 100.

For proliferation analyses, proliferating ECs were defined as Ki67^+^ERG^+^ cells, and the contributions of the Ebf1-lineage or the non-Ebf1-lineage to the proliferating endothelial population were calculated as TdTomato^+^Ki67^+^ERG^+^ cells / tdTomato^+^ERG^+^ cells × 100 or TdTomato^-^Ki67^+^ERG^+^ cells / tdTomato^-^ERG^+^ cells × 100. Three remodeled vessels or vascular regions were analyzed per mouse and averaged to generate one biological replicate value.

### Quantification of PA branching

Mouse or rat lungs were vibratome-sectioned at 150-200 µm. For consistency across animals, sections were collected from the mid region of the right lung lobes at comparable anatomical depth (excluding apical and basal extremes). Free-floating sections were stained for 24 h with anti-αSMA-Cy3 (Sigma, C6198, 1:500), hydrazide-Alexa Fluor633 (Invitrogen, A30634, 1:500), and *Lycopersicon esculentum* (tomato) lectin-DyLight 488 (Invitrogen, L32470, 1:1000), washed in PBST for 1 h, and cleared in benzyl alcohol/benzyl benzoate (BABB). Cleared sections were imaged on a Leica THUNDER microscope using a 20× objective. Image processing and analysis were performed using Fiji (ImageJ). Images were calibrated using the embedded scale bar (Analyze → Set Scale). Lung tissue regions were manually outlined as ROIs (polygon tool) and ROI area was recorded (mm²). PAs were identified by a patent/dark lumen surrounded by hydrazide- and αSMA-positive vessel wall labeling. The total number of PAs and PA branching points were counted manually within each ROI. PA density was reported as PAs per mm²; branching density as branch points per mm²; and branching per PA as branch points divided by total PAs. For each animal, values from 2-3 sections were averaged to generate a single biological replicate. Quantification was performed blinded to genotype/treatment group. Manual quantification of branch-point-based vascular metrics and area normalization are consistent with established vascular network quantification frameworks^120^.

### Quantification of vascular occlusion

The degree of occlusion of small peripheral pulmonary arteries (<50 µm) was assessed by double-staining of lung sections with an anti-αSMA (Sigma, C6198; 1:200) and anti-CD31 (Biolegend, 102516, 1:100) antibodies. For each vessel, the area within the internal elastic lamina (αSMA^+^) and the luminal area (CD31^+^) were traced in ImageJ to calculate percent free luminal area (lumen/internal elastic lamina × 100). Vessels were categorized as non-occlusive (≥75% free lumen), partially occlusive (75-25%), or fully occlusive (<25%). 15-35 vessels per animal were analyzed. For each animal, the percentage of vessels in each category was calculated by dividing the number of vessels in that category by the total number of vessels analyzed for that animal. Animal-level percentages were used as independent biological replicates for statistical analysis.

### Whole mount immunofluorescence staining of embryonic samples

Embryos were collected and dissected in ice cold PBS and immediately fixed in 4% paraformaldehyde (PFA) at 4 °C for 1 h, followed by two 15 min washes in PBS at 4 °C. Samples were incubated with primary antibodies diluted in PBS containing 0.5% Triton X-100 (0.5% PBST; ≥5 sample volumes) for 2-3 days at 4 °C with gentle agitation. Samples were washed in 0.5% PBST (≥20 volumes) for 6 h at room temperature (RT), then incubated with Alexa Fluor-conjugated secondary antibodies diluted in 0.5% PBST (≥5 sample volumes) for 2-3 days at 4 °C. Following secondary incubation, samples were washed in 0.5% PBST (≥20 volumes) for 6 h at room temperature. Stained embryos were post-fixed in 4% PFA at 4 °C for 1 h, dehydrated through a graded methanol series, and cleared in benzyl alcohol:benzyl benzoate (1:2; BABB, Sigma, #305197, #B6630) prior to imaging. All steps were performed with gentle continuous shaking; antibody incubations and BABB clearing were carried out in 15 ml glass tubes. Primary antibodies used, at indicated concentrations, were: EBF1 (Abcam, ab221033; 1:150); PECAM1 (BD Biosciences, #553370, 1:500) or PECAM1 (R&D, AF3628,1:250); Cy3-conjugated αSMA (Sigma, C6198; 1:500); tdTomato (Rockland, #44567; 1:300); E-cadherin (Invitrogen, ECCD-2, 1:500); ERG (Abcam, ab92513, 1:500); PDGFRb (Invitrogen, 14-1402-82, 1:150). All secondary antibodies were Alexa Fluor conjugated (350, 488, 555, 647, 750, Invitrogen, 1:400). DAPI (Invitrogen, D1306; reconstituted in PBS, 2 μg/ml) was used to stain nuclei. All images were taken using Leica Stellaris confocal microscope. Maximum projection images and 3D reconstruction of the Z-stack images were performed using Imaris 10 or 11.

### Quantification of embryonic images

All quantifications were performed using Imaris (10 or 11) using confocal images acquired with 20x, 40x, or 63x objectives. To quantify numbers of Ebf1-lineage, ERG endothelial cells, Z-stacks from *Ebf1^CreERT2^* reporter samples (stained with tdTomato, ERG, CD31, αSMA, DAPI) were analyzed in Imaris. tdTomato cells were segmented using the *Surfaces* workflow with *Split Touching Objects* (estimated diameter ∼5-10 µm) and a quality threshold set to include discrete tdTomato signals while minimizing background. Segmentation was manually inspected in 3D and parameters were adjusted as needed to prevent merged objects or missed cells. Regions of interest (ROIs) were defined to restrict analyses to plexus or PA compartments, and signals outside ROIs were excluded. Total tdTomato objects were obtained from the Statistics tab. Co-localization with ERG was quantified in Imaris Vantage by measuring overlap between tdTomato objects and ERG-defined nuclear surfaces (overlapped volume ratio), and by reporting the fraction of tdTomato objects meeting a predefined ERG-overlap threshold (>10%). Quantification outputs were exported from Vantage as .csv files for downstream analysis. For proximal PA, 6 PA regions (from different embryos) were analyzed yielding a total of 155 tdTomato single cells overlapping with ERG surfaces. To avoid pseudo-replication, cell-level measurements within each vessel were aggregated, and each PA was treated as one data point for plotting and statistical comparisons. For plexus analyses, 20 plexus regions from 6 embryos were analyzed, resulting in 353 tdTomato single cells overlapping with ERG surfaces. Plexus measurements were aggregated per embryo, and each embryo was treated as one data point for plotting and statistical comparisons.

To quantify aSMA coverage, proximal PA regions were defined by manually tracing the vessel to generate a CD31-based *Surface* object. An additional *Surface* object was generated from the αSMA channel with thresholding and smoothing optimized to capture continuous αSMA signal without merging distinct smooth muscle bands. Surface area values for αSMA and vessel were obtained from the *Statistics/Results* outputs.

To quantify the lung plexus volume, Z-stacks containing the CD31 channel were imported into Imaris and voxel dimensions/channel assignments were verified in the *Properties* panel. Images were cropped to a defined lung plexus volume (2,250,000 µm³). A 3D *Surface* object was generated from the CD31 channel using the *Add Surfaces* wizard with an intensity threshold set to best capture vascular signal. Segmentation quality was verified in 3D renderings and orthogonal views. Plexus volume (µm³) was obtained from the *Statistics* tab and exported as .csv files.

To quantify PA diameters, orthogonal cross-sections were generated at different positions along the vessel axis using the Orthoslice tool in Imaris. Diameters were measured in each cross-section using measurement points or line segments. At least three orthogonal sections per vessel were quantified and averaged to generate a representative diameter for each PA segment.

### Human lung microvascular endothelial cell (hMVEC-L) culture

Primary human lung microvascular endothelial cells (hMVEC-L) were purchased from Lonza (#CC-2527) and cultured in complete Endothelial Growth Medium-2 Microvascular (EGM-2 MV BulletKit, Lonza, #CC-3202) according to the manufacturer’s instructions. Cells between passages 4 and 7 were used in this study. To generate Ebf1-overexpression cells (Ebf1^OE^), hMVEC-L cells were transduced with *AdEBF1* for 3 days. Control cells (Ebf1^neg^) were transfected with adenovirus with empty vectors.

### *In vitro* tube formation assay in hMVEC-L

Tube formation assays were performed to assess the vasculotrophic effects of EBF1-expressing endothelial cells (Ebf1^OE^) on surrounding endothelial populations (Ebf1-negative, Ebf1^neg^). Human microvascular endothelial cells (hMVECs) were used for all experiments. Briefly, Growth factor-reduced Matrigel (Corning) was thawed overnight at 4 °C and polymerized in μ-Slide Angiogenesis chambers (ibidi, #81506) at 37 °C for 60 min. hMVEC-L were trypsinized and resuspended in complete growth medium. For co-culture, 9,500 EBF1-negative hMVEC-L were mixed with 500 hMVEC-L expressing either *EBF1* (Ebf1^OE^) or GFP control (adenoviral transduction for 3 days) and seeded in 50 µl per well onto polymerized Matrigel. Where indicated, EBF1-negative cells were pretreated for 24 h and maintained during the assay with BMS-754807 (1 µM, IGF1R inhibitor; Selleckchem #S1124), AMD3465 (5 µM, CXCR4 antagonist; #S2879), or LY-411575 (5 µM, γ-secretase inhibitor; #S2714); vehicle was used as control. Tube networks were imaged by phase-contrast microscopy (10×) between 0-6 h after seeding at 37 °C and 5% CO□. Quantitative analysis of tube formation was performed using Fiji/ImageJ with the Angiogenesis Analyzer plugin. Image calibration was performed based on the scale bar for each image (Analyze → Set Scale; known distance = 150 µm; unit = µm). Images were converted to 8-bit format (Image → Type → 8-bit). Manual thresholding was not applied. Images were subsequently converted to RGB format (Image → Type → RGB Color) prior to analysis. The Angiogenesis Analyzer was run using phase-contrast settings with identical parameters applied across all experimental conditions. Primary angiogenesis parameters quantified included total master segment length, number of master junctions, number of meshes, and mean mesh size. All image acquisition and analysis settings were kept constant within each experiment.

### Real-time reverse transcription quantitative PCR (RT–qPCR) for EBF1 mRNA

hMVEC-L were treated for 16-18 h with recombinant human ligands (TGF-β, IGF1, CXCL12, BMP9/GDF2, Activin A, GDF11, VEGF165, TNFα, IL-6, and IL-1β) at the indicated working concentrations (**Extended Data Fig. 5**). Where indicated, cells were transduced with *shEBF1* adenovirus (MOI 5) or non-targeting control 3 days before ligand stimulation. Total RNA was extracted using RNeasy Plus Mini Kit (QIAGEN, #74136) and reverse-transcribed with the High-Capacity cDNA Reverse Transcription Kit (Applied Biosystems, #4374966). qPCR was performed using Power SYBR Green Master Mix (Applied Biosystems, #4367659) on a QuantStudio 7 Flex (384-well format; 40 cycles, 95 °C for 15 s and 60 °C for 60 s). Relative expression was calculated by the ΔΔCt method and normalized to 18S rRNA. Primer sequences were: EBF1 forward: 5’-CGCAACTCAAGCAGCGTATC-3’, EBF1 reverse 5’-TCTGTTTCAC GGCTGAGACC-3’; 18S rRNA forward 5’-CTGCCATTAAGGGTGTTGGC-3’, reverse 5’-CAGCCCTCTGGTGGGTCAAT-3’.

### Single-cell RNA sequencing

#### Cell isolation for scRNA-seq studies. Rat lung scRNA-seq

Rat lungs were collected after perfusion through the right ventricle with 50 ml ice cold PBS. After mincing with scissors, tissue was suspended in dissociation buffer prepared using Multi Tissue Dissociation kit 2 (Miltenyi, #130-110-203). The suspension was transferred into a gentleMACS C tube (Miltenyi), which was then attached upside down onto the gentleMACS Dissociator (Miltenyi). Program 37C_Multi_C_01 was used to dissociate the lung samples. At the end of the program, digestion reaction was stopped by adding ice cold PBS with 1% FBS and incubated at 37 °C for 15-25 min with gentle trituration by micropipette every 5 min. Sample suspension was filtered through a 70 um MACS SmartStrainer. Cells were pelleted down by centrifuge at 500g for 5 min and suspended in Red Blood Cell Lysis Solution (Miltenyi, #130-094-183) to remove red blood cells and dead cells. Cells were then washed with staining buffer (PBS with 1% FBS) once and stained with BV480-anti-CD31 (BD, #746594), APC-Cy7-anti-Epcam (Biolegend, #324246), and PE-Cy5-anti-CD45 (BD, #559135) antibodies for 25 min at RT, followed with incubation of Live/Dead cell dye for 10min. At the end of staining, cells were washed with staining buffer one more time and kept on ice until fluorescence-activated cell sorting (FACS). **Embryonic scRNA-seq.** Endothelial and stromal cells were separately enriched from the embryos of wildtype C57BL/6 using FACS. Lungs were first micro-dissected and minced with forceps. Samples were pooled into a 15 ml tube with dissociation buffer consisting of 500 U/ml collagenase IV (Worthington #LS004186), 1.2 U/ml dispase (Worthington #LS02100), 32 U/ml DNase I (Worthington #LS002007) in Hanks’ Balanced Salt Solution (Thermo Fisher). The suspension was incubated at 37 °C for 15-25 min with gently trituration by micropipette every 5min. After the incubation, digestion was stopped by topping off the 15ml tube with ice cold PBS with 1% FBS (staining buffer). The cells were then filtered through a 70-μm cell strainer (BD Bioscience). Cells were then washed with staining buffer once and stained with APC-anti-CD31(Biolegend, #102410, 1:400), PE-Cy7-anti-CD326 (Biolegend, #118216, 1:400), APC-Cy7-anti-CD45 or Percp-Cy5.5-anti-CD45 (Biolegend, #103116 or #103132, 1:400), and APC-Cy7-anti-Ter119 (Biolegend, #116223, 1:400) antibodies for 25 min at RT, followed with incubation of Zombie NIR viability dye (Biolegend, #423106, 1:1000) for 10 min. At the end of staining, cells were washed with staining buffer one more time and kept on ice until FACS.

#### Library preparation and sequencing

The dissociated cells were loaded onto a commercially available droplet-based single-cell barcoding system (10x Chromium Controller, 10x Genomics). The Chromium Single Cell 3′ Reagent Kit v3 (10x Genomics) was used to prepare single-cell 3′ barcoded cDNA and Illumina-ready sequencing libraries according to the manufacturer’s instructions. The cDNA libraries were sequenced using an Illumina HiSeq 4000 machine with a mean of ∼76,000 reads per cell.

### Bioinformatic methods

#### Single cell mRNA sequencing read alignment

Using Cell Ranger (version 5.0, 10x Genomics), sequencing reads from single cells isolated using 10x Chromium were demultiplexed and then aligned to custom-built references: Rat Rnor 6.0 (rn6) supplemented with fluorescent gene EGFP, or Mouse GRCm38 (mm10) supplemented with the Cre transgene and fluorescent genes *EGFP* and *tdTomato*.

#### Iterative cell clustering and annotation

Expression profiles of cells from different conditions were clustered together using R software package Seurat (v4.3.3). Preprocessing and normalization were performed as described. Briefly, Unique molecular identifiers (UMIs) were normalized across cells, scaled per 10,000 (10x), and converted to log scale using the ‘NormalizeData’ function. These values were converted to z-scores using the ‘ScaleData’ command and highly variable genes were selected with the ‘FindVariableGenes’ function. Principal components were calculated for these selected genes and then projected onto all other genes with the ‘RunPCA’ command. Clusters of similar cells were detected using the Leiden method for community detection implemented in Seurat.

After separating clusters by expression of tissue compartment markers, we assigned cell types to canonical identities using the most sensitive and specific markers for mouse and human cell types identified in the human lung cell atlas (e.g., ‘Arterial’). We used previously defined markers for developmental annotation of the plexus endothelial progenitors. For subtypes that showed substantial transcriptional change, a representative marker gene was prepended to their presumed canonical identity (e.g., ‘*Dll4*^+^ arterial-like EC’). Marker genes for cluster annotation were selected based on statistical significance (adjusted p-value < 0.05), log fold-change threshold (>0.5), and differential expression in a minimum percentage of cells within the cluster (>25%).

#### Integration analysis and assessment of batch effects

To evaluate whether disease-associated cell populations identified across scRNA-seq datasets reflect biological states rather than technical artifacts, we performed data integration and quantitative assessment of mixing across technical and biological variables for each dataset. Data were partitioned by the dominant source of technical variation (‘processing batch’ or ‘experiment’, defined as biological replicate runs in which multiple samples from different conditions were processed together). This integration strategy balances removal of batch effects while avoiding overcorrection that could artificially merge distinct biological states.

For each partition, expression data were normalized using ‘NormalizeData’, and variable features were identified with ‘FindVariableFeatures’. Integration anchors were computed across partitions using reciprocal PCA (RPCA) with ‘FindIntegrationAnchors’, and batch-corrected embeddings were generated with ‘IntegrateData’. Integrated data were scaled, subjected to principal component analysis (PCA), and visualized using uniform manifold approximation and projection (UMAP).

#### Quantification of integration effectiveness using local neighborhood mixing

To quantitatively distinguish technical variation from biological specificity, we computed the Local Inverse Simpson Index (iLISI) for each cell in the integrated PCA space. iLISI quantifies the effective number of distinct categories present in a cell’s local neighborhood and provides a continuous measure of local diversity. For each cell, a k-nearest-neighbor graph was constructed in integrated PCA space using ‘FindNeighbors’. iLISI values were computed separately for relevant categorical variables including condition (e.g., control vs. disease), processing batch, and experiment or sequencing library. iLISI values approach 1 when a cell’s neighborhood is dominated by a single category and increase with greater local diversity. High batch or experiment iLISI indicates effective removal of technical effects, whereas low condition iLISI coupled with high technical iLISI indicates condition-specific biological enrichment rather than batch-driven clustering. Cell-level iLISI scores were summarized at the cluster level using median and mean values. To further assess potential technical dominance, we calculated the number of contributing experiments and batches, and the maximum fraction of cells contributed by any single technical source within each cluster.

#### Differential gene expression analysis

For cluster annotation, marker genes identifying each cell cluster relative to all other clusters were identified using ‘FindMarkers’ in Seurat with the MAST statistical framework, or using ‘rank_genes_groups’ with the Wilcoxon rank-sum test in Scanpy for initial clustering. Genes were considered cluster markers if they met the following criteria: adjusted p-value < 0.05, log fold-change > 0.5, and expression in ≥25% of cells within the cluster. This approach was used for cell type discovery and annotation within individual datasets. For the mouse developmental dataset, differential expression between Ebf1 knockout and wildtype animals within annotated cell types was performed using ‘FindMarkers’ with the Wilcoxon rank-sum test, appropriate for the isogenic small-cohort experimental design. For cross-condition differential gene expression in the human PAH dataset, a pseudobulk framework was implemented using edgeR (v4.0.16) as described in the Human PAH scRNA-seq analysis section below.

#### Trajectory inference

Cell lineage relationships were reconstructed using Slingshot (v2.0.0), a trajectory inference method designed to identify branching differentiation pathways from single-cell transcriptomic data by first constructing a minimum spanning tree (MST) on cell clusters to identify ordered sets of lineages that share a common starting cluster and lead to distinct terminal clusters, and then assigning pseudotime by extending principal curves to model transcriptional progression along each lineage.

Specifically, we used Seurat-defined clusters and computed UMAP coordinates to obtain a global lineage structure with the ‘getLineages’ function, which implements MST construction and identifies likely lineage transitions between nodes. To estimate transition probabilities, we used the shared nearest-neighbor (SNN) graph constructed in Seurat. Transition strength between clusters was quantified by summing the number of intermediate cells connecting each pair of clusters in the SNN graph. Given an SNN adjacency matrix *A*, where *A*_ij_ represents the connection strength between cells *i* and *j*, we computed a pairwise interaction matrix S, where *S*_mn_ is the transition strength between clusters *C*_m_ and *C*_n_, determined by summing edges in the SNN graph that connect cells from each cluster. Cluster centroids were computed as the mean UMAP coordinates. The final lineage graph was visualized with edge thickness proportional to transition probability, overlaid on the UMAP single cell visualization.

#### Pseudotime gene expression analysis

To identify genes exhibiting dynamic expression changes along pseudotime, we applied TradeSeq (v1.6.0), which models gene expression as a smooth function of pseudotime. The ‘fitGAM’ function was used to fit generalized additive models (GAMs) to individual gene expression profiles, and differential expression testing was performed using ‘associationTest’, which identified genes with significant lineage-dependent expression trends, revealing transcriptional programs associated with developmental processes or disease progression.

#### Receptor-ligand analysis

Receptor-ligand analysis was performed using R package CellChat (v2.1.2) from single cell datasets. Ligand-receptor pairs were inferred using ‘computeCommunProb’ and filtered to keep ligand-receptor interactions involving at least 10 cells per interacting group. To visualize interactions, we generated chord diagrams with ‘netVisual_aggregate (layout = "chord")’.

### Dataset-specific implementations Rat PAH scRNA-seq analysis

#### Quality control and filtering

Raw count matrices were processed using Scanpy (v1.5.1) with Python 3 (numpy v1.18.5, pandas v1.0.5, scipy v1.4.1). Cells were filtered to retain those with ≥500 UMI counts, ≥50 genes detected, and ≤10,000 genes detected to exclude low-quality cells and potential doublets. Mitochondrial gene percentages were calculated and visualized but no explicit threshold was applied. Doublets were identified using Scrublet (v0.2.3) and flagged in metadata. Cells with duplicate barcodes across sequencing libraries were flagged as potential artifacts of index hopping. No ambient RNA correction was performed. Total cells passing QC: 23,075 (11,648 WT, 6,560 silent carrier, 4,867 PAH).

#### Clustering parameters

Data were normalized to 10,000 counts per cell using ‘normalize_total’ and log-transformed with ‘log1p’. Highly variable genes were identified using mean expression thresholds (0.0125-6) and dispersion cutoff (≥0.5). Data were scaled with maximum value 10.

Principal component analysis was computed using the Arpack SVD solver. The first 30 principal components were selected for downstream analysis based on examination of PC loadings and variance explained. Neighborhood graphs were constructed using 30 nearest neighbors. UMAP (v0.4.4) embedding was computed from the neighbor graph. Leiden clustering (leidenalg v0.8.0) was performed at resolution 1.0.

#### Marker gene selection

Marker genes for cluster annotation were identified using ‘rank_genes_groups’ with Wilcoxon rank-sum test (adjusted p-value < 0.05, log fold-change > 0.5, minimum expression in 25% of cells within cluster). Top differentially expressed genes were used in combination with canonical cell type markers from published mouse and human lung atlases.

#### Endothelial integration

Endothelial cells were subsetted based on curated annotations including arterial, venous, general capillary (gCap), aerocyte (aCap), lymphatic, proliferative gCap (gCap-p), *Dll4*^+^ endothelial, and *Ebf1*^+^ endothelial populations (n = 16,332 cells). Integration was performed across experiments (3 experiments representing biological replicate processing runs) using Seurat (v4.3.3) in R. For each experiment, 3,000 variable features were selected using ‘FindVariableFeatures’. Integration anchors were computed using reciprocal PCA (RPCA) with ‘FindIntegrationAnchors’, and batch-corrected embeddings were generated with ‘IntegrateData’. Integrated data were scaled, subjected to principal component analysis (PCA) using the first 30 dimensions, and visualized using UMAP (**Extended Data Figure 1e-g**). Integration quality was assessed using the inverse Simpson’s Local Inverse Simpson Index (iLISI), which quantifies the effective diversity of cell sources in local k-nearest neighbor graphs (k = 50). iLISI was computed for three variables: condition (wild-type, silent carrier, PAH), experiment, and processing batch (sequencing library/lane). iLISI values near 1 indicate strong separation by the variable of interest, while higher values indicate effective mixing. This analysis confirmed that canonical endothelial populations (gCap, aCap, arterial, venous, lymphatic) showed robust condition mixing (median condition iLISI: 1.37-2.02) and excellent batch mixing (median batch iLISI: 1.69-3.49), while disease-enriched populations (*Ebf1*^+^ ECs, gCap-p) maintained condition-specific neighborhoods (median condition iLISI = 1.0) despite spanning multiple experiments (3 experiments each) and processing batches (4 and 6 batches, respectively), supporting biological specificity rather than technical artifacts. *Dll4^+^* ECs showed condition-enrichment (median condition iLISI = 1.86) with excellent batch mixing (median batch iLISI = 2.19) across 2 processing batches. The intermediate iLISI-condition value, higher than EBF1^+^ ECs and gCap-p (iLISI = 1.0) but lower than canonical endothelial populations (iLISI = 1.45-2.02), indicates these cells occupy neighborhoods with greater condition diversity than the purely disease-specific populations, consistent with an arterial-like state that shows disease-associated enrichment. Complete integration quality metrics are provided in **Supplementary Table 5**.

#### Trajectory analysis

A neointimal cluster of endothelial cells from PAH condition were subsetted for trajectory analysis and re-processed (n = 2,125 cells; including arterial, venous, gCap, aCap, gCap-p, Dll4+ EC, and EBF1+ EC populations). Cells were normalized using SCTransform with 3,000 variable features, scaled (maximum value 10), and PCA computed. The first 18 principal components were used for neighbor graph construction (*k* = 15) and UMAP embedding. Leiden clustering was performed at resolution 1.0.

Trajectory inference was performed using Slingshot (v2.0.0). Lineages were inferred from UMAP coordinates using ‘getLineages’ with general capillary cells designated as the start population. Principal curves were fitted using ‘getCurves’. Multiple lineages were identified; two major trajectories were selected for detailed analysis representing (1) general capillary to *Dll4*^+^ arterial-like differentiation and (2) general capillary to *EBF1*^+^ neointimal transformation. Pseudotime values were extracted for each lineage and scaled 0-1. To separate the two trajectories emanating from the shared root population, general capillary cells were randomly split 50/50 (seed = 42) and assigned to either the arterial or EBF1 lineage based on trajectory membership.

Transition probabilities between cell populations were estimated using the shared nearest-neighbor (SNN) graph constructed in Seurat. Pairwise interaction strength was calculated by summing edges connecting cells between each population pair. The minimum spanning tree from Slingshot was used to identify biologically relevant transitions, and weak interactions were filtered for visualization.

Differential expression along pseudotime was performed using TradeSeq (v1.6.0). Gene counts were filtered to retain genes with >3 counts in >1% of cells. Generalized additive models were fitted using ‘fitGAM’. Three complementary tests were performed: (1) ‘diffEndTest’ to identify genes differentially expressed at trajectory endpoints between lineages, (2) ‘associationTest’ to identify genes with significant expression changes along pseudotime within each lineage, and (3) pattern testing to identify genes with divergent expression patterns between lineages. P-values were adjusted for multiple testing using the false discovery rate (FDR) method. Genes with FDR < 0.05 were considered significant.

### Human PAH scRNA-seq analysis

#### Quality control and filtering

Raw count matrices were processed using the Seurat R package (v5.1.0). Cells were filtered to retain those with ≥200 genes detected and sample-specific upper thresholds ranging from 1,000 to 7,000 genes detected to exclude low-quality cells and potential doublets. Cells with mitochondrial gene content >10% were excluded. Doublets were identified using scDblFinder (v1.16.0) and removed for each sample. Data were normalized, variable features were identified per sample, and datasets were integrated using canonical correlation analysis (CCA) as described above. No ambient RNA correction was performed. Principal component analysis was computed, and the first 30 principal components were selected for downstream analysis. Neighborhood graphs were constructed and UMAP embedding was computed from the neighbor graph. Graph-based clustering was performed using the Leiden method at resolution 0.5. Total cells: 55,111 endothelial cells from 40 patients across 44 sequencing libraries. Distribution by disease condition: Control (n = 12,351 cells, 10 patients), DTPAH (n = 10,150 cells, 10 patients), IPAH (n = 9,826 cells, 10 patients), SPSPAH (n = 22,784 cells, 10 patients).

#### Endothelial integration

Human pulmonary arterial hypertension endothelial cells (n = 55,111 cells) from 40 patients across 44 sequencing libraries were integrated using Seurat’s CCA-based integration workflow. Briefly, each sample was log-normalized individually, variable features were identified per sample, and the top 2,000 shared variable features were used to identify integration anchors via canonical correlation analysis (CCA). The datasets were then integrated using these anchors, and all downstream analyses — including PCA, neighbor finding, clustering, and UMAP — were performed in the integrated assay space to pool signal across genetically heterogeneous patients with variable disease severity and diverse etiologies, enabling identification of conserved gene expression patterns.

Integration quality was assessed post-hoc using the inverse Simpson’s Local Inverse Simpson Index (iLISI), which quantifies the effective diversity of cell sources in local k-nearest neighbor graphs (k = 50). iLISI was computed for three variables: disease condition (Control, DTPAH [drug/toxin-associated PAH], IPAH [idiopathic PAH], SPSPAH [systemic and pulmonary shunt associated PAH]), patient identity, and sequencing library. iLISI values near 1 indicate strong separation by the variable of interest, while higher values indicate effective mixing. This analysis confirmed that all seven endothelial cell types showed robust patient mixing (median patient iLISI: 10-16) and excellent batch mixing (median batch iLISI: 10-16), with all populations spanning all 40 patients and 44 sequencing libraries. EBF1^+^CAP cells showed the lowest condition iLISI (2.72) compared to other endothelial populations (SVEC: 2.76, VEC: 3.16, SAEC: 3.23, gCap: 3.28, aCap: 3.33, AEC: 3.49), indicating disease-associated enrichment (60.8% SPSPAH, 16.7% DTPAH, 16.5% IPAH, 6% Control) while maintaining presence across multiple disease subtypes and controls. The intermediate condition iLISI value for EBF1^+^CAP cells, lower than canonical endothelial populations but substantially higher than the disease-exclusive populations observed in the controlled rat model (iLISI = 1.0), likely reflects biological heterogeneity across human disease etiologies and genetic backgrounds. Complete integration quality metrics are provided in **Supplementary Table 5**.

#### Differential gene expression analysis

For cross-condition differential gene expression, a pseudobulk framework was implemented using edgeR (v4.0.16). Raw counts were aggregated per patient per annotated cell type to generate pseudobulk count profiles. Pseudobulk samples with fewer than 20 cells were excluded. For each cell type, a design matrix was constructed with disease condition as the primary variable of interest, with Control as the reference level. Library sizes were normalized using edgeR’s trimmed mean of M-values (TMM) method with ‘calcNormFactors’, and genes were filtered using ‘filterByExpr’ to retain only those with sufficient counts across samples. Dispersions were estimated using ‘estimateDisp’ and model fitting was performed using the quasi-likelihood framework with ‘glmQLFit’ and ‘glmQLFTest’.

Two complementary analyses were performed: a binary comparison (PAH versus Control, collapsing DTPAH, IPAH, and SPSPAH into a single PAH group) and a subtype-resolved analysis comparing each PAH subtype (DTPAH, IPAH, SPSPAH) independently to Control, with an additional global likelihood ratio test across all condition terms. P-values were adjusted for multiple testing using the Benjamini-Hochberg false discovery rate method within each cell type and contrast.

#### Trajectory analysis

Trajectory inference was performed on the cleaned endothelial dataset (n = 55,111 cells) using Slingshot (v2.0.0) on integrated UMAP coordinates. Lineages were inferred using ‘getLineages’ with gCap (cluster 0) designated as the start population. Principal curves were fitted using ‘getCurves’, identifying 7 distinct trajectories. Three major lineages were selected for detailed analysis representing (1) CAP to EBF1+ neointimal transformation, (2) CAP to arterial differentiation (Lineage 4), and (3) CAP to neo-arterialization. Cells assigned to multiple lineages were randomly split to ensure mutually exclusive lineage membership (seed = 42-45). Pseudotime values were extracted for each lineage and scaled 0-1. A merged pseudotime axis was created by coalescing lineage-specific pseudotimes using the first non-NA value per cell. Transition probabilities between cell populations were estimated using the shared nearest-neighbor (SNN) graph (k=20 neighbors, 30 dimensions). Pairwise interaction strength was calculated by summing edges connecting cells between each population pair. The minimum spanning tree from Slingshot was used to identify biologically relevant transitions.

Differential expression was performed using TradeSeq (v1.6.0) on Lineages 3, 4, 5, and 6. Gene counts were filtered to retain genes with >3 counts in >1% of cells. Generalized additive models were fitted using ‘fitGAM’ with 6 knots. Two complementary tests were performed: (1) ‘diffEndTest’ (global and pairwise) to identify genes differentially expressed at trajectory endpoints between lineages, and (2) ‘associationTest’ to identify genes with significant expression changes along pseudotime within each lineage. P-values were adjusted for multiple testing using the Benjamini-Hochberg false discovery rate (FDR) method.

#### Statistical testing of disease association

Lineage composition and pseudotime progression differences between disease states were assessed using per-sample Wilcoxon rank-sum tests (PAH vs Control) and Kruskal-Wallis tests for multi-group comparisons (Control, DTPAH, IPAH, SPSPAH). Per-sample lineage fractions and median pseudotime values were computed for each patient across all lineages. P-values were adjusted using the Benjamini-Hochberg method. Cell type compositions were broadly comparable between control and disease, with the exception of aCaps showing marked reduction in disease (29.6% of control ECs vs 15.8-17.7% in PAH, χ² residual +30.4).

#### CellChat analysis and network visualization

CellChat (v2.1.2) was performed on human PAH and control samples from the Dai et al. dataset. Communication probabilities were calculated for all vascular cell type pairs using ‘computeCommunProb’ (‘triMean’ method) and filtered to retain interactions with ≥10 cells per group. PAH-enriched interactions were defined as PAH-only or shared interactions with positive delta probability (PAH - Control).

For network visualization, interactions between vascular cell types meeting stringent thresholds were displayed: communication probability >0.3 in PAH or Δprob >0.1. This filtering identified 23 edges between six cell types (EBF1^+^ EC, CXCL12^+^ EC, SVEC, AEC, ASMC, AF2). Nodes were positioned with EBF1^+^ EC centered and partners arranged circularly. Edge width was scaled proportionally to PAH communication. Edge color indicates signaling direction: red (EBF1+CAP sending), blue (EBF1^+^CAP receiving), gray (other). Line style encodes PAH enrichment: solid (>1.5-fold increase vs. control), dashed (<1.5-fold). Send/receive (S/R) ratios were calculated as total outgoing divided by incoming communication probability.

### Mouse developmental lung scRNA-seq analysis

#### Quality control and filtering

Raw count matrices were processed using Scanpy (v1.5.1) with Python 3 (numpy v1.18.5, pandas v1.0.5, scipy v1.4.1). Cells were filtered to retain those with ≥500 UMI counts and ≥50 genes detected. Cells with >10,000 genes detected were excluded as potential doublets. Mitochondrial gene percentages were calculated using genes with names prefixed ‘mt-’, and cells with >40% mitochondrial counts were excluded. No doublet detection or ambient RNA correction was performed. After quality control, 26,563 cells were retained for analysis (21,222 endothelial, 5,341 stromal).

#### Clustering parameters

Data were normalized to 10,000 counts per cell using ‘normalize_total’ and log-transformed with ‘log1p’. Highly variable genes were identified using mean expression thresholds (0.0125–6) and a dispersion cutoff (≥0.5). Data were scaled with maximum value 10. Principal component analysis was computed using the Arpack SVD solver. The first 30 principal components were selected for downstream analysis based on examination of PC loadings and variance explained. Neighborhood graphs were constructed using 30 nearest neighbors. UMAP embedding was computed from the neighbor graph. Leiden clustering was performed at resolution 1.0.

#### Endothelial integration

Endothelial and stromal cells from embryonic lungs (E11.5-E15.5, n=26,563 cells) were integrated across 7 sequencing batches using Seurat’s reciprocal PCA (RPCA) method. Briefly, data were split by batch, each batch was log-normalized individually and variable features identified (3,000 features per batch), and integration anchors were identified using RPCA. Datasets were integrated using these anchors, and all downstream analyses were performed in the integrated assay space. Principal component analysis was computed, and the first 30 principal components were used for neighbor graph construction and UMAP embedding.

Integration quality was assessed using iLISI as described above, computed for developmental stage (E11.5, E12.5, E13.5, E15.5) and sequencing batch. This analysis confirmed that all populations showed robust cross-batch mixing (median batch iLISI: 2.0-3.48), with detection across multiple independent batches (**Supplementary Table 5**). Stage-specific progenitor populations maintained their developmental specificity after integration (median age iLISI: 1.27-2.31) while showing excellent batch mixing, confirming these represent genuine developmental populations rather than batch artifacts. For example, alveolar fibroblast progenitors (n=1,284) showed strong E13.5 enrichment (99.7%) with detection across 4 batches (median batch iLISI = 3.09), while Aplnr+ plexus progenitors (n=19,126, 72% of endothelial cells) spanned all developmental stages with detection across all 6 batches (median batch iLISI = 2.94). Ebf1+ endothelial cells showed developmental progression from E13.5 progenitors (54%) to E15.5 differentiated cells (46%) with detection across 4 batches. All stage-specific populations demonstrated low age iLISI (biological stage enrichment) combined with high batch iLISI (successful integration), confirming batch effects were effectively removed while preserving developmental biology. Complete integration quality metrics are provided in **Supplementary Table 5**.

#### Genotype-independent cell type validation

To validate that cell type assignments were robust and not confounded by Ebf1 genotype, we performed integration analysis on the complete dataset including both wildtype and Ebf1 knockout embryos (n=45,531 cells, 7 batches, 2 genotypes: WT and *Cdh5*-Cre;*Rosa26*^tdTomato^*;Ebf1^fl^*^/fl^. Cell types were annotated identically across genotypes using the same marker-based classification approach (**Extended Data Fig. 6e-h**). Integration quality assessment using iLISI confirmed robust cross-batch mixing (median batch iLISI: 1.54-4.42) and genotype mixing (median genotype iLISI: 1.42-1.81) for all cell types except Ebf1^+^ endothelial cells, which showed complete genotype separation (genotype iLISI = 1.0) with 90.1% wildtype and 9.9% knockout representation. This pattern validated that: (1) cell type annotations are biologically accurate and reproducible independent of Ebf1 status, (2) integration successfully removed technical batch effects while preserving biological differences, (3) Ebf1^+^ endothelial cells represent a bona fide Ebf1-dependent population rather than a batch artifact, and (4) Ebf1 knockout specifically affects the Ebf1^+^ EC population in a cell-autonomous manner without disrupting the cell specification of other cell types. Complete integration quality metrics for both wildtype-only and full dataset analyses are provided in **Supplementary Tables 5**.

#### Differential gene expression analysis

Differential expression between Ebf1 knockout and wildtype animals within annotated cell types was performed using ‘FindMarkers’ with the Wilcoxon rank-sum test, applied separately for early deletion (E12.5) and late deletion (E15.5) timepoints on the isogenic dataset.

#### Trajectory analysis

Endothelial cells were subsetted to populations participating in early arterial specification from E12.5-E13.5 developmental stages (n=2,125 cells including plexus, vein, artery, and Ebf1^+^ endothelial populations). Trajectory inference was performed using Slingshot (v2.0.0) on UMAP coordinates. Lineages were inferred using ‘getLineages’. Principal curves were fitted using ‘getCurves’, identifying multiple trajectories. Two major lineages were selected for detailed analysis representing (1) plexus to arterial differentiation and (2) plexus to Ebf1^+^ endothelial transformation. To separate trajectories emanating from the shared root population, plexus cells (cluster 0) were randomly split 50/50 (seed=42) and assigned to either the arterial or Ebf1 lineage based on trajectory membership. Pseudotime values were extracted for each lineage and scaled 0-1. A merged pseudotime axis was created by coalescing lineage-specific pseudotimes using the first non-NA value per cell.

Transition probabilities between cell populations were estimated using the shared nearest-neighbor (SNN) graph constructed in Seurat. Pairwise interaction strength was calculated by summing edges connecting cells between each population pair. The minimum spanning tree from Slingshot was used to identify biologically relevant transitions.

Differential expression along pseudotime was performed using TradeSeq (v1.6.0). Gene counts were filtered to retain genes with >3 counts in >1% of cells. Generalized additive models were fitted using ‘fitGAM’. Two complementary tests were performed: (1) ‘diffEndTest’ (global and pairwise) to identify genes differentially expressed at trajectory endpoints between lineages, and (2) ‘associationTest’ to identify genes with significant expression changes along pseudotime within each lineage. P-values were adjusted for multiple testing using the Benjamini-Hochberg false discovery rate (FDR) method.

### Statistical Analysis

No statistical methods were used to predetermine sample size. Sample sizes were selected based on prior experience with these models and are reported in each figure legend. Data collection and quantification were performed blinded to genotype and/or treatment where indicated. Unless otherwise stated, *n* denotes the number of biologically independent samples (animals, human donors, or independent cell culture experiments). Technical replicates (for example, replicate wells or repeated measurements from the same sample) are indicated in the figure legends where applicable. For experiments with hierarchical sampling (for example, multiple sections, vessels, or ROIs per specimen), repeated measurements were averaged to obtain a single value per animal/donor, and statistical testing was performed on these per-animal/per-donor values to avoid pseudo-replication. Data are presented as mean ± s.e.m., with individual data points shown, unless otherwise stated. Two-group comparisons were performed using two-sided Mann-Whitney tests. For scRNA-seq analyses, animals/donors were treated as independent biological replicates; cell-level measurements were not treated as independent observations. Statistical analyses were performed using GraphPad Prism (v10). Exact tests, *n*, and *P* values are reported in the corresponding figure legends.

## Data Availability

Raw sequencing data, UMI tables, cellular annotation metadata, Seurat objects, and scanpy objects are being deposited and will be released as soon as possible (at latest, upon acceptance of this manuscript).

## Code Availability

Code to reproduce the analyses and figures is being deposited to Github and will be released as soon as possible (at latest, upon acceptance of this manuscript).

## Extended Data

**Extended Data Fig. 1 Single cell RNA-seq analyses of rat PAH vascular lesions. a,** Experimental schematic. Lung endothelial (CD31^+^), stromal (Epcam^-^CD31^-^CD45^-^), epithelial (Epcam^+^), and immune (CD45^+^) populations were enriched by FACS from wildtype (WT), heterozygous control (*Bmpr2*^+/-^ mutant), or PAH (*Bmpr2*^+/-^ mutant with intratracheal instillation of AdALOX5 to induce disease) rats in a previously published animal model of PAH^14^. n=3 per animal group. Single cell libraries were generated from the endothelial and stromal fractions. **b**-**d**, UMAP visualization of integrated vascular cell clusters. Clusters are color-coded by cell-type in **b**, by condition (WT, heterozygous control, PAH) in **c**, and by sequencing library (sample ID) in **d**. **e-g**, UMAP projections of endothelial cells integrated by experimental batch, color-coded by annotated cell type in **e**, by condition in **f**, and by sequencing library in **g**. Effective mixing across sequencing libraries with preservation of disease-enriched endothelial states (*Ebf1^+^*ECs, gCap-p, *Dll4*^+^ ECs) confirms successful integration with retention of biological specificity. For quantitative assessment of integration quality, see **Supplementary Table 5** and Methods. **h**, Heatmap of Pearson correlation coefficients of mean gene expression profiles across vascular cell clusters, based on normalized average expression. Proliferating subsets (Fib-p, gCap-p) were excluded. Rows and columns are clustered by correlation distance. *Bmpr2,* bone morphogenetic protein receptor 2.

**Extended Data Fig. 2 EBF1^+^ ECs are broadly induced across animal models of PAH. a, b,** Representative immunofluorescence images of PAH lungs from two well-established rat models of disease: athymic plus SU5416 model^30^ (**a)** and monocrotaline model^29^ (**b**). Insets highlight PAs with neointimal lesions. 3D renderings were generated in Imaris. Dotted outlines mark clusters of EBF1^+^ERG^+^ ECs within neointima. n=8 rats per model in **a**, **b**. **c, d,** Representative single molecule fluorescent *in situ* hybridization (smFISH) combined with immunofluorescence images of PAH lungs from two well-established mouse models of disease: house-dust-mite-induced model^32^ (**c**) and lung-specific TNFα overexpression model^31^ (**d**). Insets highlight PAs with neointimal lesions. Dotted outlines mark clusters of *Ebf1*^+^*Cldn5*^+^ ECs (**c**) or *Ebf1*^+^ERG^+^ ECs (**d**). n=3 mice per model in **c**, **d**. DAPI (blue) marks nuclei. Scale bars, 15μm in **a**, **b** and 40μm in **c**, **d**.

**Extended Data Fig. 3 EBF1^+^ ECs are a rich source of developmental vasculotrophic signals associated with PAH pathogenesis**. Chord diagrams showing ligand-receptor interactions inferred by CellChat from rat PAH scRNA-seq data, separated into endothelial-to-endothelial (left) and endothelial-to-stromal (right) signaling. Each diagram shows a distinct signaling pathway: VCAM, Apelin/APJ, CXCL12/CXCR4, and Notch (endothelial-to-endothelial); Laminin, TGF-β, Semaphorin, and Prostaglandin (endothelial-to-stromal). Chord width is proportional to communication probability. EBF1^+^ EC and *Dll4*^+^ EC labels are highlighted in red and blue respectively.

**Extended Data Fig. 4 Human PAH features disease-associated EBF1^+^ ECs and *CXCL12*^high^ neo-arterial ECs. a**, UMAP visualization of annotated endothelial cell clusters from human control and PAH lungs (n=40 patients, 55,111 cells). **b**, Dot plot of representative marker genes across clusters (fraction expressing and mean expression among expressing cells). **c**-**e**, UMAP projections of integrated endothelial cells color-coded by disease condition (Control, DTPAH [drug- and toxin-induced PAH], IPAH [idiopathic PAH], SPSPAH [systemic and pulmonary shunt-associated PAH]) in **c**, by sample identity in **d**, and by sequencing library in **e**. Effective mixing across patients and sequencing libraries with preservation of disease-enriched populations confirms successful integration with retention of biological specificity (see Methods and **Supplementary Table 5**).

**Extended Data Fig. 5 PAH-associated signals regulate EBF1 expression in microvascular ECs. a, b,** mRNA expression of *EBF1* in human pulmonary microvascular endothelial cell (hMVEC-L) cultures following stimulation with factors implicated in PAH pathogenesis. Cells were treated for 16-18 hours with TGF-β pathway ligands (BMP9, TGFβ, Activin A, or GDF11) or pro-angiogenic molecules (IGF1, VEGF-A, or CXCL12). *EBF1* mRNA was quantified by qPCR and compared to vehicle-treated controls. n=6. Data are presented as mean ± s.e.m. with individual data points shown. Statistical comparisons were performed using two-sided Mann-Whitney tests; p values are indicated on the graphs.

**Extended Data Fig. 6 scRNA-seq analyses of the developing mouse lung vasculature (E11.5-E15.5). a**-**d**, UMAP visualization of RPCA-integrated endothelial and stromal cells from wildtype embryos across developmental stages (E11.5-E15.5), color-coded by annotated cell type in **a**, Ebf1 expression in **b**, developmental stage in **c**, and sequencing library in **d**. Effective mixing across sequencing libraries with preservation of stage-enriched populations confirms successful integration. **e**-**h**, UMAP visualization of RPCA-integrated endothelial and stromal cells from both wildtype and *Cdh5*-CreERT2;*Ebf1*^fl/fl^ KO embryos, color-coded by annotated cell type in **e**, genotype in **f**, developmental stage in **g**, and sequencing library in **h**. Effective mixing of WT and KO cells across all shared populations, with selective depletion of the *Ebf1^+^* EC cluster in KO embryos, confirms successful integration with retention of genotype-specific biological differences. For quantitative assessment of integration quality, see **Supplementary Table 5** and Methods.

**Extended Data Fig. 7 Pulmonary vasculature arises from *Aplnr^+^* lung plexus progenitors. a,** Loess-smoothed expression (*y* axis) of the indicated genes along the pseudotime trajectories of the Ebf1 (blue) and the arterialization programs (red) in **Fig. 3f. b,** Experimental schematic and representative whole-mount immunofluorescence image of an *Aplnr*-CreERT2;*Rosa26*^tdTomato^ lineage-traced embryo at E10.5 following pulse labeling induced by 4-hydroxytamoxifen (4OHT) at E8.5. n=6. Cartoon schematic (right) illustrates *Aplnr*-lineage-positive endothelial progenitors within the heart-lung complex. OFT, cardiac outflow tract. IFT, cardiac inflow tract. AO, aorta. **c-f,** Representative whole-mount immunofluorescence images of *Aplnr*-CreERT2;*Rosa26*^tdTomato^ lineage-traced embryos collected at E12 (**c**), E12.5 (**d**), E13.5 (**e**), or E14.5 (**f**) following pulse labeling with 4OHT at E11.5. n=6 for **c**, **d**, **f**; n=8 for **e.** Yellow dotted outlines mark the proximal PAs. L1 branch, the first PA branch in the left lung. **g,** Experimental schematic and representative whole-mount immunofluorescence image of a *Cdh5-*CreERT2*;Rosa26*^tdTomato^ lineage-traced embryo at E14.5 with lineage labeling initiated by tamoxifen (TAM) at E8.5. n=6. Scale bars, 100 μm in **b** and 200 μm in **c-g.**

**Extended Data Fig. 8 EBF1*^+^* ECs comprise a subset of Aplnr-lineage lung plexus cells. a,** Experimental schematic for **b-d**. **b**-**d**, Representative whole-mount immunofluorescence images of *Aplnr*-CreERT2;*Rosa26*^tdTomato^ lineage-traced embryos following pulse labeling with 4OHT at E11.5, collected at E12.5 (**b**), E13.5 (**c**), or E14.5 (**d**). n=8 for **b**, **c**; n=6 for **d**. Panel **a** cartoon illustrates the distribution of EBF1^+^*Aplnr*-lineage-positive ECs within the lung plexus. Scale bar, 200μm. **e**, Representative whole-mount immunofluorescence image of a *Gja5-* CreERT2*;Rosa26*^tdTomato^ lineage-traced embryo at E13.5, showing minimal overlap between EBF1 antibody signals and *Gja5-*CreERT2 lineage-labeled mature arterial and pre-arterial ECs. Pulse labeling of *Gja5-*CreERT2 was initiated at E11.5 by 4OHT. Blue arrowheads indicate EBF1^+^*Gja5*-lineage-negative cells. Magenta arrowheads indicate rare EBF1^+^*Gja5*-lineage-positive cells in the proximal PA. Blue dotted outlines mark PA branches. n=6. Scale bars, 20μm in the proximal PA panel and 30μm in the L1 branch and plexus panel.

**Extended Data Fig. 9 Distribution of Ebf1^lin^ cells during the embryonic stage of lung development. a,** Schematic of the CRISPR/Cas9-mediated Ebf1-P2A-CreERT2 (*Ebf1-*CreERT2) knock-in allele, in which CreERT2 is inserted in-frame at the endogenous Ebf1 stop codon via homology-directed repair. **b,** Experimental designs for **c**-**e**. Pulse labeling was initiated by 4OHT at E8.5. **c**-**e,** Representative whole-mount immunofluorescence images of *Ebf1*-CreERT2; *Rosa26*^tdTomato^ lineage-traced embryos following pulse labeling with 4OHT at E8.5, collected at E9.5 (**c**), E10.5 (**d**), or E11.5 (**e**). n=8 for **c** and **d**; n=6 for **e**. Cartoons in **d** and **e** illustrate the locations and possible behaviors of *Ebf1-*CreERT2 lineage-labeled endothelial cells (*Ebf1^lin^* ECs). OFT, cardiac outflow tract. IFT, cardiac inflow tract. PV, pulmonary vein. PA, pulmonary artery. Scale bars, 200 μm (left panel) and 150 μm (right panel) in **c**, 300 μm (left panel) and 150 μm (right panel) in **d**, and 20 μm in **e**.

**Extended Data Fig.10 Ebf1^lin^ ECs show minimal incorporation into developing PA endothelium. a,** Experimental schematic for **b-f**. **b,** Representative whole-mount immunofluorescence images of *Ebf1*-CreERT2;*Rosa26*^tdTomato^ lineage-traced lungs collected at E13.5 showing the plexus area. **c**, Representative whole-mount immunofluorescence image of an *Ebf1*-CreERT2;*Rosa26*^tdTomato^ lineage-traced embryo at E13.5. Schematics indicate the enrichment of *Ebf1^lin^*ECs around the developing PA branches. *Ebf1*^lin^ ECs were defined as tdTomato lineage-labeled cells visualized as a mask over the ERG antibody channel in Imaris. **d**-**f**, Representative whole-mount immunofluorescence images of *Ebf1*-CreERT2;*Rosa26*^tdTomato^ lineage-traced lungs following pulse labeling with 4OHT at E11.5, collected at E12.5 (**d**) or E13.5 (**e**, **f**). 3D renderings and orthogonal sectional views highlight the relative intraluminal (yellow arrowheads) or abluminal (blue arrowheads) localization of *Ebf1^lin^* cells. n=8 for **b-e**, n=4 for **f**. Scale bars, 20 μm in **b**, **f**; 150 μm in **c**; and 100 μm in **d**, **e**.

**Extended Data Fig. 11 *Ebf1*^lin^ ECs are transiently proliferative and migratory with rich vasculotrophic signals. a,** Experimental schematic for **b, c**. **b,** Time-lapse imaging of an E12.5 *Ebf1*-CreERT2;*Rosa26*^tdTomato^ lung explant over 48 hours. Insets show Ebf1^lin^ cells (tdTomato^+^) in the lung plexus at each time point. n=6. **c,** Representative single-cell tracking trace showing cell generation number over assay time (seconds), demonstrating clonal expansion of a single *Ebf1*^lin^ cell through four generations over 48 hours. **d,** Experimental schematic for **e, f**. **e,** Representative smFISH and immunofluorescence images of lung sections from healthy *Ebf1*-CreERT2;*Rosa26*^tdTomato^ mice showing rare *Ebf1*^lin^ ECs (tdTomato^+^ERG^+^ *Ebf1^+^Pecam1^+^*) within the pulmonary microvasculature. Dotted outlines mark both an *Ebf1*^lin^ EC (tdTomato^+^ERG^+^ *Ebf1^+^Pecam1^+^*) and an *Ebf1*^lin^ stromal cell (tdTomato^+^ERG^-^aSMA^+^*Ebf1^+^Pecam1^-^*). n=5. Quantification shown in **Extended Data Fig. 14f** (Ctr group). **f**, Representative whole-mount immunofluorescence image of healthy *Ebf1*-CreERT2;*Rosa26*^tdTomato^ adult lung showing rare *Ebf1*^lin^ ECs and their relationship to Ki67^+^ERG^+^ proliferating ECs. n=5. Quantification shown in **Extended Data Fig. 14s** (control group). **g,** Chord diagrams showing ligand-receptor interactions inferred by CellChat from the developing lung scRNA-seq data (**Fig. 3** and **Extended Data Fig. 6a-d**), separated into endothelial-to-endothelial (top) and endothelial-to-stromal (bottom) signaling. Chord width is proportional to communication probability. *Ebf1*^+^ EC labels are highlighted in red. Scale bars, 100 μm in **b**, 20 μm in **e**, and 25 μm in **f**.

**Extended Data Fig. 12 Plexus-specific *Ebf1* is essential for PA development. a,** Experimental schematic for **b**, **c** (pan-endothelial KO) with recombination initiated by 4OHT at E8.5 (embryonic stage of lung development). **b,** Representative brightfield and fluorescent images of the lung-heart complex from *Cdh5^Cre-^*;*Ebf1^fl/fl^* (Cre-negative control, n=12), *Cdh5^CreERT2^*;*Rosa26*^tdTomato^;*Ebf1^fl/wt^* (EC-Ebf1^fl/w*t*^ Ctr, n=12), and *Cdh5^CreERT2^*;*Rosa26*^tdTomato^; *Ebf1^fl/fl^* (EC-Ebf1 KO, n=12) embryos collected at E13.5. Lung size is comparable between Cre-negative control and EC-Ebf1^fl/wt^ Ctr, whereas EC-Ebf1 KO embryos show reduced lung size and delayed development. Heart size is similar across genotypes. **c,** Representative whole-mount immunofluorescence images of Cre-negative control and EC-Ebf1 control lungs at E13.5 (n=12 per group). Yellow arrowheads indicate SMA coverage in proximal PAs; pink arrowheads indicate PA branching. Quantification of proximal PA diameter and SMA coverage is shown to the right. Data are presented as mean ± s.e.m. with individual data points shown; statistical comparisons were performed using two-sided Mann-Whitney tests; p values are indicated on the graphs. **d,** Experimental schematic (plexus-restricted KO) and representative brightfield image of *Aplnr-*CreERT2;*Ebf1*^fl/wt^ (Plexus-Ebf1^fl/wt^ Ctr, n=16) and *Aplnr-*CreERT2;*Ebf1*^fl/fl^ (Plexus-Ebf1 KO, n=17) lungs collected at E13.5. **e,** Experimental schematic and representative brightfield and immunofluorescence images of *Gja5-*CreERT2;*Rosa26*^tdTomato^;*Ebf1*^fl/fl^ (Artery-Ebf1 KO) embryos collected at E13.5. n=4. **f,** Experimental schematic and representative brightfield and immunofluorescence images of *Pdgfra-*CreERT2;*Rosa26*^tdTomato^;*Ebf1*^fl/fl^ (Stromal-Ebf1 KO) embryos collected at E13.5. n=4. **g**, Volcano plot of differential gene expression in the plexus population comparing EC-Ebf1 KO versus Cre-negative littermates. Endothelial and stromal compartments were enriched for scRNA-seq. n=8 per genotype. **h,** Representative brightfield images of *Aplnr-*CreERT2*;H11*^+/+^ (Plexus-OE control, n=10) and *Aplnr-*CreERT2*;H11*^LSL-Ebf1^ (Plexus-Ebf1 OE, n=12) lungs collected at E13.5. **i**, Experimental schematic and representative brightfield and immunofluorescence images of *Gja5-*CreERT2;*Rosa26*^tdTomato^;*H11*^LSL-Ebf1^ (Artery-Ebf1 OE) embryos collected at E13.5. n=4. **j,** Experimental schematic for **k-m** (pan-endothelial KO) with recombination induced by 4OHT at E11.5 (pseudoglandular stage of lung development). **k,** Representative brightfield images of *Cdh5-*CreERT2;*Ebf1*^fl/wt^ (EC-Ebf1^fl/wt^ control, n=6) and *Cdh5-*CreERT2;*Ebf1*^fl/fl^ (EC-Ebf1 KO, n=6) lungs at the time of collection (E15.5). Black arrowheads mark areas of hemorrhage. **l,** Representative whole-mount immunofluorescence of *Cdh5-*CreERT2;*Rosa26*^tdTomato^;*Ebf1*^fl/wt^(EC-Ebf1^fl/wt^ control, n=6) and *Cdh5-*CreERT2;*Rosa26*^tdTomato^;*Ebf1*^fl/fl^ (EC-Ebf1 KO, n=6) lungs at E15.5. Yellow dotted outlines mark distal PAs and branches. Yellow arrowheads indicate reduced PA branching and aSMA coverage in EC-Ebf1 KO embryos. **m**, Volcano plot of differential gene expression in stromal populations comparing EC-Ebf1 KO versus Cre*-*negative littermates. n=6 per genotype. **n**, Schematic summary of pulmonary vascular phenotypes following early endothelial-specific deletion or overexpression of *Ebf1*. Scale bars, 200 μm in **c**, 100 μm in **e**, **f**, **i**, and 50 μm in **l**.

**Extended Data Fig. 13 Ebf1^+^ ECs promote angiogenesis of neighboring Ebf1-negative cells. a,** Experimental schematic. Human pulmonary microvascular endothelial cells were transduced with an adenovirus overexpressing *EBF1* (Ad*EBF1*, MOI=2.5) to generate Ebf1^OE^ ECs. Control cells without EBF1 overexpression expressed minimal endogenous EBF1 and are denoted as Ebf1^neg^ ECs. For tube formation assays, 9,500 Ebf1^neg^ ECs were cocultured with 500 Ebf1^neg^ or 500 Ebf1^OE^ ECs in medium containing BMS-754807 (IGF signaling inhibitor), AMD3465 (CXCL12 pathway blocker), or LY411575 (NOTCH antagonist). Endothelial tube formation was monitored at 0, 4, and 6 hours by phase-contrast imaging. **b,** Tube formation assay cocultures of 9,500 Ebf1^neg^ ECs with 500 Ebf1^neg^ ECs. **c,** Tube formation assay cocultures of 9,500 Ebf1^neg^ ECs with 500 Ebf1^OE^ ECs. Scale bars, 150 μm. Data are presented as mean ± s.e.m. with individual data points shown. Statistical comparisons were performed using two-sided Mann-Whitney tests; p values are indicated on the graphs.

**Extended Data Fig. 14 gCap-derived EBF1-expressing ECs contribute to PAH vascular lesions. a**, Experimental schematic for lineage labeling of *Aplnr*-expressing gCap cells in healthy adult *Aplnr*-CreERT2;*Rosa26^tdTomato^* (gCap reporter) mice in **b, c**. **b**, Representative whole-mount image from healthy adult gCap reporter mice. *Aplnr*-lineage (*Aplnr*^lin^) ECs were identified by tdTomato and ERG expression, and EBF1 expression was detected by smFISH or immunofluorescence. n=5. **c**, Quantification of *Ebf1* expression within *Aplnr-*lineage ECs (*Ebf1^+^*tdTomato^+^ERG^+^ cells / tdTomato^+^ERG^+^ cells × 100) in control lungs and after 1, 2 or 3 weeks of SuHx. **d**, Experimental schematic for lineage tracing *Ebf1*-expressing cells during SU5416 plus hypoxia (SuHx)-induced pulmonary vascular remodeling using *Ebf1*-CreERT2;*Rosa26^tdTomato^* (*Ebf1*-reporter) mice in **e, f**. **e**, Representative lung sections from *Ebf1*-reporter mice after 2 or 3 weeks of SuHx, showing *Ebf1*^lin^ ECs (tdTomato^+^ERG^+^*Pecam^+^*) within remodeled PAs and neointimal lesions. n=6. **f**, Quantification of *Aplnr^+^Ebf1*^lin^ ECs among all *Ebf1* lineage-labeled cells (left, *Aplnr^+^* tdTomato^+^ERG^+^ cells/ tdTomato^+^ERG^+^ cells x 100) and *Ebf1*^lin^ ECs among all ECs (right, tdTomato^+^ERG^+^ cells/ ERG^+^ cells x 100) in control lungs and during SuHx-induced remodeling. **g**, Experimental schematic for lineage tracing of *Aplnr*^+^ gCap cells during house-dust-mite antigen (HDM)-induced pulmonary vascular remodeling using the *Aplnr*-CreERT2;*Rosa26^tdTomato^* (gCap-reporter) mice in **h**, **i**. **h,** Representative lung-section images from gCap-reporter mice after 3 or 5 weeks of HDM exposure. *Aplnr*^lin^ ECs acquiring *Ebf1* expression (tdTomato^+^ERG^+^ *Ebf1^+^Pecam^+^*) were detected within remodeled PAs and neointimal lesions. n=6. **i**, Quantification of *Aplnr*-lineage contribution to the remodeled endothelium (left, tdTomato^+^ERG^+^ cells/ ERG^+^ cells x 100) and *Aplnr*-lineage coverage of *Ebf1*^+^ ECs (right, *Ebf1^+^*tdTomato^+^ERG^+^ cells/ *Ebf1^+^*ERG^+^ cells x 100) in control lungs and after 3 or 5 weeks of HDM exposure. **j**, Experimental schematic for lineage tracing of *Ebf1*-expressing cells during HDM-induced pulmonary vascular remodeling using *Ebf1*-reporter mice in **k**-**m**. **k, l**, Representative lung sections from *Ebf1* reporter mice after 3 or 5 weeks of HDM exposure, showing accumulation of *Ebf1^l^*^in^ ECs within remodeled PAs (**k**) and their co-expression of *Aplnr* (tdTomato^+^ERG^+^*Aplnr^+^*, **l**). n=6. **m**, Quantification of *Ebf1*-lineage contribution to the remodeled endothelium (left, tdTomato^+^ERG^+^ cells/ ERG^+^ cells x 100) and *Ebf1*-lineage coverage of *Ebf1*^+^ ECs (right, *Ebf1^+^*tdTomato^+^ERG^+^ cells/ *Ebf1^+^*ERG^+^ cells x 100) in control lungs and after 3 or 5 weeks of HDM exposure. n=6. **n**, Experimental schematic for lineage tracing of endothelial cells during SuHx-induced pulmonary vascular remodeling using EC-reporter mice (*Cdh5*-CreERT2;*Rosa26^tdTomato^*) in **o**. **o**, Representative lung sections from EC-reporter mice after 2 or 3 weeks of SuHx exposure, showing *Cdh5*-lineage labeling of ERG ECs within remodeled PAs and neointimal lesions, including *Ebf1 ERG* lesion ECs. n=5. **p**, Quantification of *Cdh5*-lineage contribution to the remodeled endothelium (left, tdTomato^+^ERG^+^ cells/ ERG^+^ cells x 100) and *Cdh5*-lineage coverage of *Ebf1*^+^ ECs (right, *Ebf1^+^*tdTomato^+^ERG^+^/*Ebf1^+^*ERG^+^) during SuHx-induced remodeling. **q, r**, Representative whole-mount images from *Ebf1*-reporter mice after 3 weeks of SuHx (**q**) or 5 weeks of HDM (**r**), showing the spatial relationship between tdTomato-labeled *Ebf1*^lin^ ECs and Ki67^+^ERG^+^ proliferating ECs within remodeling regions. Boxed regions and corresponding single-channel images are shown at higher magnification; whole-mount images include 3D surface rendering. n=5. **s**, Quantification of proliferative activity within *Ebf1*-lineage (tdTomato Ki67 ERG cells/ tdTomato ERG cells ×100) and non-*Ebf1*-lineage (tdTomato Ki67 ERG cells/ tdTomato ERG cells ×100) ECs during SuHx-induced remodeling. For each mouse, 3-5 remodeled vessels or vascular regions were analyzed. For **c**, **f**, **i**, **m**, **p**, and **s**, ROI results were averaged to generate one biological replicate value; n=5-7 mice per group. Data are presented as mean ± s.e.m. with individual data points shown. Statistical comparisons in **c**, **f**, **i**, **m**, and **p** were performed using Kruskal-Wallis tests followed by Dunn’s multiple-comparisons tests; comparisons in **s** were performed using Wilcoxon matched-pairs test; p values are indicated on the graphs. Scale bars, 20 μm for **b, e, h, k, l** and **o** and 25 μm for **q** and **r**.

**Extended Data Fig. 15 gCap-specific *Ebf1* deletion and endothelial *Ebf1* silencing protect against PAH. a**, Experimental schematic for genetic deletion of *Ebf1* in adult *Aplnr*^+^ gCap cells during SuHx-induced PAH. Tamoxifen was administered to *Aplnr*-CreERT2;*Rosa26^tdTomato^*control mice (gCap-Ctr) and *Aplnr*-CreERT2;*Rosa26^tdTomato^;Ebf1^fl/fl^* mice (gCap-Ebf1 KO) before initiation of SuHx. Mice received SU5416 (20 mg/kg, s.c., weekly) and were maintained under hypoxia conditions for three weeks. **b**, Representative smFISH combined with immunofluorescence from gCap-Ctr and gCap-Ebf1 KO lungs after three weeks of SuHx. Dotted outlines and boxed regions mark *Ebf1*^+^*Aplnr*^lin^ ECs (tdTomato^+^ERG^+^ *Ebf1^+^Pecam^+^*) within remodeled PAs. n=7 mice per group. **c,** Experimental schematic for endothelial-targeted *Ebf1* knockdown in the *Bmpr2**^+/-^*** rat PAH model. *Bmpr2**^+/-^*** rats were treated with an endothelial-tropic AAV carrying a tdTomato reporter and either three independent *Ebf1*-targeting shRNAs (PAH+*shEbf1*) or matched scrambled shRNAs (PAH+*scramble* group) two weeks before PAH induction. *Bmpr2**^+/-^*** animals without PAH induction served as negative controls (Ctr). *Bmpr2**^+/-^*** rats without AAV treatment but with PAH induction were used as positive controls (PAH). **d**, Representative immunofluorescence images from PAH+*scramble* group at 1 week and 3 weeks after PAH induction. Dotted outlines mark AAV reporter-labeled (red)ERG^+^ ECs detected in the capillary bed (at 1 week) and within PA (at 3 weeks). Quantification of AAV-BI30 endothelial transduction efficiency was calculated as the % ERG ECs that were AAV-tdTomato (tdTomato ERG /total ERG cells × 100). Three vascular volumes (100 × 100 × 50 μm) were analyzed per rat by three-dimensional Imaris surface analysis and averaged to generate one biological replicate. n = 5 rats; data are presented as mean ± s.e.m. with individual animals shown. **e**, Hemodynamic measurements. n=7 rats per group. **f,** Representative H&E images from the indicated treatment groups. n=7. Dotted outlines mark the vessel walls. **g,** Representative immunofluorescence images of rat lungs collected at 3 weeks. n=7. **h,** Representative smFISH combined with immunofluorescence images of rat lungs collected 3 weeks following PAH induction. n=7. Yellow dotted outlines mark *Ebf1^+^Pecam^+^*ERG^+^ECs. **i,** Representative smFISH combined with immunofluorescence images of PAH+*shEbf1* rats showing *Dll4^+^Pecam^+^*ERG^+^ECs (yellow dotted outlines). n=7. **j,** Representative immunofluorescence images from control, PAH+scramble, and PAH+*shEbf1* rats. Red arrowheads indicate PA branching points. n=7. **k**, Quantification of PA number and branches in the indicated groups (see Methods). n=7 rats per group. Data in **e** and **k** are presented as mean ± s.e.m. with individual data points shown; statistical comparisons were performed using Kruskal-Wallis tests followed by Dunn’s multiple-comparisons tests; p values are indicated on the graphs. Scale bars, 20 μm in **b**, **h, i**, 25 μm in **d**, **g**, and 50 μm in **j**.

**Extended Data Fig. 16 Ectopic endothelial *Ebf1* expression causes vascular remodeling and PAH. a,** Experimental schematic for **b-d**. **b**, **c**, Representative immunofluorescence images of adult lung samples collected from *Rosa26*^tdTomato^;*H11*^LSL-Ebf1^ (Cre-negative control) or *Cdh5-* CreERT2;*Rosa26*^tdTomato^;*H11*^LSL-Ebf1^ (EC-Ebf1 OE) mice. Yellow dotted outlines mark PA branches. Yellow arrowheads indicate increased αSMA coverage. n=6 mice per group. **d**, Quantification of αSMA coverage in the distal PA. n=6 mice per group. For each mouse, 5 ROIs (PA<50µm in size) were analyzed and averaged to generate a single value. **e,** Experimental schematic of **f-k**. **f,** Representative immunofluorescence images of adult lung samples collected from *Aplnr-*CreERT2; *Rosa26*^tdTomato^ (gCap control) or *Aplnr-*CreERT2;*Rosa26*^tdTomato^; *H11*^LSL-Ebf1^ (gCap-Ebf1 OE) mice after 3 weeks of treatment of SU5416. Red arrowheads indicate *Aplnr-* CreER lineage-labeled ECs (tdTomato^+^, white) co-stained with ERG in the remodeled PA. n=6. **g, h**, Representative smFISH combined with immunofluorescence images of d21 lung sections from SU5416-treated gCap-Ctr and gCap-*Ebf1* OE mice showing *Cxcl12* RNA. n=6 mice per group. **i**, Quantification of *Cxcl12^+^* ECs (% of ERG^+^ nuclei with *Cxcl12* puncta) within ROIs. n=6 mice per group. For each lung, 5 vascular ROIs (PA<50μm) were analyzed and averaged to generate a single value. **j,** Representative immunofluorescence images of adult lung samples collected from *Aplnr-*CreERT2;*Rosa26*^tdTomato^ (gCap-Ctr, n=14) or *Aplnr-* CreERT2;*Rosa26*^tdTomato^; *H11*^LSL-Ebf1^ (gCap-Ebf1 OE, n=14) mice. Red arrowheads indicate PA branching points. **k**, Quantification of PA branch density, PA density, and branch points per PA in gCap-Ctr and gCap-Ebf1 OE lungs (see Methods). n=10 mice in the gCap-Ctr group and n=14 mice for gCap-Ebf1 OE group. **l,** Proposed model of the *Ebf1*^+^ EC-driven, artery-forming niche in PA development. During lung morphogenesis, *Aplnr^+^* plexus progenitors give rise to both pre-arterial ECs and an *Ebf1*^+^ endothelial niche population. Without directly incorporating into the PA endothelium, these *Ebf1*^+^ ECs serve as the “PA organizing cells” by secreting vasculotrophic signals required for the proliferation and arterialization of the plexus progenitors, as well as for the recruitment and muscularization of the stromal progenitors. **m**, Working model of reactivation of the *Ebf1*^+^ EC-driven, artery-forming niche in adult PAH. During vascular remodeling, the rare pre-existing *Ebf1*-lineage ECs within the *Aplnr^+^* gCap compartment accumulate within remodeled vessels, together with an additional population arising through *de novo Ebf1* induction from the broader *Aplnr^+^* gCap pool. Through comparable vasculotrophic signaling, these *Ebf1*^+^ ECs induce the proliferation and arterialization of capillary stem cells, as well as the recruitment and activation of cells from the stromal layers, thereby contributing to PA vascular remodeling. Data in **d, i** and **k** are presented as mean ± s.e.m. with individual data points shown; statistical comparisons were performed using two-sided Mann-Whitney tests; p values are indicated on the graphs. Scale bars, 50 μm in **b**, **c**, and **j** and 20 μm in **f**, **g**, and **h**.

## Supplementary Information

**Supplementary Fig. 1 Generation of the *Ebf1^CreERT2^*mice. a,** Schematic of the generation of a mouse model with *Ebf1-P2A-CreERT2* knock-in allele using a CRISPR/Cas9-mediated strategy. Cas9 creates double-strand breaks (DSB) at the location specifically targeted by two gRNAs (green arrows). DSBs are then repaired by homology-directed repair (HDR) using a donor template. Two pairs of screening primers (F1a/seq677 and seq557/R2a) were designed to screen founders. **b,** Genotyping primers. **c,** Repair donor sequence (2,478 bp). Underlined sequences are Ebf1 genomic sequences used as homology arms.

**Supplementary Fig. 2 Generation of the Ebf1-overexpression mice. a,** Schematic of the generation of a mouse model with *H11-EF1a-loxP-STOP-loxP-EBF1-P2A-H2B-mTagBFP2* knock-in allele using a CRISPR/Cas9-mediated strategy. Cas9 creates double-strand breaks (DSB) at the location specifically targeted by two gRNAs (green arrows). DSBs are then repaired by homology-directed repair (HDR) using a donor template. Two pairs of screening primers were designed to screen founders. **b,** Genotyping primers. **c,** Repair donor sequence (5,904 bp). Sequences in gray are *Ebf1* genomic sequences used as homology arms.

**Supplementary Fig. 3| Generation of the endothelial-targeted *Ebf1* knockdown system in rats.** To achieve endothelial-targeted knockdown of *Ebf1* in a rat model, we generated an adeno-associated virus packaged with the endothelial-tropic AAV-BI30 capsid (an engineered AAV9 variant reported to efficiently transduce endothelial cells)^95^. This platform additionally incorporates a two-vector PiggyBac system to enable stable genomic integration and durable lineage labeling *in vivo*: one vector encodes a hyperactive, nuclear-localized PiggyBac transposase, and the companion vector carries the PiggyBac-flanked (transposon) cargo comprising the shRNA cassette (and a tdTomato reporter). The knockdown cassette contains three independent miR30-based shRNA hairpins targeting *Ebf1* to enhance robustness; matched control vector encodes non-targeting scramble hairpins. **a**, Vector map of the shEbf1 knockdown cassette (7,184 bp). **b**, Vector map of the scramble control cassette (7,184 bp). **c**, Vector map of the PiggyBac transposase cassette (7,249 bp).

## Acknowledgements

We thank A. Gillich for methodological guidance on *Aplnr*-CreERT2 lineage labeling of the embryonic pulmonary vascular plexus. Human lung tissue and single-cell data were provided by the Pulmonary Hypertension Breakthrough Initiative (PHBI). This work was supported by I01 BX005628, R01HL158714, R01 HL150583, R01 HL138473. M.A.K. and K.R.-H. are investigators of the Howard Hughes Medical Institute.

## Author contributions

W.T., S.G. and S.L. performed all *in vivo* experiments. W.T. and S.L. performed all *in vitro* experiments. T.H.W. performed all bioinformatic analyses. A.A. performed immunostaining in the HDM model. W.T., S.G., S.L., J.C., R.V. and C.H. performed embryo dissections. D.K. and Y.Z. performed FACS sorting. W.T., S.G., S.L., H.Z. and D.K. performed single-cell library preparation. W.T., S.G., R.V., J.C., C.H., K.S. and E.B. managed animal colonies. A.A., R.Z., K.Y., A.C., J.H., S.A., V.A.d.J.P., Z.D. and M.R. provided materials. W.T., T.H.W., J.P., P.K., T.D., R.J., L.P., L.B., X.J., M.R. and K.R.-H. designed experiments and interpreted the data. M.N. supervised the study. W.T. led experimental design and execution. T.H.W. led the analytical framework and statistical modeling. H.Z., D.Y. and Z.D. contributed human datasets and translational integration. P.E.B. and M.A.K. provided the foundation for the developmental studies. M.A.K. supported and mentored T.H.W. W.T., T.H.W. and M.N. conceived the study, interpreted the data and wrote the manuscript. All authors reviewed and edited the manuscript.

