## Extended Data Figures for "EBF1 controls an embryonic artery-forming niche that reactivates in pulmonary arterial hypertension"

### Extended Data Fig. 1 Single cell RNA-seq analyses of rat PAH vascular lesions

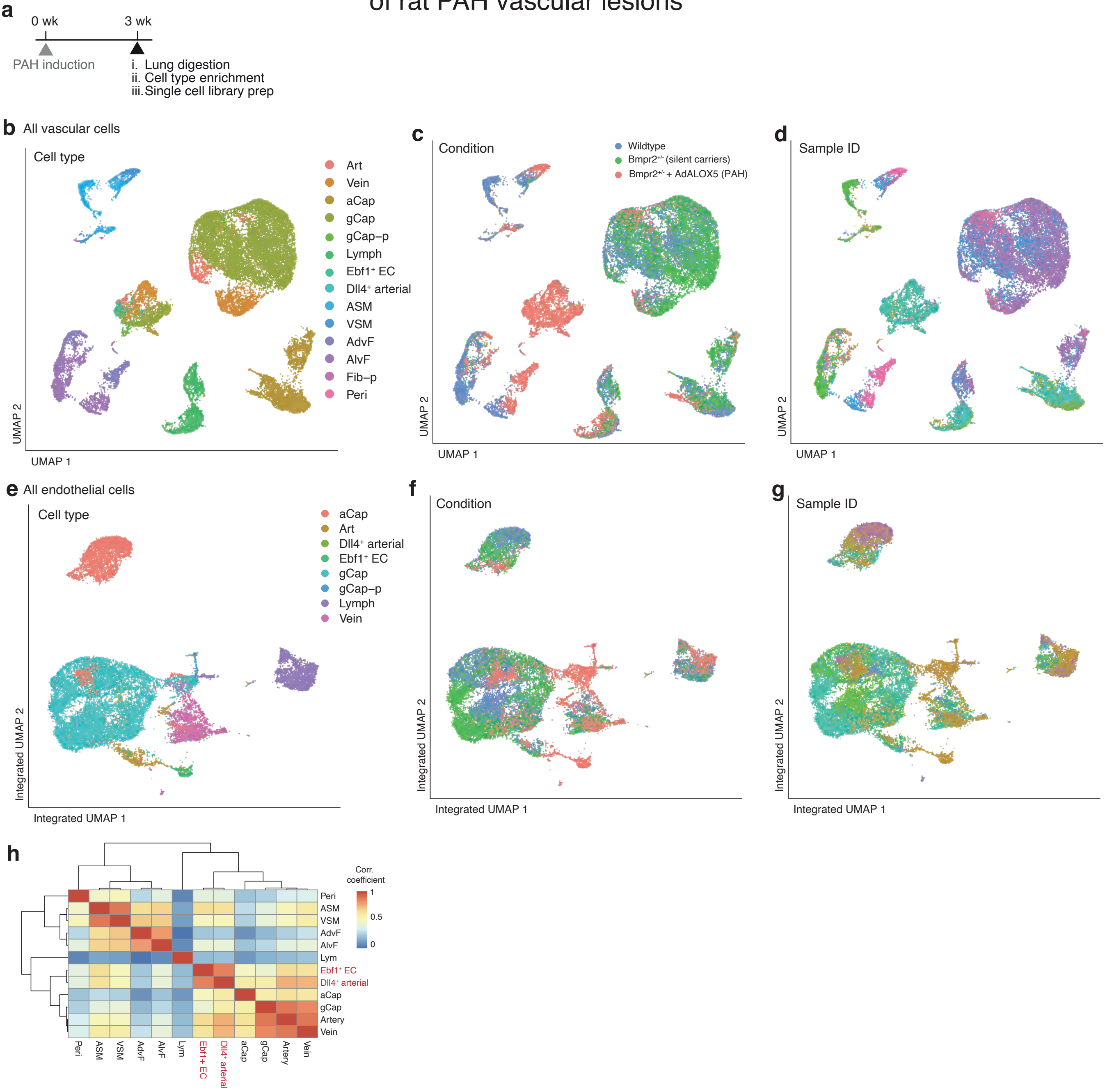

### Extended Data Fig. 2 EBF1⁺ ECs are broadly induced across animal models of PAH

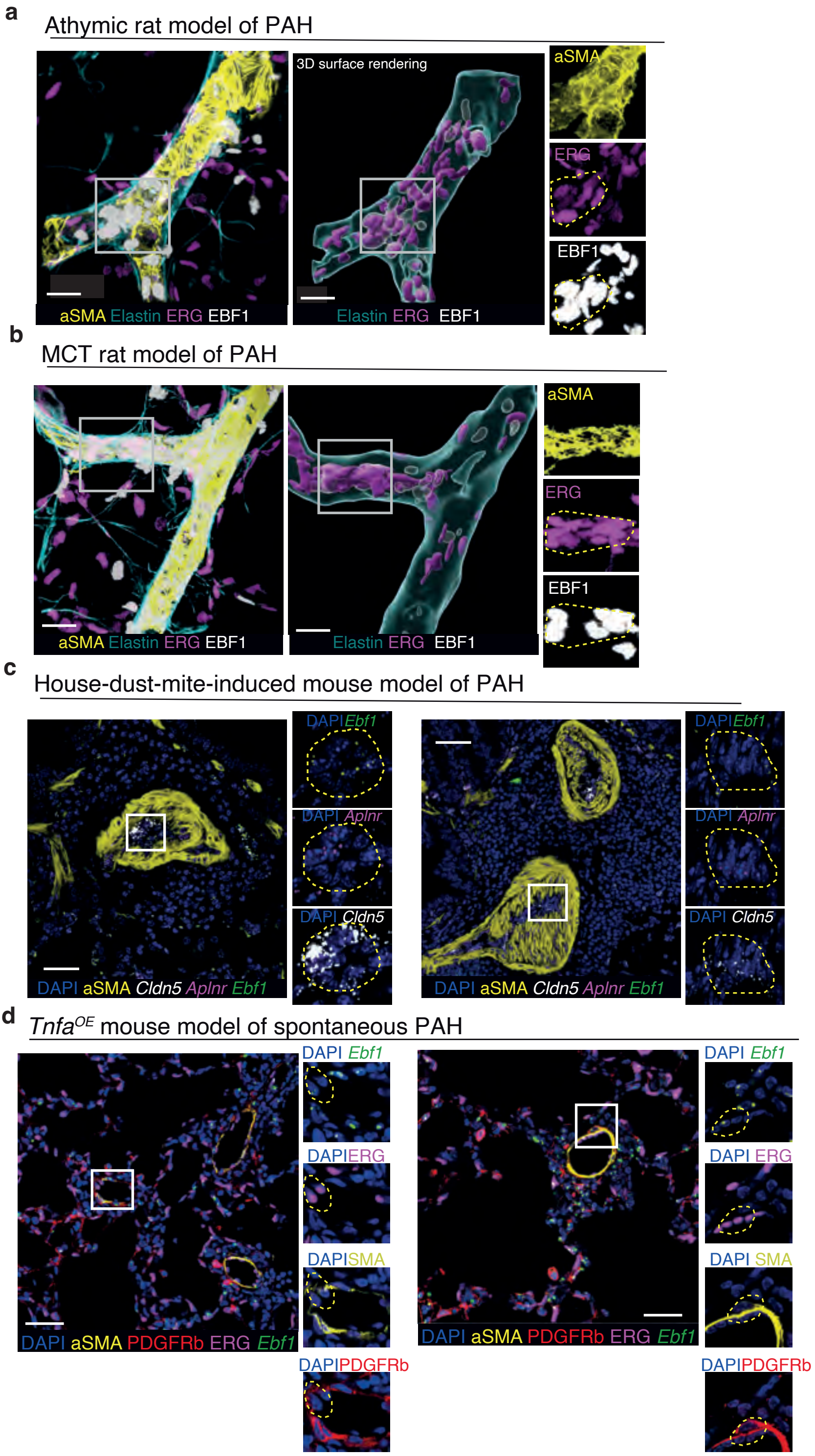

### Extended Data Fig. 3 EBF1⁺ ECs are a rich source of vasculotrophic signals associated with PAH pathogenesis

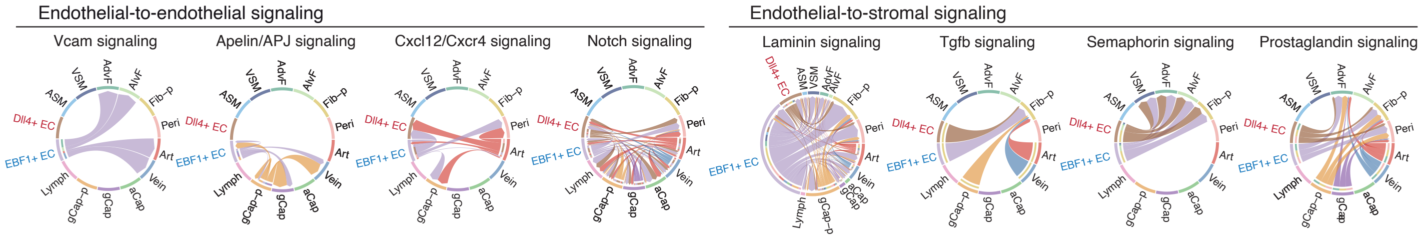

### Extended Data Fig. 4 Human PAH features disease-associated EBF1⁺ECs and CXCL12$^{high}$ neo-arterial ECs

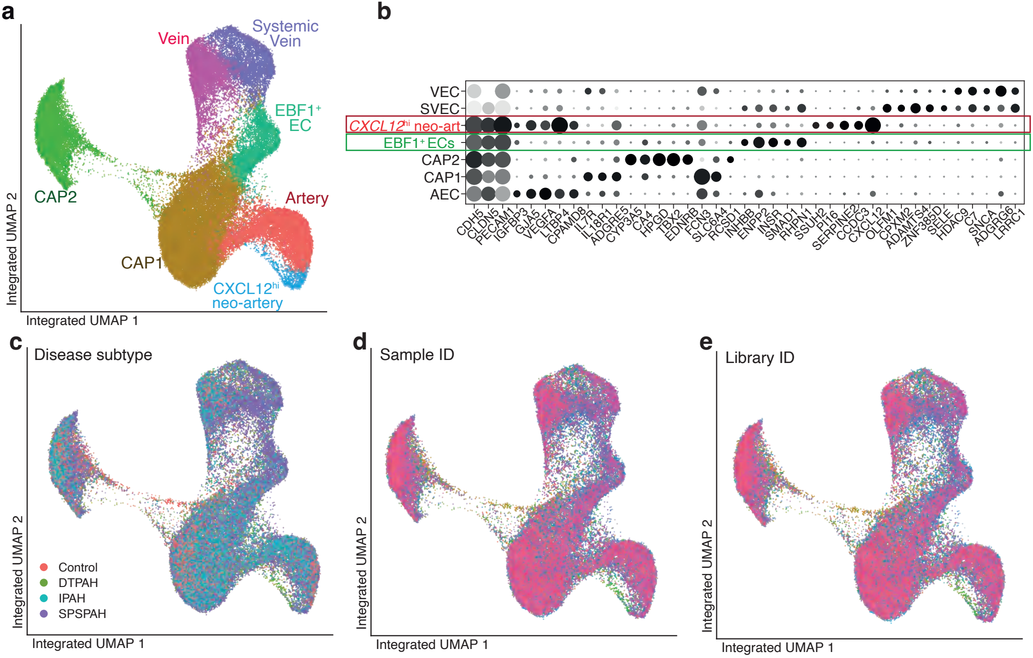

### Extended Data Fig. 5 PAH-associated signals regulate EBF1 expression in microvascular ECs

**a**

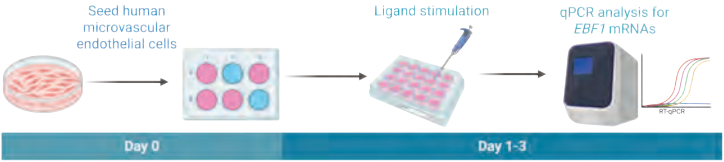


| Ligands | Concentrations |
|---|---|
| BMP9 | 5ng/ml |
| TGFb1 | 5ng/ml |
| ActivinA | 10ng/ml |
| GDF11 | 25ng/ml |
| IGF1 | 100ng/ml |
| VEGF-165 | 10ng/ml |
| CXCL12 | 20ng/ml |

**b**

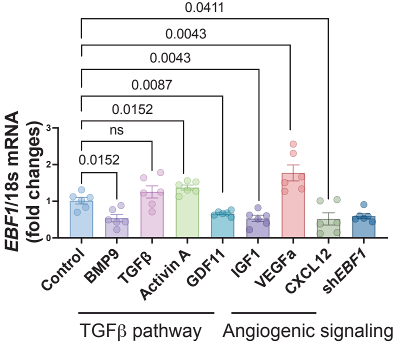

### Extended Data Fig. 6 scRNA-seq analyses of the developing mouse lung vasculature (E11.5-E15.5)

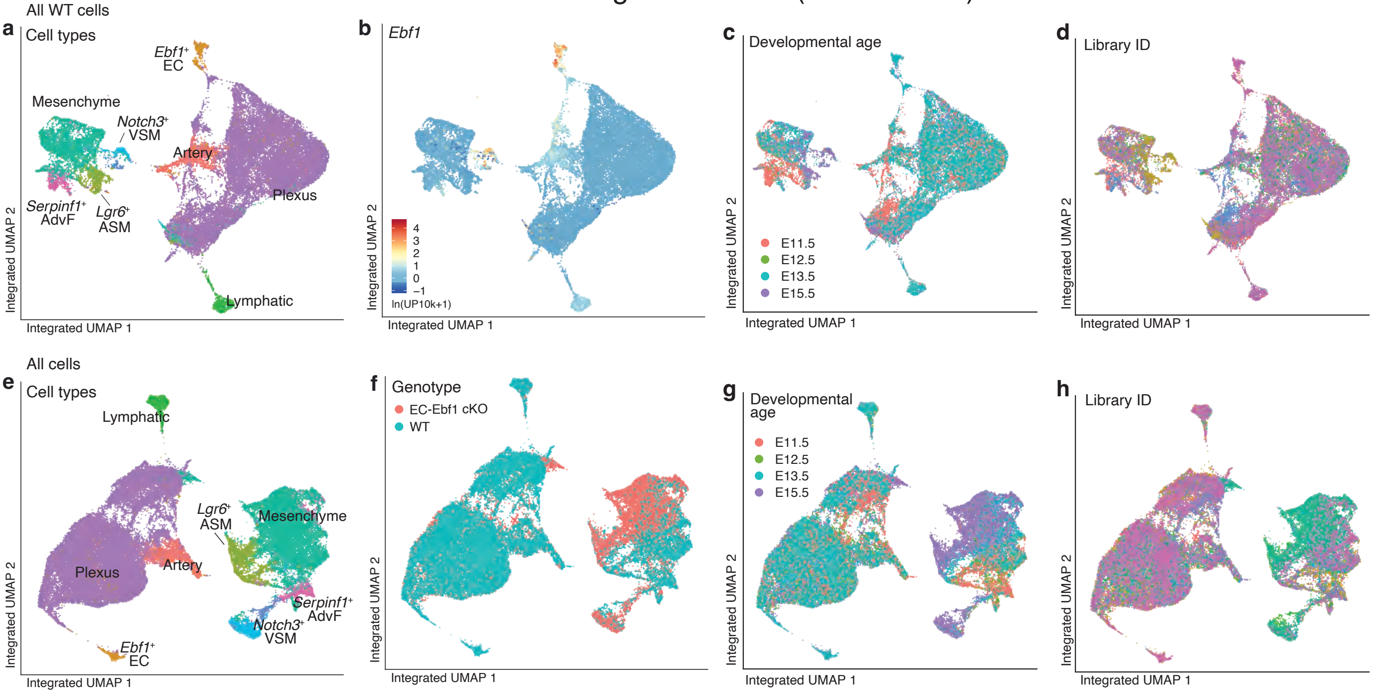

Extended Data Fig. 7 Pulmonary vasculature arises from *Aplnr*⁺ lung plexus progenitors

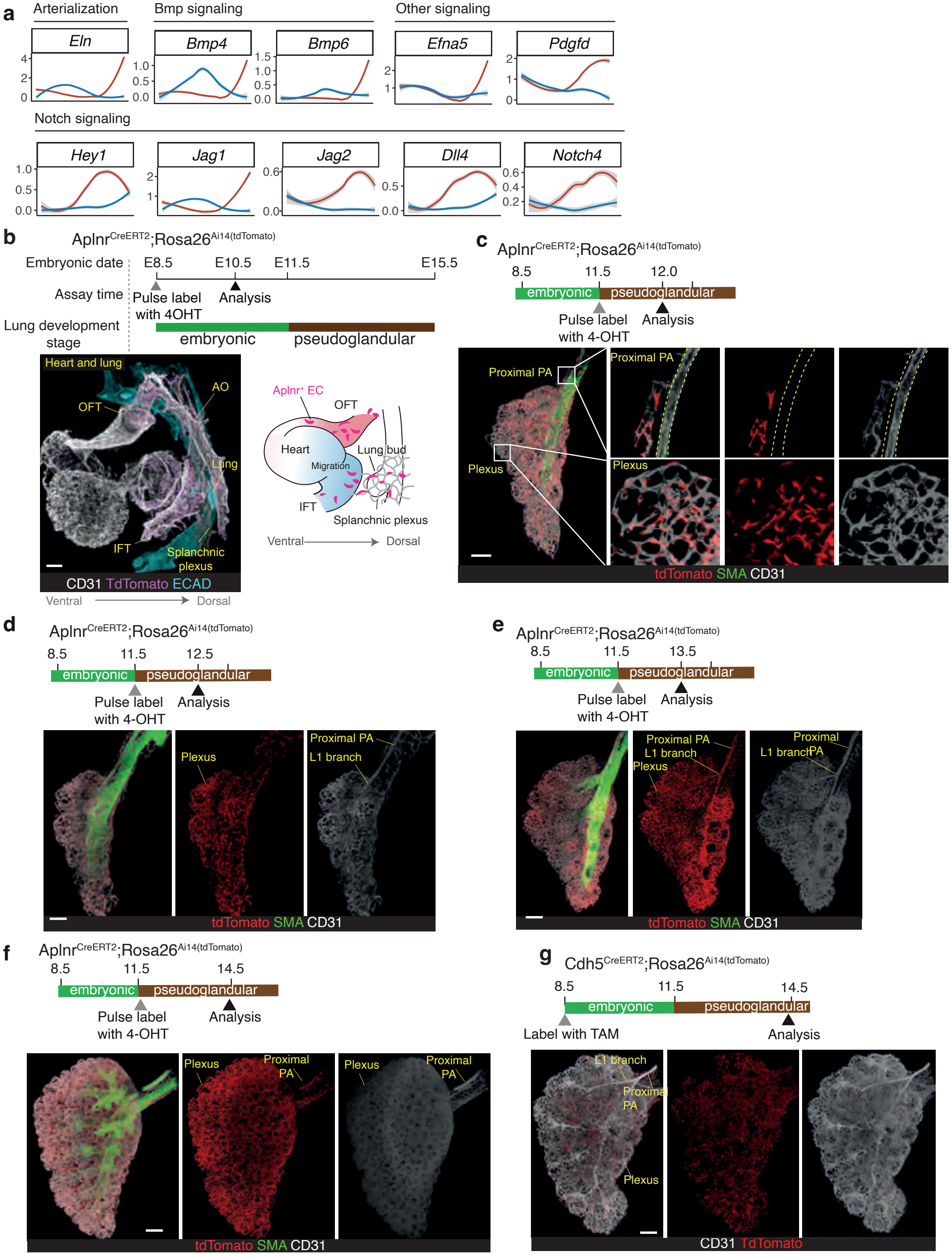

### Extended Data Fig. 8 EBF1⁺ ECs comprise a subset of Aplnr-lineage lung plexus cells

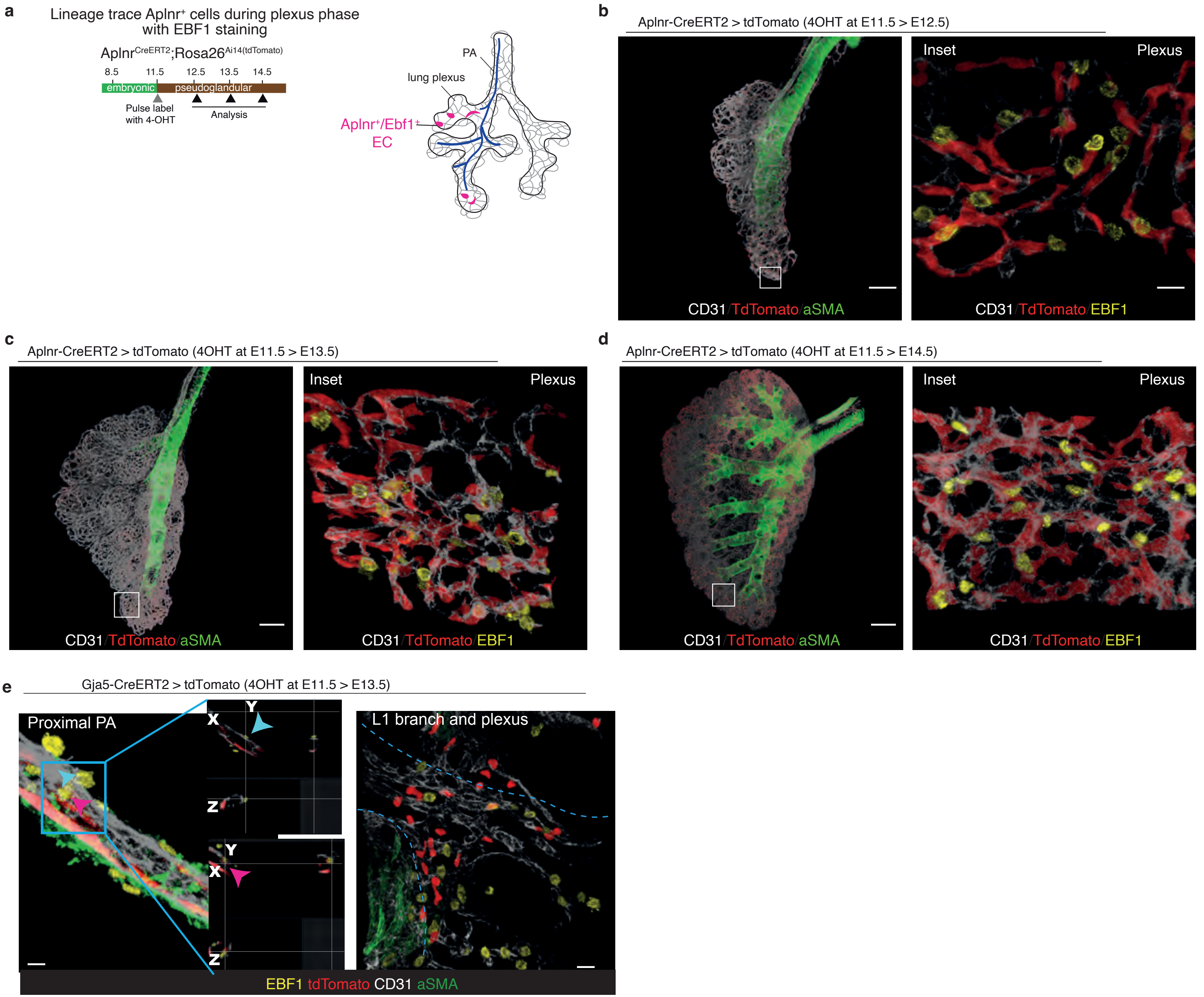

### Extended Data Fig. 9 Distribution of Ebf1$^{lin}$ cells in the embryonic stage of lung development

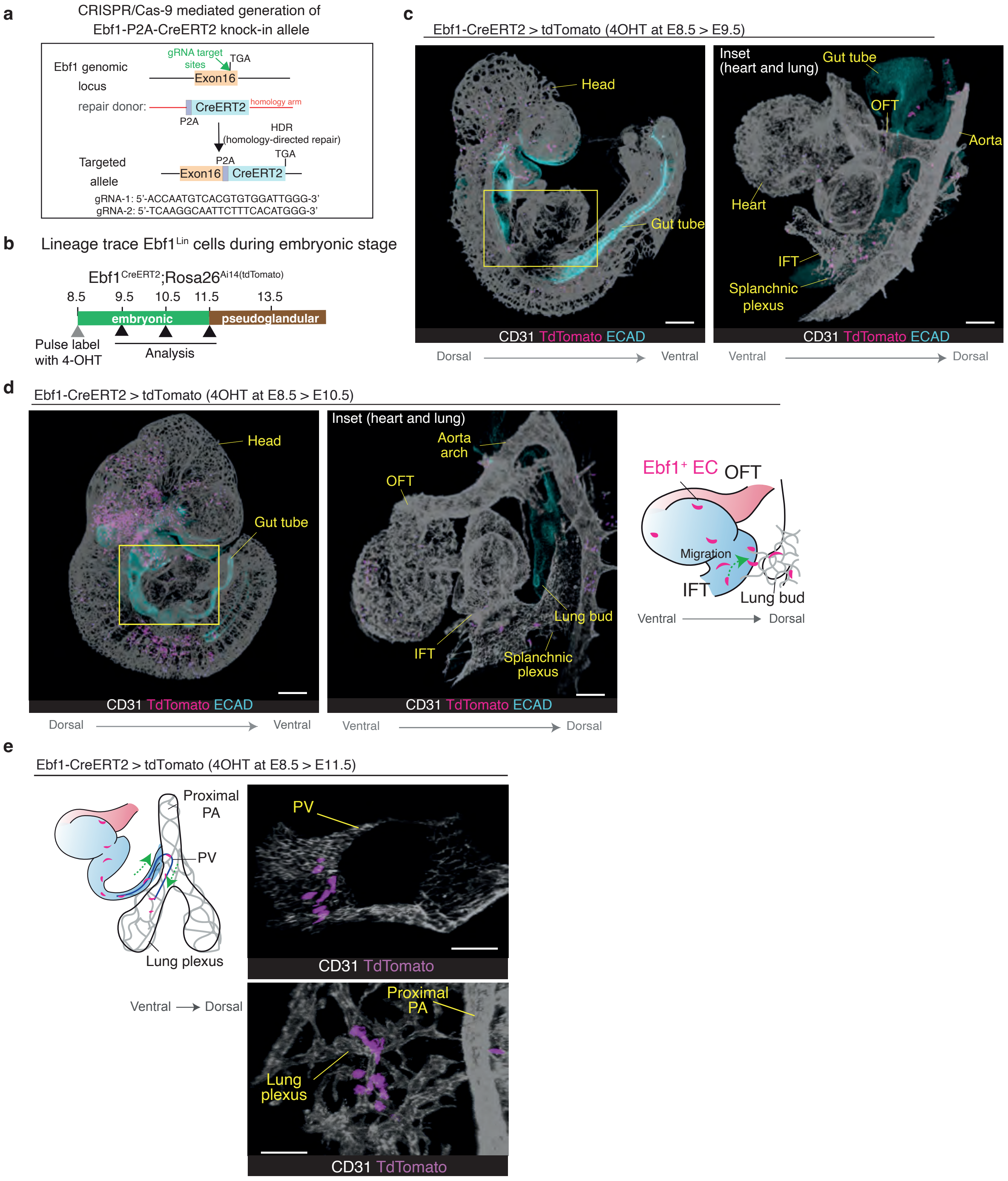

### Extended Data Fig. 10 *Ebf1*$^{lin}$ ECs do not directly trace into the developing PA endothelium

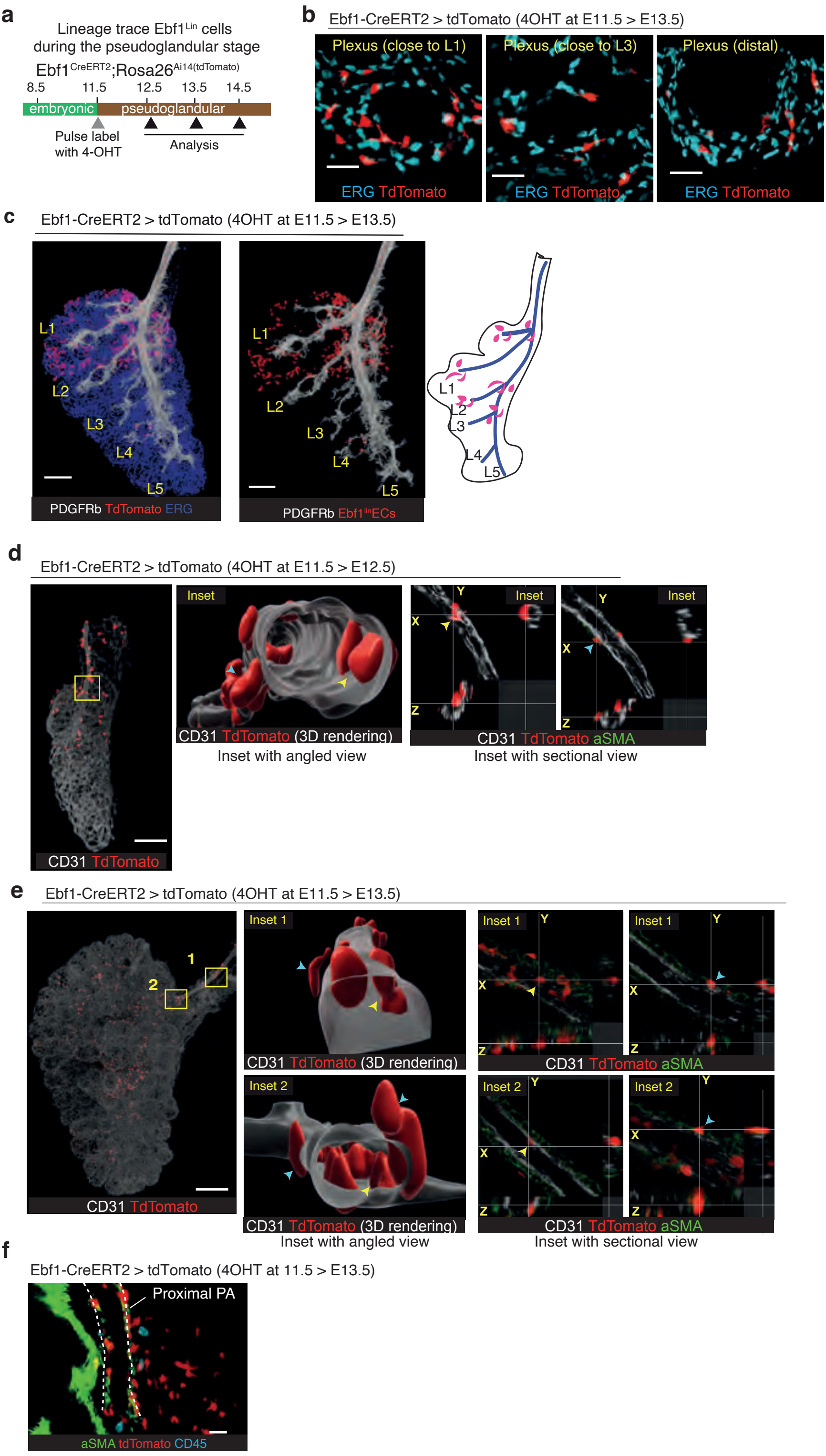

### Extended Data Fig. 11 *Ebf1^lin* ECs are transiently proliferative and migratory with rich vasculotrophic signals

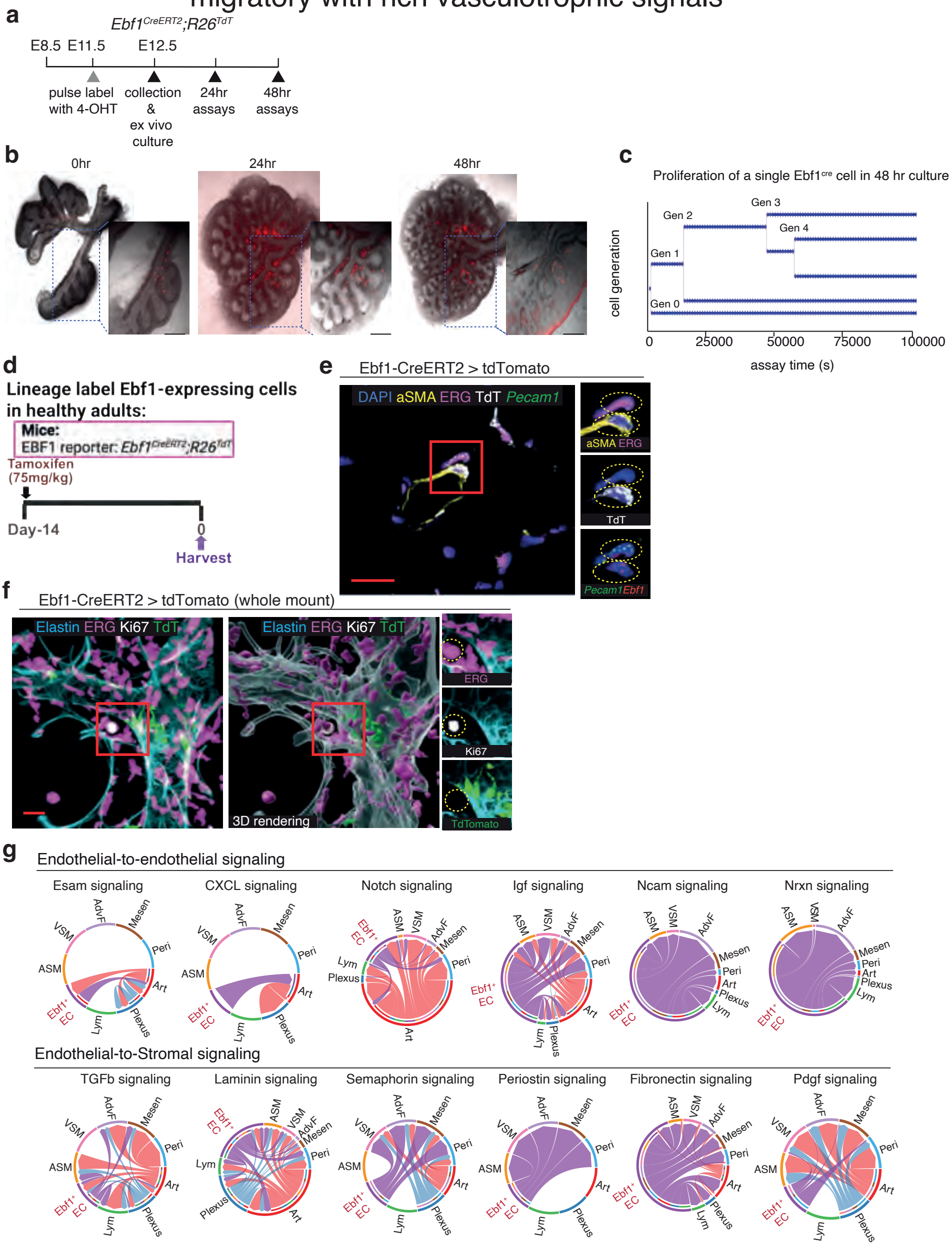

Extended Data Fig. 12 Plexus-specific *Ebf1* is essential for PA development

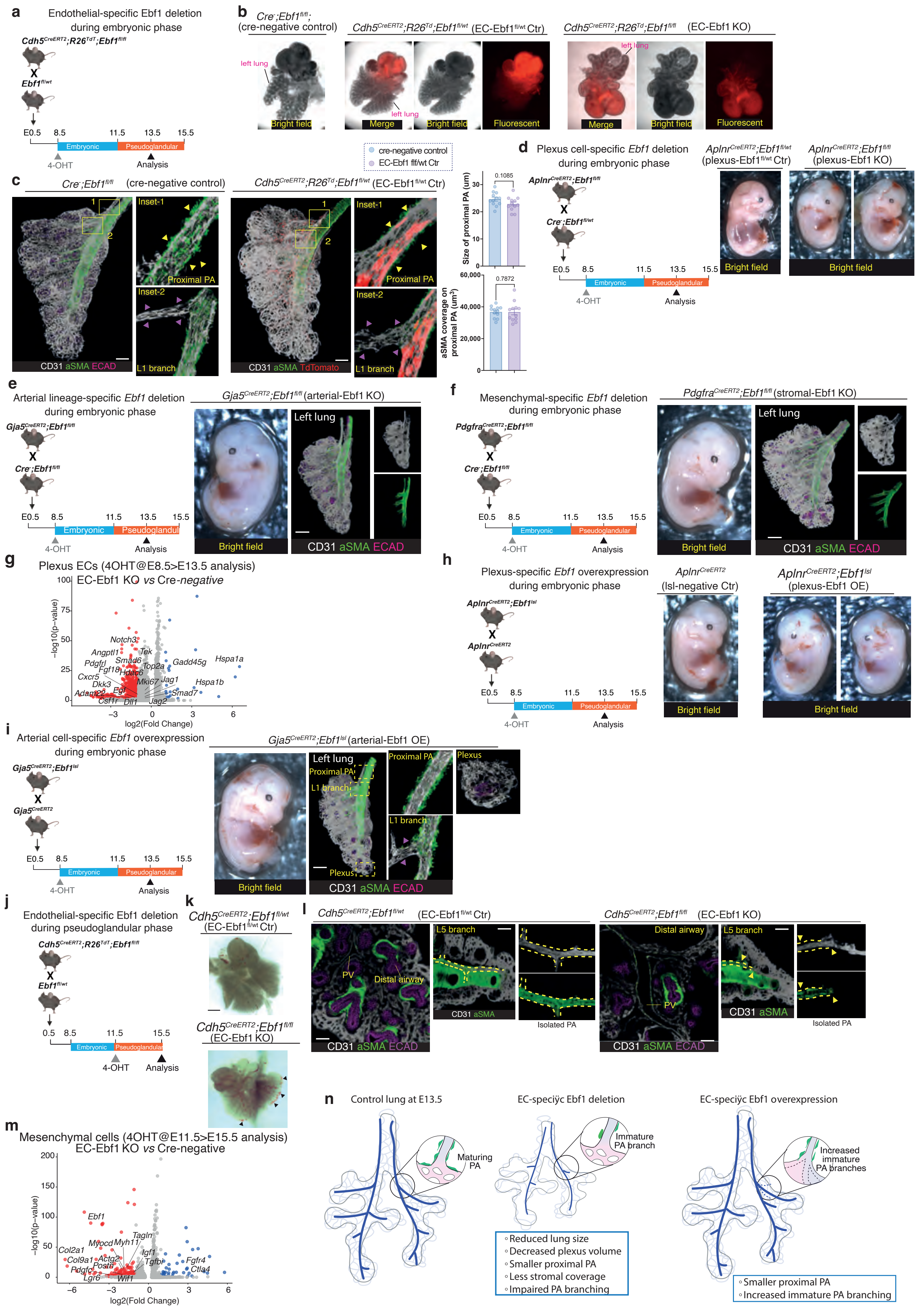

### Extended Data Fig. 13 EBF1⁺ECs promote angiogenesis of EBF1-negative cells

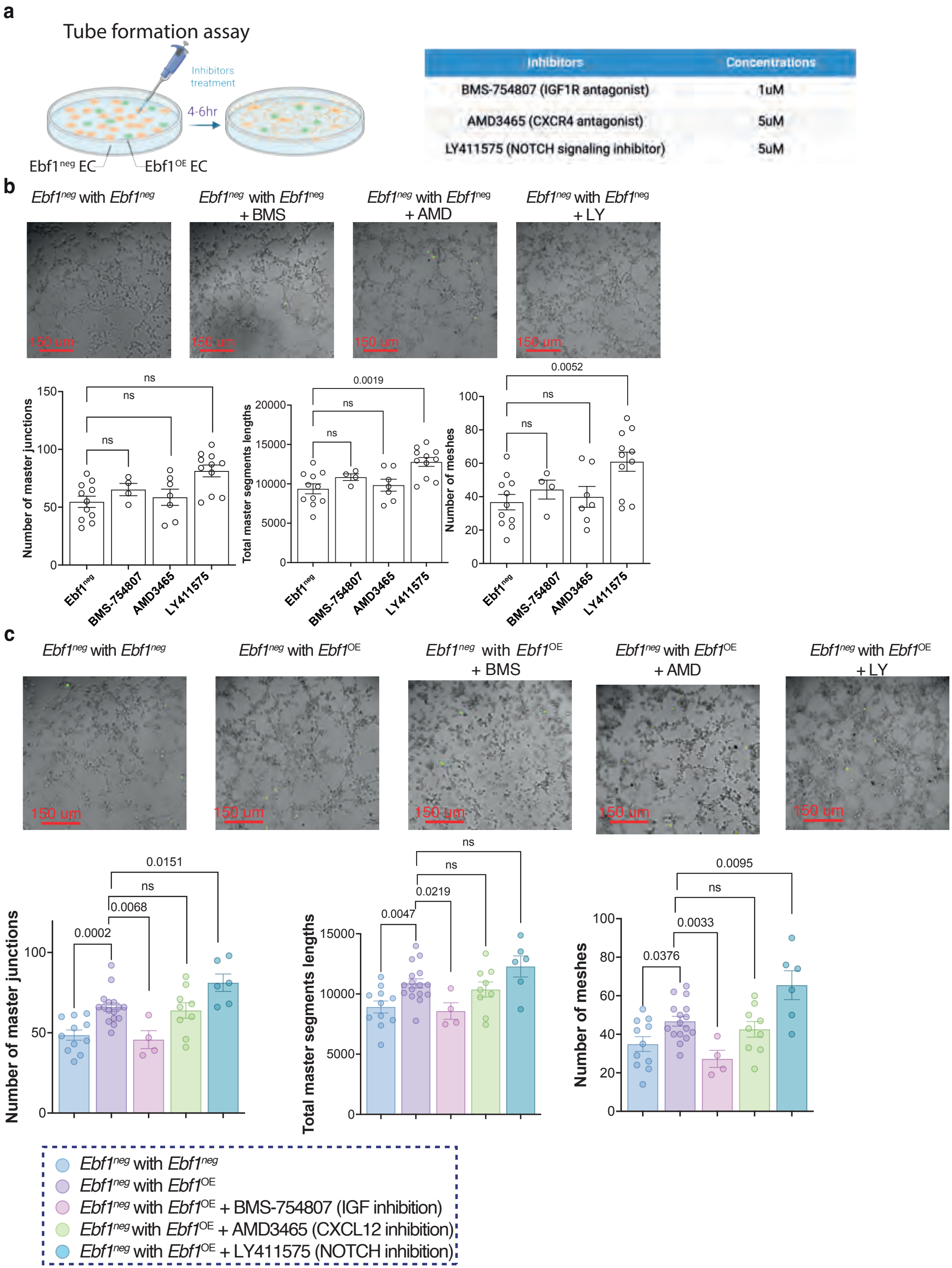

Extended Data Fig. 14 gCap-derived EBF1-expressing ECs contribute to PAH vascular lesions

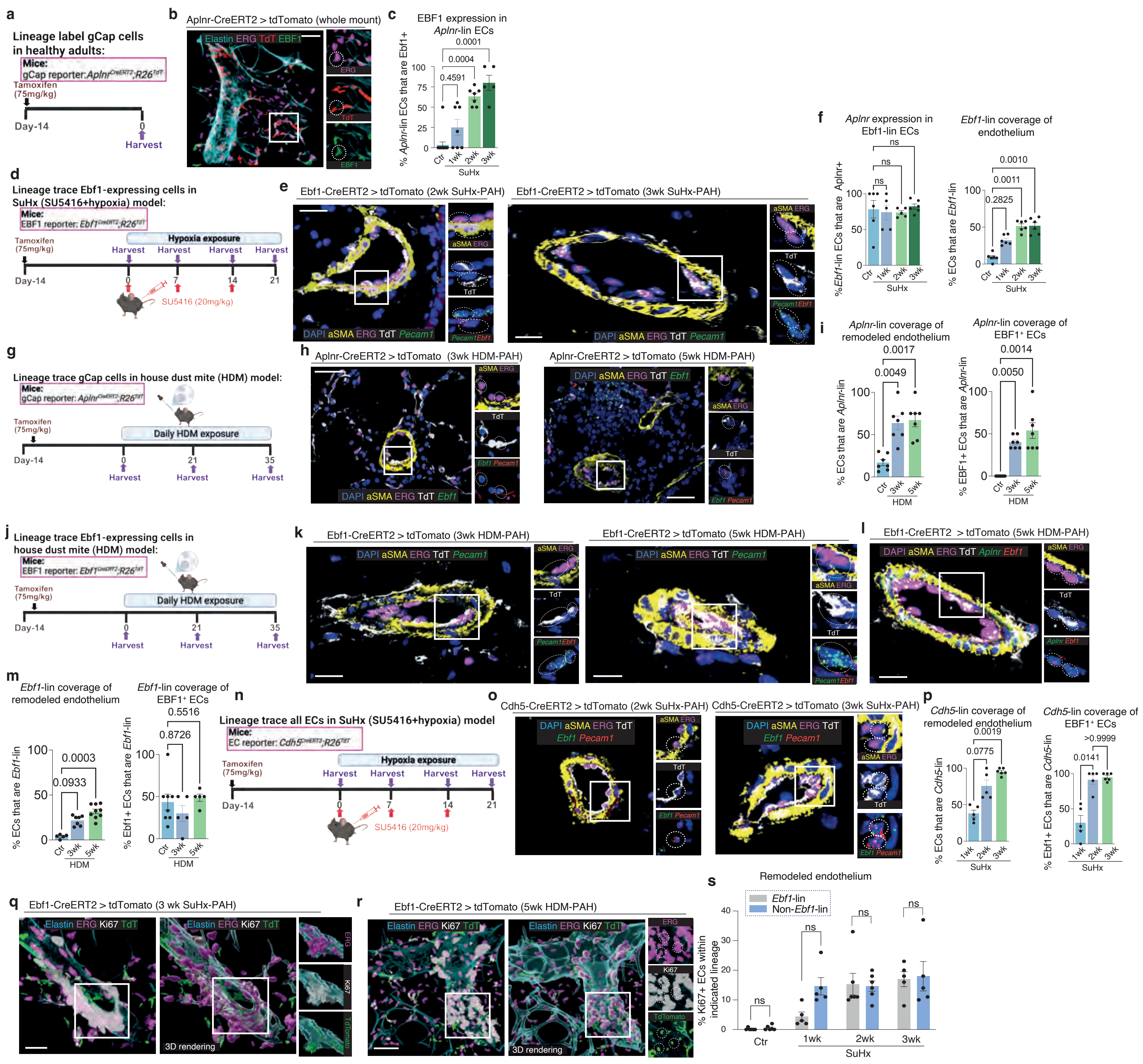

### Extended Data Fig. 15 gCap-specific *Ebf1* deletion and endothelial *Ebf*1 silencing protect against PAH

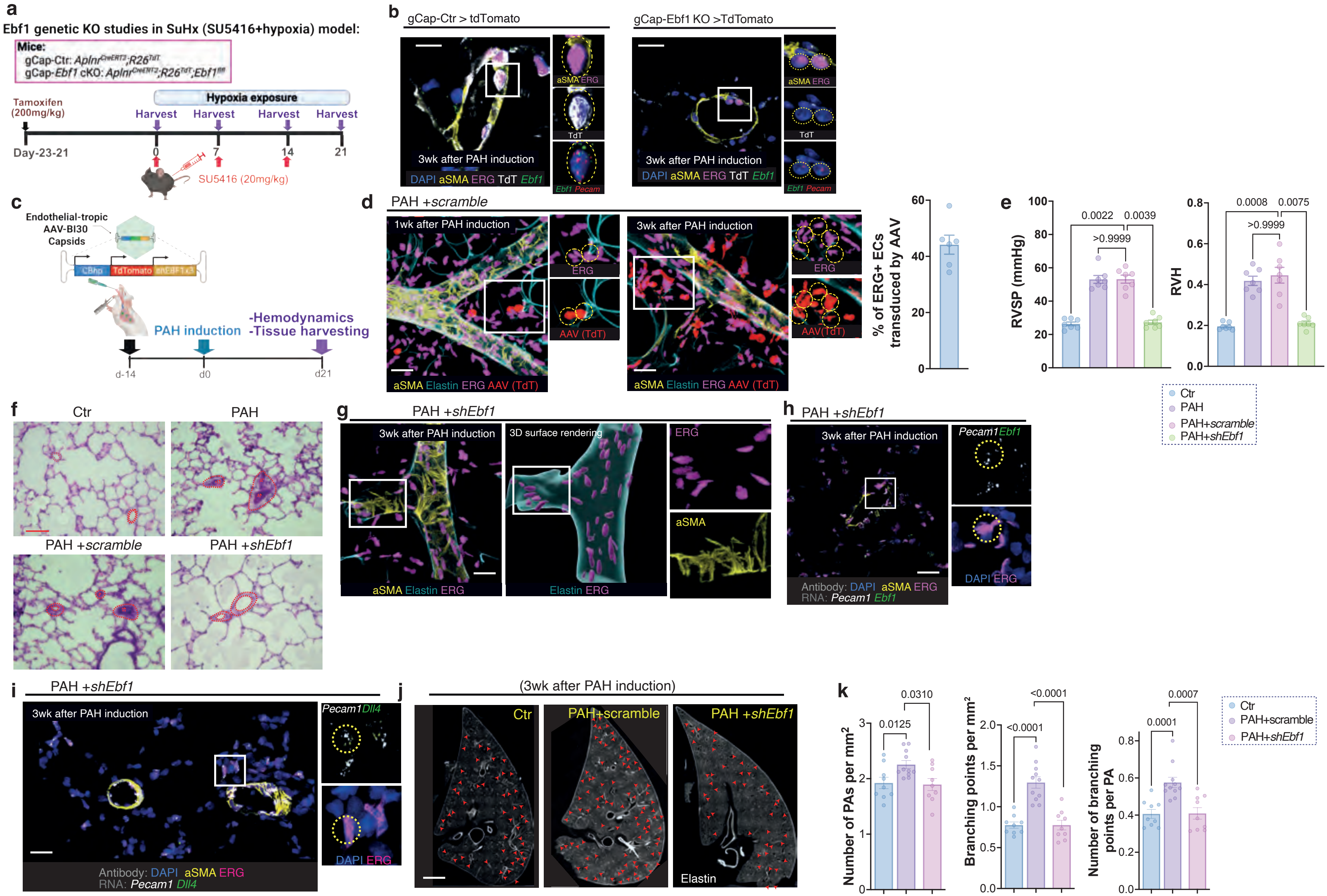

### Extended Data Fig. 16 Ectopic endothelial *Ebf1* expression causes vascular remodeling and PAH

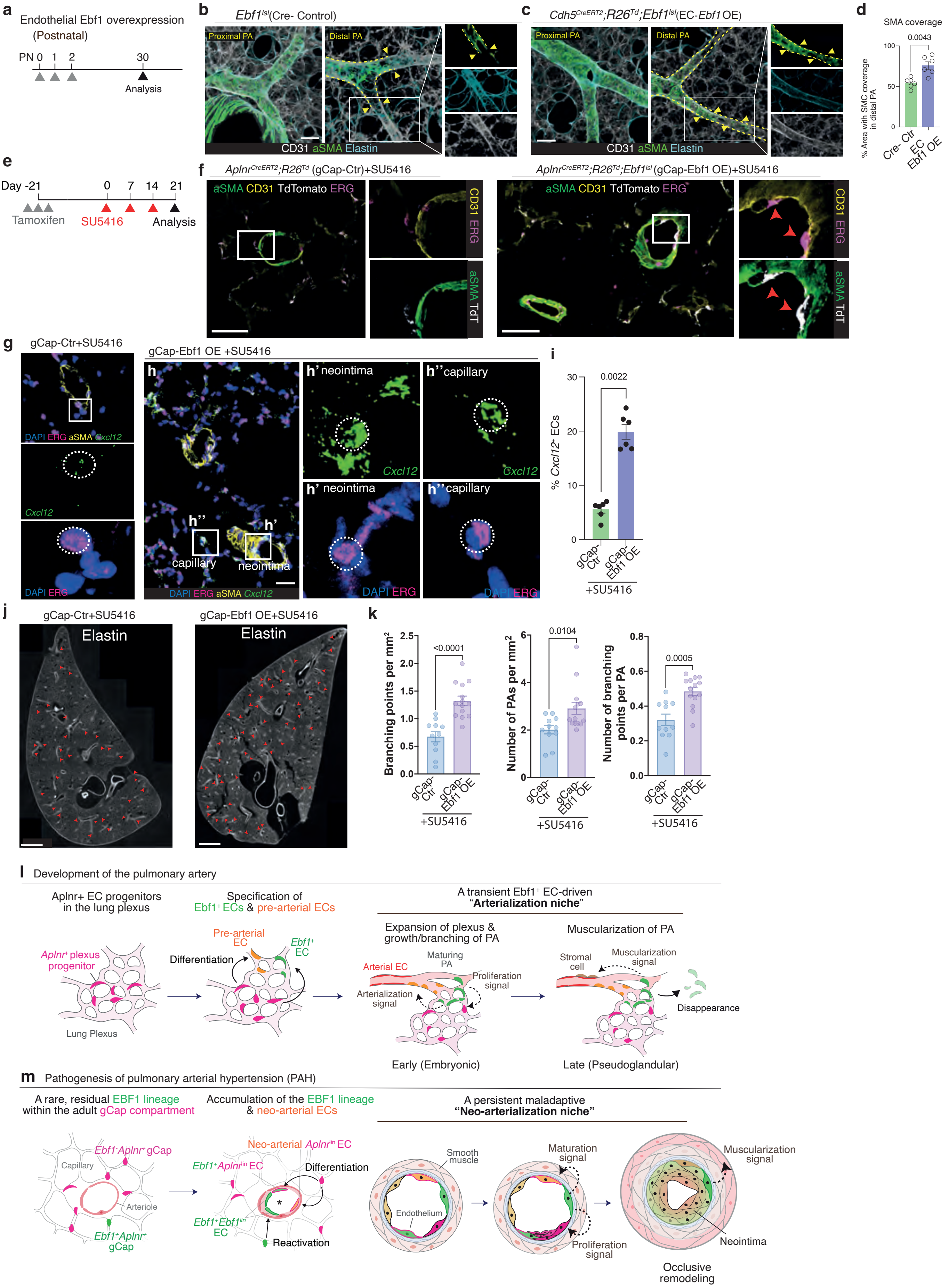
